# Multi-timescale PTM architecture in human RNA polymerase II

**DOI:** 10.64898/2026.06.27.734958

**Authors:** Ildefonso M. De la Fuente, José Carrasco-Pujante, Borja Camino-Pontes, María Fedetz, Leire Legarreta, Iker Malaina, Gorka Pérez-Yarza, Luis Martínez, Jesús M Cortés, José I. López

## Abstract

Human RNA Polymerase II (Pol II) is characterized by a dense layer of over 775 post-translational modifications (PTMs) that form a dynamic, rewritable regulatory architecture integrating large numbers of cellular signals to coordinate transcription initiation, elongation, termination, and co-transcriptional RNA processing. While genomic information has been extensively catalogued, the potential information capacity associated with Pol II PTM patterns has remained largely unquantified. Here, we analyze the PTM sites across the human Pol II complex (Rpb1–Rpb12) and estimate the state-space information capacity using Shannon entropy theory. We first provide a theoretical upper bound of ~707.98 bits per Pol II molecule (~88.50 bytes) corresponding to ~5.68 × 10^7^ bits per nucleus (~7.10 MB), assuming ~80,200 Pol II molecules per cell. We distinguish this maximal capacity from a conservative, kinetically addressable estimate of ~114.88 bits per molecule (~1.15 MB per nucleus), reflecting physiological kinetic constraints and site coupling that restrict the simultaneously addressable PTM state space in vivo. Finally, we show that major PTM classes (phosphorylation, proline isomerization, O-GlcNAcylation and ubiquitination) operate over distinct lifetimes, from seconds to minutes and hours-scale processes, supporting a multi-timescale biochemical architecture of this enzyme. Together, these results provide a quantitative information framework that distinguishes maximal PTM state-space capacity from kinetically addressable physiological regulatory capacity, supporting a view of Pol II PTM patterning as a high-dimensional, dynamically reconfigurable, multi-timescale regulatory information layer.

**Highlights:** A systems-level framework quantifies regulatory information in Pol II PTMs Known PTM modification sites provide 707.98 bits per Pol II as a regulatory upper bound Physiological kinetic constraints reduce accessible regulatory capacity to 114.88 bits Distinct PTM modification classes define fast, intermediate, and slow regulatory layers

**Graphical Abstract:** 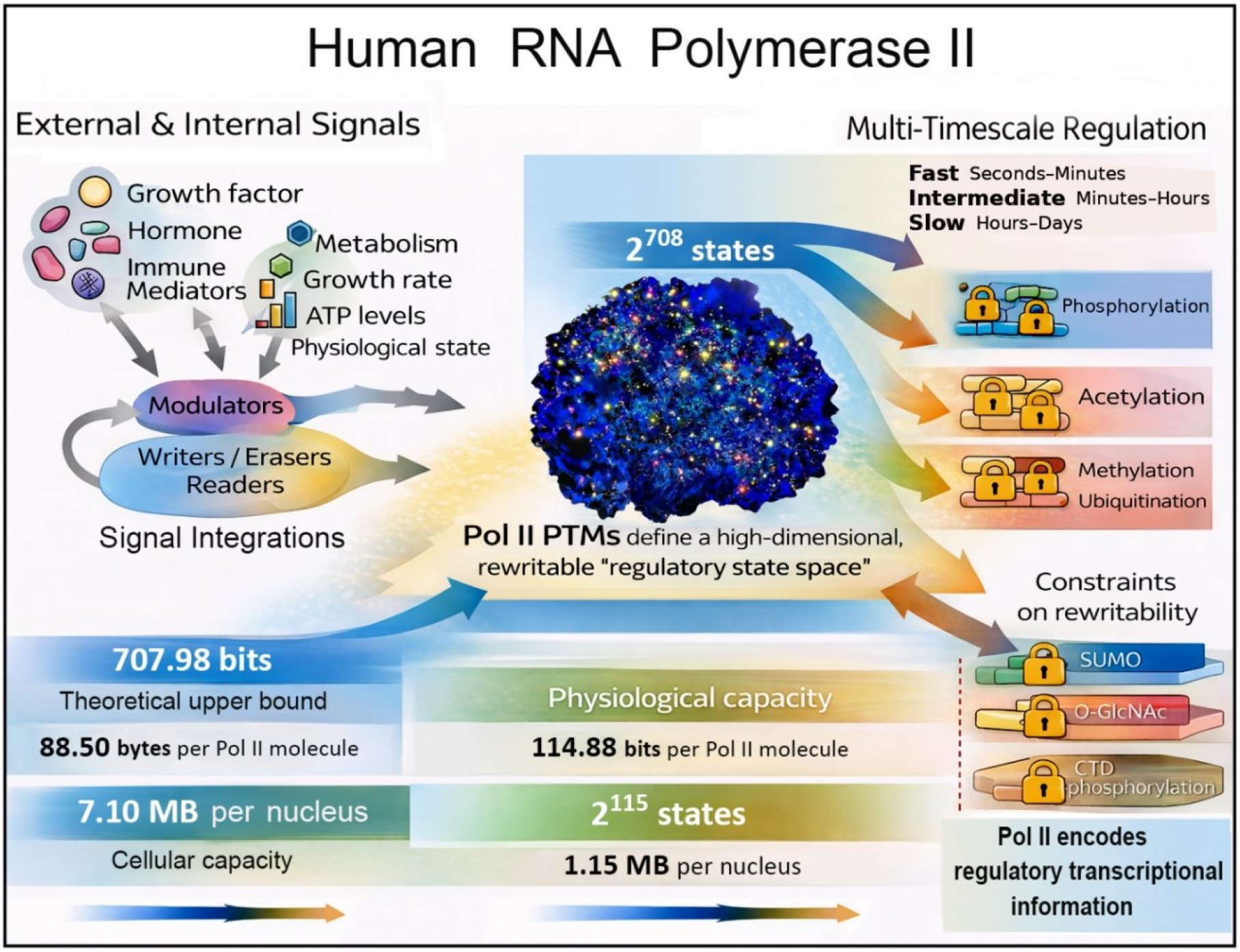

## Introduction

RNA polymerase II (Pol II) is the central enzyme that converts genomic information into RNA output in human cells. This fundamental cellular process allows DNA to be read and RNA transcripts to be synthesized, including all protein-coding mRNAs and multiple classes of non-coding RNAs (e.g., snRNA, miRNA, and snoRNA), in a manner closely adapted to cellular conditions^1^.

To maintain an accurate transcription, Pol II operates within a highly complex regulatory network that helps determine which genomic regions are transcribed, and to what extent, throughout the cellular life, ensuring that only the genomic information necessary for optimal physiological function is transcribed at any given time in the cell cycle^2^.

The activity of this sophisticated enzyme requires a continuous supply of chemical energy from nucleotide hydrolysis, which is converted into mechanical work generating forces on the order of tens of piconewtons during transcription elongation, comparable to or exceeding those reported for many ATP-dependent molecular motors^3^. This force pulls the DNA template through the enzyme, allowing sustained elongation rates spanning from sub-kilobase to several kilobases per minute in live-cell estimates^4^. Despite this speed, Pol II maintains high fidelity through proofreading and backtracking mechanisms, yielding low transcriptional error rates^5^.

To synthesize the appropriate RNAs at each moment, Pol II acts as one of the most complex regulatory hubs in the cell, integrating a wide range of extracellular and intracellular signals. External stimuli, such as hormones^6^, growth factors^7^, heat stress^8^, developmental cues^9^, hypoxia^10^, oxidative stress^11^, and inflammation^12^, modify the transcriptional outcome. Pol II also responds to intracellular variables including ATP availability^13^, growth rate^14^, metabolic state and nutrient availability^15,16^, endocytic signaling^17^, endoplasmic reticulum stress responses^18^, mitochondrial activity^19^, cytoskeletal reorganization^20^, cell-cycle state^21^, chromatin organization^22^, circadian programs^23^, and DNA damage responses^24^.

Mechanistically, Pol II operates within a large and complex dynamic molecular network that primarily involves the general transcription factors, the Mediator complex, chromatin remodelers, RNA-processing machinery, elongation/pausing regulators, and termination processes^25^. Throughout the transcription cycle, Pol II engages numerous enzymatic activities, including kinases, phosphatases, helicases, translocases, methyltransferases, acetyltransferases, exonucleases, ubiquitin ligases, and cyclin-dependent kinases, that continuously tune transcriptional dynamics in response to cellular context^26–30^. As a result, individual human cells exhibit rapid and continuous transcriptional adaptation, producing thousands of nascent transcripts at any given time^31^.

A key reason Pol II can integrate so many inputs is its extensive and reversible layer of post-translational modifications (PTMs), in which chemical groups such as phosphate, methyl, or acetyl moieties are covalently attached to specific amino acid residues. These modifications can rapidly reconfigure interaction surfaces, alter the binding avidity of regulatory proteins, and modulate transitions between transcriptional steps^32^. In this sense, Pol II can encode regulatory information through combinatorial patterns of PTMs that are written, erased, and interpreted by dedicated enzymes and reader proteins, enabling context-dependent control of transcriptional output^33–35^. As a consequence, these covalent PTM marks provide a rapid and reversible mechanism for integrating environmental and intracellular signals into the structure and functionality of Pol II^36^, establishing an information-rich molecular landscape that orchestrates all phases of the transcription cycle^32^.

Despite significant progress in cataloging covalent modifications in Pol II^37–41^, a central question remains unanswered: what is the potential information capacity of these biochemical covalent marks? While the genomic information content has been studied, for example, in ENCODE Project Consortium^42^, a comparable quantification for Pol II is lacking. Without such quantification, the magnitude of this regulatory layer remains conceptually imprecise, despite strong links between PTM dysregulation and disease-related phenotypes^43^.

Here we address this gap using Shannon information theory applied to residue-level PTM state spaces. We compiled the human Pol II PTM sites from the experimental scientific literature, publicly available databases and defined explicit discrete-state models for major PTM classes. Under independent and equal-probability assumptions, we estimate an upper-bound theoretical capacity of ~707.98 bits per Pol II molecule, equivalent to ~5.68 × 10^7^ bits per nucleus (~7.10 MB) given ~80,200 Pol II molecules per cell. We further separate this theoretical capacity from a physiological/kinetically accessible *in vivo* bound, estimated at ~114.88 bits per Pol II molecule (~9.21 × 10^6^ bits per nucleus, ~1.15 MB, assuming ~80,200 Pol II), reflecting that only a subset of all covalent marks can be independently established and stably maintained within a given time window.

Furthermore, we show that these covalent biochemical marks span distinct lifetimes revealing a complex multi-timescale architecture. In fact, phosphorylation is widely understood as encoding regulatory responses^44^ lasting ~5 seconds to >15 minutes^45,46^, proline isomerization^47,48^ and O-GlcNAcylation^49–51^ form a mesoscale temporal structure spanning from ~1 min to ~30 minutes, and ubiquitination establishes a slower transient process lasting from ~15 min to ~24 hours^52,53^.

This information-theoretic perspective positions Pol II not only as an exceptional transcriptional machine but also as a complex information-processing system across multiple timescales. The framework is grounded in curated PTM resources and a broad experimental literature. All numerical results shown here are fully reproducible using the Python Modules provided in Supplementary Information S4, which regenerate all tables, PTM counts, capacity values, physiological estimates, sensitivity scenarios, and discrete-state models.

## Results

We have systematically analyzed the canonical residue-level PTM states in human Pol II and calculated their Shannon information content. Specifically, this **~**500 kDa enzyme comprises 12 subunits (Rpb1– Rpb12) that form a highly conserved catalytic core, structurally and mechanistically maintained from archaea to humans^54^. The largest subunit, Rpb1, contains the extended C-terminal domain (CTD), which serves as the main regulatory scaffold for Pol II. The CTD undergoes extensive, reversible PTMs enabling it to coordinate virtually all phases of the transcription cycle, including initiation, promoter escape, early and productive elongation, termination, and co-transcriptional RNA processing^44^ ( see for more details Supplementary Information S1).

### I. Combinatorial PTM state space of the mammalian CTD

The C-terminal domain (CTD) of Rpb1 comprises 52 Tyr-starting heptad repeats related to the consensus Tyr1–Ser2–Pro3–Thr4–Ser5–Pro6–Ser7 (YSPTSPS) and a 10-residue C-terminal motif^55^. In the canonical human sequence only 21 repeats match the exact consensus, whereas the remaining 31 are nonconsensus with substitutions primarily at positions 2, 4, 5, and especially 7; notably, eight of them contain Lys (K7) and two contain Arg (R4 and R7) (The canonical UniProt POLR2A sequence, P24928^37^). These CTD residues (Table 1) are extensively modified by multiple PTM types, generating an exceptionally large combinatorial regulatory space. The covalent modifications considered include phosphorylation, acetylation, O-GlcNAcylation, mono-, di- and trimethylation, asymmetric dimethylation, symmetric dimethylation, SUMOylation and ubiquitination (retrieved from PTM resources UniProt^37^, iPTMnet^38^, PhosphoSitePlus^39^, GlyGen^40^ and dbPTM^41^), and proline isomerization^47,48,56–60^, two additional O-GlcNAcylation marks^49^, and citrullination^61^ (compiled from the literature).

**Table 1.**
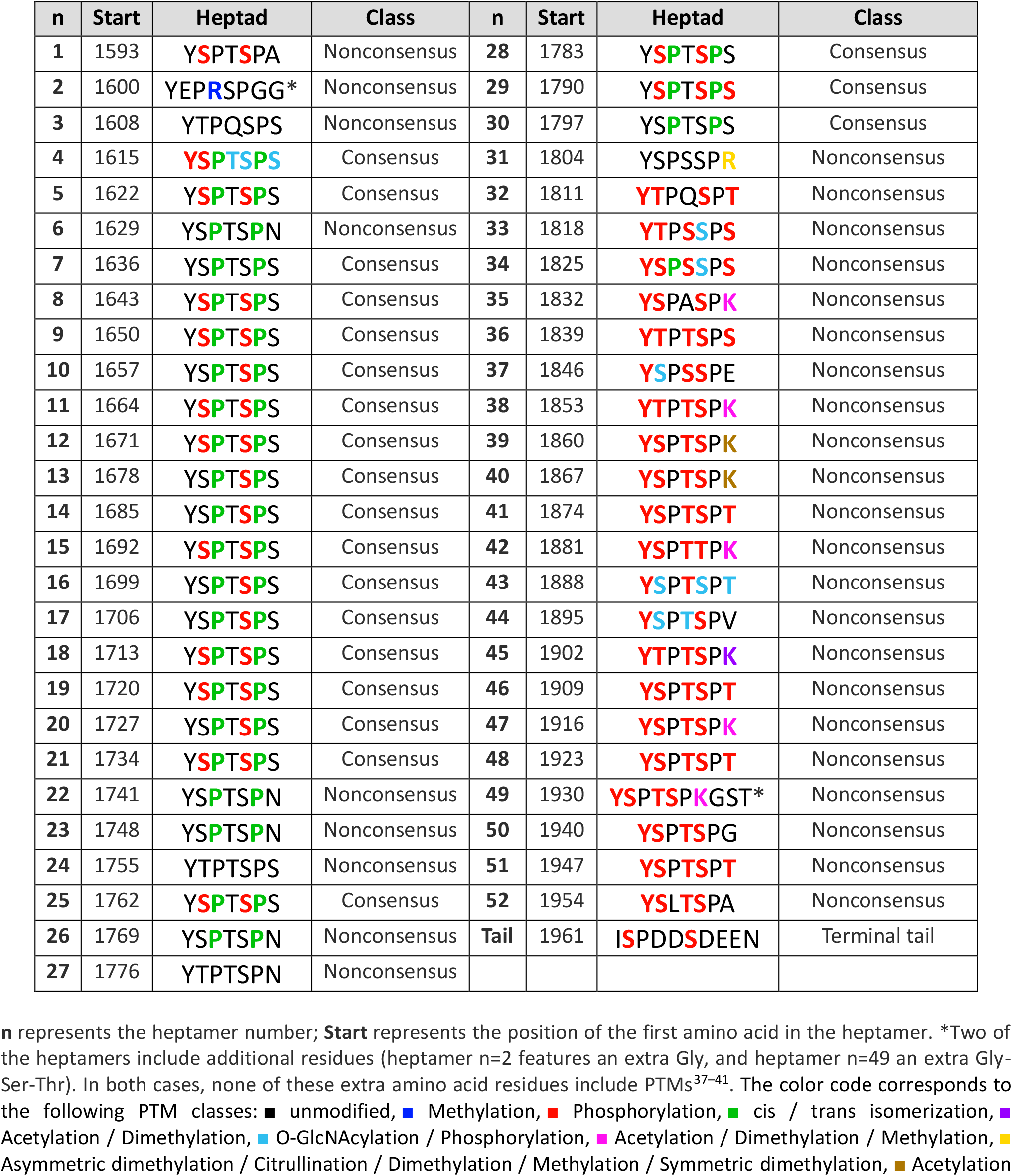
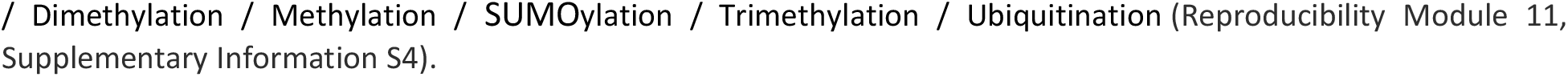
CTD structure comprising 52 heptapeptides and a 10-residue C-terminal tail (UniProt P24928 canonical sequence).

Quantitatively, the total CTD state-space size can be computed as:

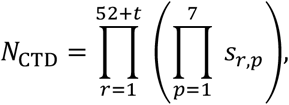

where *s*_*r,p*_ is the number of discrete states assigned to position *p* of repeat *r* according to the residue identity at that position (e.g., *s* = 2 for Tyr1; and *s* = 4 for the single Arg7; *t* corresponds to the 10-amino-acid residue C-terminal sequence.

Using the 52 heptads from UniProt P24928 (Table 1) plus the 10-residue C-terminal tail, and applying this residue-exact state mapping (Table 2) yields a CTD state-space size corresponding to:

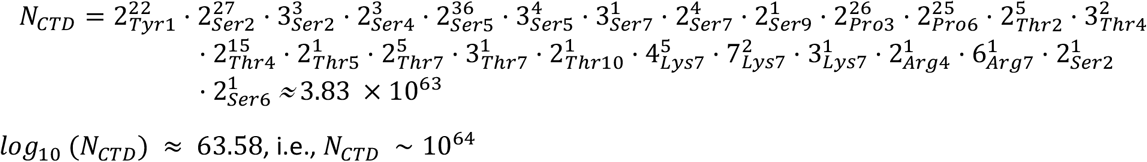

**Table 2.**
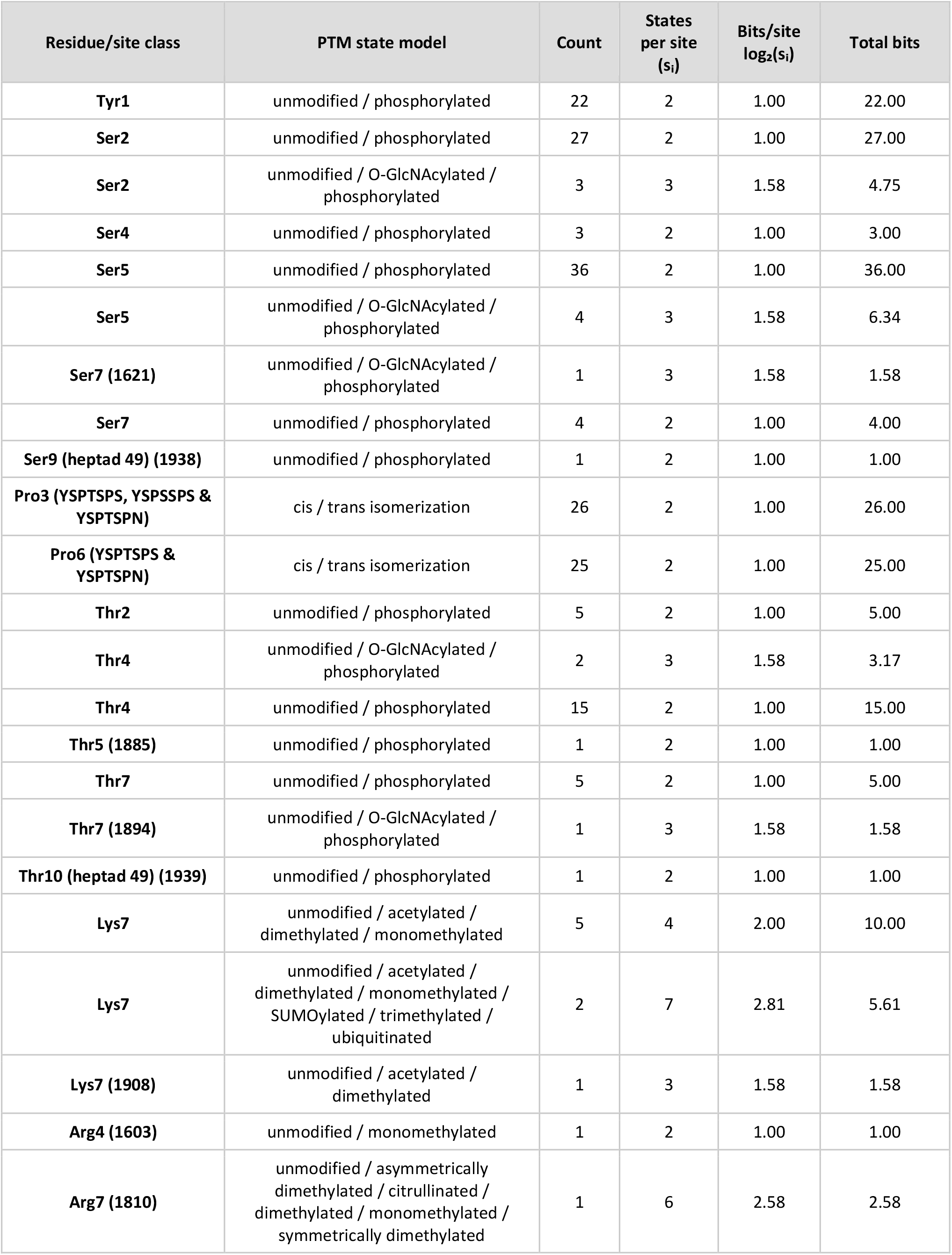
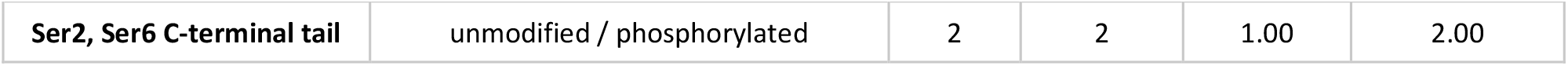
Residue-level information and contribution per PTM of the CTD (Rpb1 subunit)

| Residue/site class | PTM state model | Count | States per site ($s_i$) | Bits/site $\log_2(s_i)$ | Total bits |
| --- | --- | --- | --- | --- | --- |
| **Tyr1** | unmodified / phosphorylated | 22 | 2 | 1.00 | 22.00 |
| **Ser2** | unmodified / phosphorylated | 27 | 2 | 1.00 | 27.00 |
| **Ser2** | unmodified / O-GlcNAcylated / phosphorylated | 3 | 3 | 1.58 | 4.75 |
| **Ser4** | unmodified / phosphorylated | 3 | 2 | 1.00 | 3.00 |
| **Ser5** | unmodified / phosphorylated | 36 | 2 | 1.00 | 36.00 |
| **Ser5** | unmodified / O-GlcNAcylated / phosphorylated | 4 | 3 | 1.58 | 6.34 |
| **Ser7 (1621)** | unmodified / O-GlcNAcylated / phosphorylated | 1 | 3 | 1.58 | 1.58 |
| **Ser7** | unmodified / phosphorylated | 4 | 2 | 1.00 | 4.00 |
| **Ser9 (heptad 49) (1938)** | unmodified / phosphorylated | 1 | 2 | 1.00 | 1.00 |
| **Pro3 (YSPTSPS, YSPSSPS & YSPTSPN)** | cis / trans isomerization | 26 | 2 | 1.00 | 26.00 |
| **Pro6 (YSPTSPS & YSPTSPN)** | cis / trans isomerization | 25 | 2 | 1.00 | 25.00 |
| **Thr2** | unmodified / phosphorylated | 5 | 2 | 1.00 | 5.00 |
| **Thr4** | unmodified / O-GlcNAcylated / phosphorylated | 2 | 3 | 1.58 | 3.17 |
| **Thr4** | unmodified / phosphorylated | 15 | 2 | 1.00 | 15.00 |
| **Thr5 (1885)** | unmodified / phosphorylated | 1 | 2 | 1.00 | 1.00 |
| **Thr7** | unmodified / phosphorylated | 5 | 2 | 1.00 | 5.00 |
| **Thr7 (1894)** | unmodified / O-GlcNAcylated / phosphorylated | 1 | 3 | 1.58 | 1.58 |
| **Thr10 (heptad 49) (1939)** | unmodified / phosphorylated | 1 | 2 | 1.00 | 1.00 |
| **Lys7** | unmodified / acetylated / dimethylated / monomethylated | 5 | 4 | 2.00 | 10.00 |
| **Lys7** | unmodified / acetylated / dimethylated / monomethylated / SUMOylated / trimethylated / ubiquitinated | 2 | 7 | 2.81 | 5.61 |
| **Lys7 (1908)** | unmodified / acetylated / dimethylated | 1 | 3 | 1.58 | 1.58 |
| **Arg4 (1603)** | unmodified / monomethylated | 1 | 2 | 1.00 | 1.00 |
| **Arg7 (1810)** | unmodified / asymmetrically dimethylated / citrullinated / dimethylated / monomethylated / symmetrically dimethylated | 1 | 6 | 2.58 | 2.58 |
| Ser2, Ser6 C-terminal tail | unmodified / phosphorylated | 2 | 2 | 1.00 | 2.00 |

This state-space size (Reproducibility Module 12, Supplementary Information S4) is conservative with respect to richer multi-mark models but remains an upper-bound capacity relative to in vivo occupancy distributions. Furthermore, PTMs on the remaining Pol II subunits (Rpb1_core_–Rpb12) were not included, which would increase the theoretical state space by several additional orders of magnitude. Even under these assumptions, the combinatorial capacity of the CTD is enormous, encoding a combinatorial state space on the order of 10^64^. In vivo, only a biologically constrained subset is kinetically accessible at any given time due to transcription-cycle ordering, enzymatic coupling, and non-uniform state occupancies. Nevertheless, the magnitude of the CTD’s potential state space is consistent with its role as a dynamic regulatory scaffold capable of integrating diverse cellular signals and coordinating transitions across the transcription cycle.

### II. Maximal information capacity of CTD PTM patterns

To quantify the maximal information capacity of CTD we use Shannon’s measure of state entropy^62^:

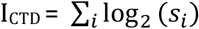

Taking into account the number of discrete states (*s*_*i*_), determined by the corresponding PTMs, the residue-level information, and considering all 52 heptapeptide repeats and the C-terminal tail (Table 2), the maximal information capacity in a human P24928 CTD yields the explicit sum:

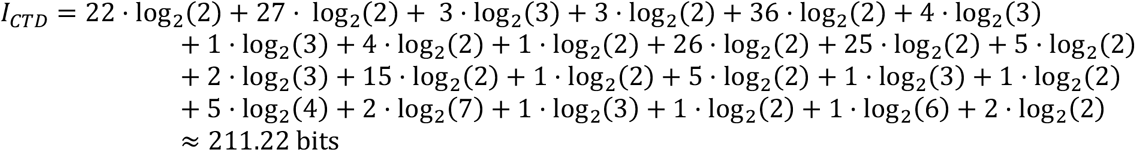

This value (Reproducibility Module 3, Supplementary Information 4) represents the upper bound of the theoretical information encoded by the known CTD PTMs, assuming independence among sites and equal probability across available states. In vivo, state probabilities are highly skewed and constrained by transcriptional stage, enzyme activity, and chromatin context; thus, the achieved entropy is expected to be lower. Nevertheless, the CTD functions as a high-dimensional regulatory medium capable of encoding complex cellular states through diverse PTM combinations (Figure 1).

**Figure 1.**
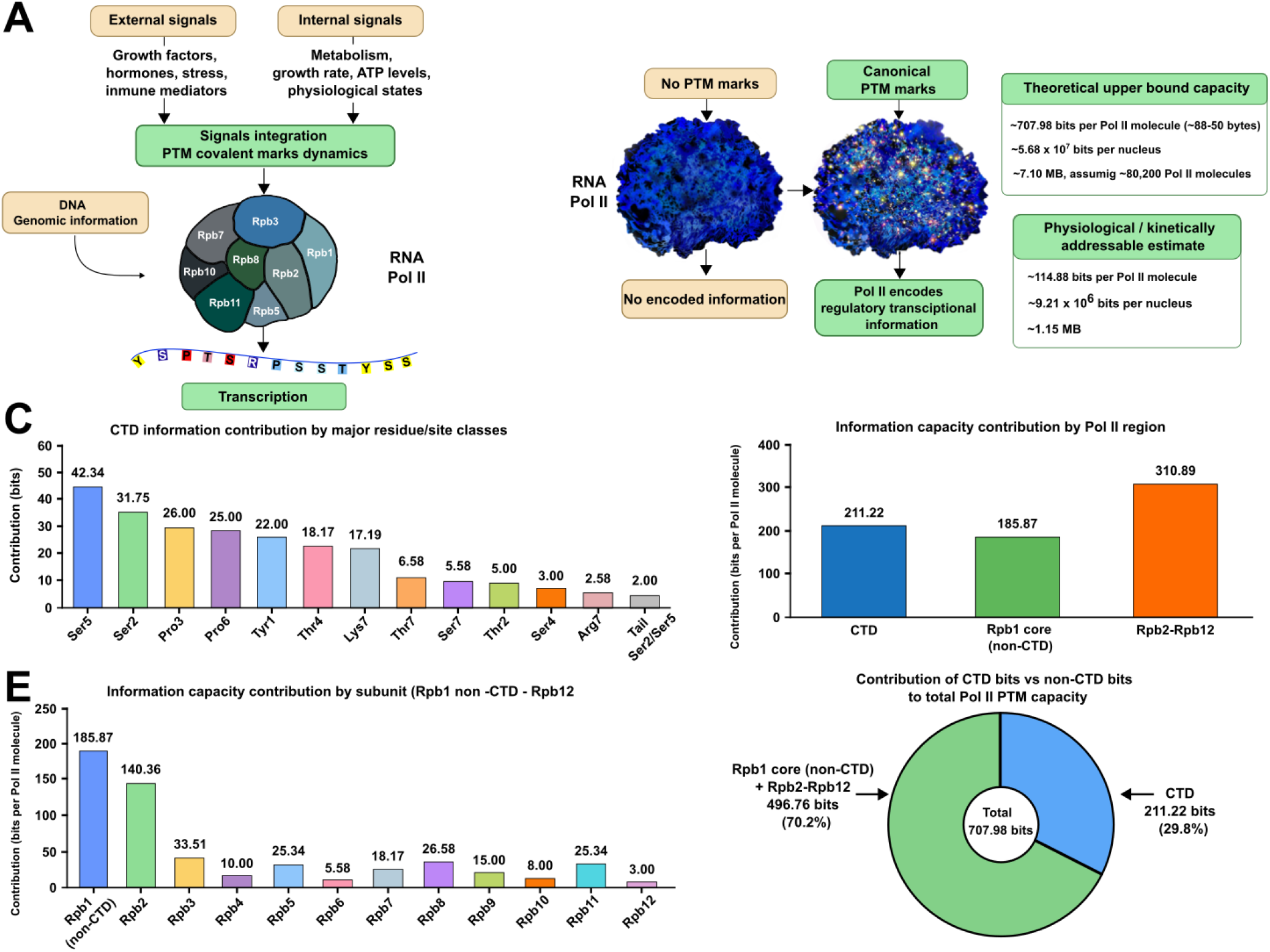
Information capacity encoded by post-translational modification patterning across human RNA polymerase II. A, Schematic representation of external and internal cellular signals converging on RNA polymerase II (Pol II) through dynamic post-translational modification (PTM) marks. PTM writers, erasers, and readers convert environmental and intracellular cues into reversible covalent marks that modulate Pol II function during transcription. B, Conceptual comparison between an unmodified Pol II molecule and a canonically PTM-decorated Pol II molecule. In the absence of PTM marks, no PTM-encoded regulatory information is represented, whereas canonical PTM marks define a high-dimensional regulatory state space. The theoretical upper-bound capacity of the full human Pol II PTM landscape is estimated at ~707.98 bits per Pol II molecule, equivalent to ~88.50 bytes, ~5.68 × 10^7 bits per nucleus, or ~7.10 MB assuming ~80,200 Pol II molecules per nucleus. The conservative kinetically addressable estimate is ~114.88 bits per Pol II molecule, corresponding to ~9.21 × 10^6 bits per nucleus or ~1.15 MB. C, Contribution of major CTD residue/site classes to CTD information capacity. Values represent summed residue-class contributions under the curated discrete-state model. D, Regional contribution to total Pol II PTM information capacity: CTD, Rpb1 core non-CTD, and Rpb2–Rpb12. E, Per-subunit contribution of Rpb1 core non-CTD and Rpb2–Rpb12 to the non-CTD PTM information capacity. F, Relative contribution of CTD and non-CTD regions to the total theoretical Pol II PTM capacity. CTD contributes 211.22 bits, whereas Rpb1 core non-CTD plus Rpb2–Rpb12 contribute 496.76 bits, yielding a total of 707.98 bits per Pol II molecule.

Counts in Table 2 refer to residue instances for which a PTM state was included in the curated CTD model; residues present in the CTD sequence but lacking curated PTM evidence were assigned a single unmodified state and therefore contribute 0 bits. I_CTD_ = ∑_*i*_ log_2_ (*s*_*i*_) = 211.22 bits. (Reproducibility Module 13, Supplementary Information S4).

### III. Information capacity of PTMs on the Rpb1 core (non-CTD)

Because Rpb1 PTM biology is not confined to the CTD, we separately quantified the human non-CTD Rpb1 PTM sites from UniProt, dbPTM, iPTMnet, PhosphoSitePlus and GlyGen (POLR2A / P24928), across residues 1–1592 (i.e., excluding the heptad-repeat CTD region to avoid double counting). These sites include acetylation, glutathionylation, methylation, O-GlcNAcylation, phosphorylation, sulfoxidation, SUMOylation and ubiquitination annotations (see Methods Section and Supplementary Table S1).

Using the same mutually exclusive state model, this conservative non-CTD set (208 PTMs across 159 unique modified residues) yields (Reproducibility Module 5, Supplementary Information 4):

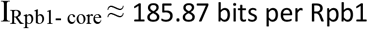

### IV. Information capacity encoded by PTMs in the subunits Rpb2–Rpb12

Although the CTD is the primary PTM-rich region of Pol II, structural subunits Rpb2–Rpb12 also carry numerous functionally relevant covalent modifications. Using human entries from UniProt, dbPTM, iPTMnet, PhosphoSitePlus and GlyGen, we compiled 278 residues modified by 338 unique PTM marks including acetylation, methylation, O-GlcNAcylation, phosphorylation, sulfoxidation, SUMOylation and ubiquitination. For each modified residue, and these individual PTM types the number of equally probable discrete states is *s*_*i*_ = 2. (Supplementary Tables S2–S12). When multiple PTM types act on the same residue we treat modifications as mutually exclusive (e.g., lysines modified by either ubiquitin or SUMO). Then, the composite state counts were calculated as:

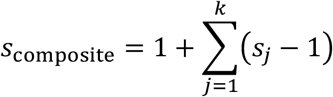

where *s*_*j*_ is the number of states for PTM type *j* at that residue. For example, a lysine capable of ubiquitination (*s*= 2) or SUMOylation (*s*= 2) yields *s*_composite_ = 3, corresponding to Shannon information log_2_(3) ≈ 1.58 bits.

The compiled dataset of Rpb2–Rpb12 PTM sites (Supplementary Tables S2-S12) shows the following aggregate contributions: phosphorylation sites in multiple serine, threonine, and tyrosine residues across subunits, each of them with *s*=2 unless part of a composite site. Acetylation sites in Alanines, Serines, lysines and N-terminal methionines, with *s*=2 unless in composite. Ubiquitination sites in lysines, each with *s*=2, except when sharing a residue with another PTM type. SUMOylation sites in lysines, with *s*=2, likewise adjusted in composites. Additionally, sulfoxidation is observed on methionine residues (three sites) and O-GlcNAcylation on one threonine residue, each with s=2. Finally, methylation sites in cysteines, lysines, methionines and arginines, with *s*=2. Note that virtually all composites involve lysines combining ubiquitination with one or more of SUMOylation, acetylation or methylation, giving s=3 or s=4. The one exception is S2 in Rpb6, which carries acetylation plus phosphorylation (s=3). After consolidating multi-PTM residues into composite sites the maximum theoretical information content in Rpb2–Rpb12 residues yields:

Rpb2: ~140.36 bits

Rpb3: ~33.51 bits

Rpb4: 10.00 bits

Rpb5: ~25.34 bits

Rpb6: ~5.58 bits

Rpb7: ~18.17 bits

Rpb8: ~26.58 bits

Rpb9: 15.00 bits

Rpb10: 8.00 bits

Rpb11: ~25.34 bits

Rpb12: 3.00 bits

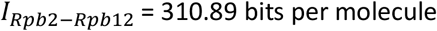

This information is equivalent to ≈ 38.86 bytes, per all Rpb2–Rpb12 complex (Reproducibility Module 14, Supplementary Information S4).

### V. Total information capacity per Pol II molecule

Summing the three components (CTD + Rpb1 core + Rpb2–Rpb12): the combined information capacity of one human RNA Pol II molecule is:

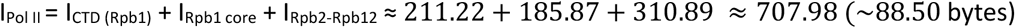

Thus, the maximal theoretical information capacity encoded by known PTM patterns on each human Pol II molecule corresponds to ~88.50 bytes of regulatory information (see Figure 1), distributed across dynamic and reversible covalent marks on Rpb1–Rpb12 (Reproducibility Module 15, Supplementary Information S4). This theoretical molecular information layer is reconfigurable and may support complex modulation of transcription through reversible chemical modifications.

### VI. Cellular level PTM information capacity of Pol II

Using an estimate of M ≈ 80,200 Pol II molecules per mammalian tissue-culture nucleus (experimental studies based on super-resolution microscopy, BioNumbers 112321^63^) the total information capacity at cellular level is:

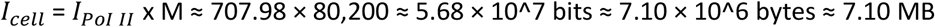

This estimate (Reproducibility Module 15, Supplementary Information S4) represents a theoretical upper bound, derived from known PTMs assuming independence between sites, equal probability between states, and equal weighting of Pol II molecules. However, the accessible state space is strongly constrained by: transcriptional stage–specific PTM covalent marks, structural coupling between them, enzyme pathway wiring (kinases, phosphatases, CTD readers, chromatin state and transcriptional context). Thus, while Pol II carries a theoretical regulatory capacity on the order of a few megabytes, only a biologically meaningful subspace is used “in vivo”. Here we report state-space information capacity (i.e., log_2_(N) under an independent-site, equal-probability model), not the achieved Shannon entropy of “in vivo” PTM occupancy distributions, which is expected to be lower due to non-uniform occupancies and site coupling (see Supplementary Information S2). Nevertheless, this quantitative framework highlights the remarkable potential of PTM patterning to encode regulatory states at the molecular scale. In addition, sensitivity analysis shows that the high-dimensional Pol II PTM information capacity is robust to plausible alternative discrete-state model choices (Supplementary Information S3).

### VII. Pol II PTMs form a multi-timescale biochemical functional architecture

Beyond static capacity, Pol II encodes regulatory information not only through the diversity of PTMs but also via the distinct kinetic lifetimes of their covalent marks and reversal mechanisms, generating a multi-timescale biochemical architecture (Figure 2). In fact, different PTM classes operate over fast, intermediate, and slow time scales, allowing the enzyme to encode and integrate signaling history over different temporal windows. This physiological information bound can be computed as:

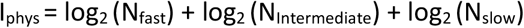

**Figure 2.**
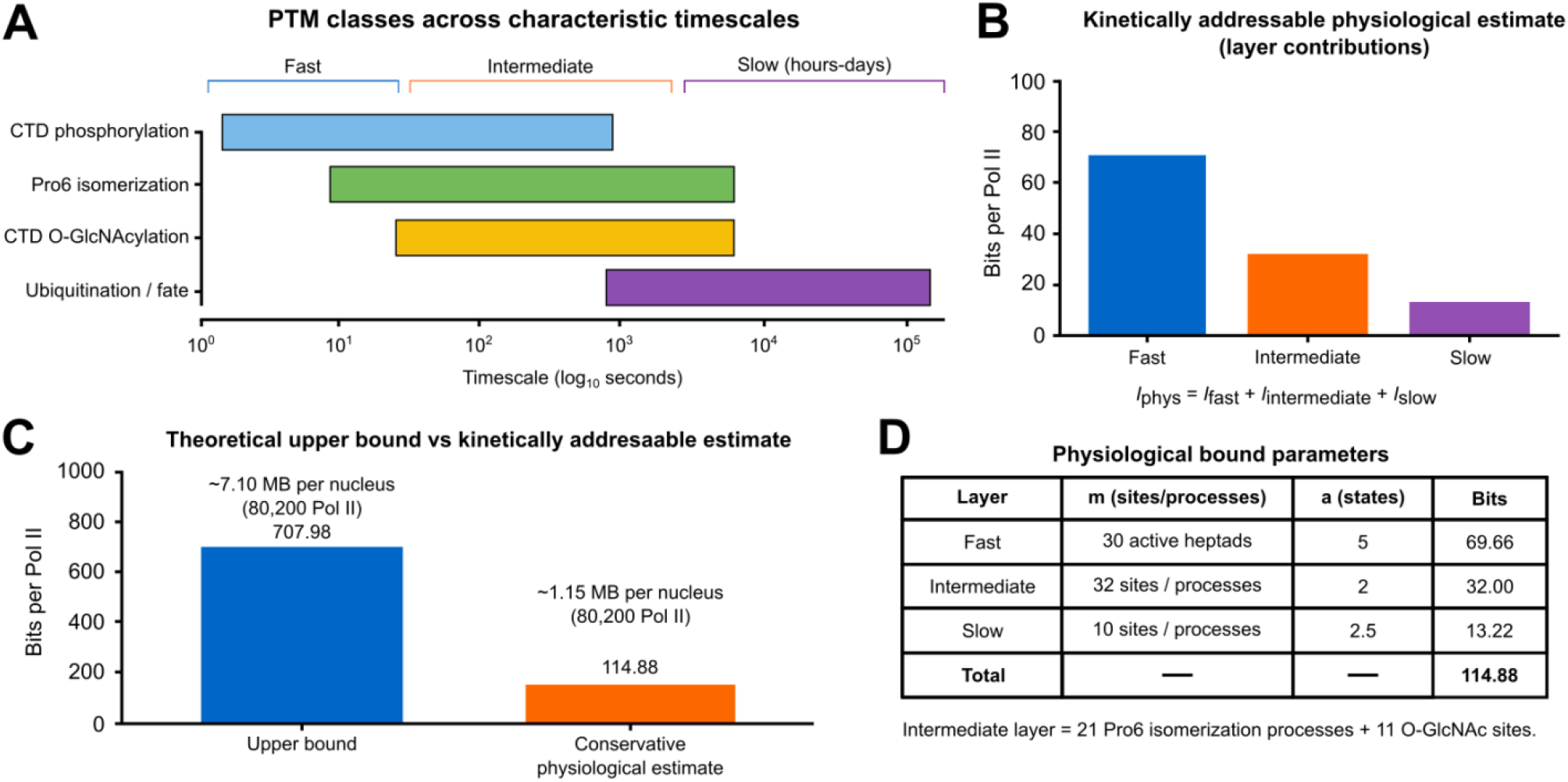
Multi-timescale physiological architecture of Pol II PTM information. A, Characteristic kinetic regimes of major Pol II PTM classes included in the physiologically addressable model. CTD phosphorylation operates predominantly on fast seconds-to-minutes timescales, Pro6 cis/trans isomerization and CTD O-GlcNAcylation define an intermediate minutes-scale layer, and ubiquitination-linked Pol II fate decisions extend into slower hours-to-days regulatory windows. B, Layer-wise contribution to the conservative kinetically addressable physiological estimate. The fast layer contributes 69.66 bits per Pol II molecule, the intermediate layer contributes 32.00 bits, and the slow layer contributes 13.22 bits, giving a total of 114.88 bits per Pol II molecule. C, Comparison between the theoretical upper-bound PTM state-space capacity and the conservative physiological estimate. The theoretical upper bound is 707.98 bits per Pol II molecule, corresponding to ~7.10 MB per nucleus assuming ~80,200 Pol II molecules, whereas the conservative physiological estimate is 114.88 bits per Pol II molecule, corresponding to ~1.15 MB per nucleus. D, Parameters used for the coarse-grained physiological estimate. The fast layer was modeled as 30 active heptads with 5 kinetically distinguishable states per heptad; the intermediate layer as 32 sites/processes with 2 states each; and the slow layer as 10 sites/processes with an average of 2.5 distinguishable states. The intermediate layer comprises 21 Pro6 isomerization processes and 11 CTD O-GlcNAc sites.

For each kinetic layer k, we define:

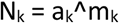

where m_k_ is the number of effective sites or repeats and a_k_ is the effective number of kinetically distinguishable states per site after accounting for coupling and kinetic constraints. Therefore:

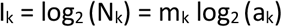

### VII-I. Fast-timescale layer (seconds–minutes)

Phosphorylation of the C-terminal domain at Tyr1, Ser2, Ser5, and Ser7 residues, mediates rapid transcriptional responses^64–66^. Kinetic studies indicate that phosphorylation and dephosphorylation cycles occur on the order of ~5 seconds to up to >15 min^45,46^. Phosphorylatable CTD heptads have an average ~3.12 phosphorylated residues (Table 1) that can be either unmodified or phosphorylated^67^ (2^3^ = 8 states per heptad). Although the canonical CTD sequence contains 52 heptads, CTD mass spectrometry in human cells shows sparse phosphorylation^67,68^, with stoichiometry studies in human CTD estimating ~30 phosphorylated heptads per CTD^69^. However, the effective distinguishable states are further reduced due to kinetic coupling and electrostatic and steric constraints, specifically to ~5 per heptad (unmodified state, ~3 distinct monophosphorylated states and a rarer double phosphorylated state)^66,67,70,71^.

We thus selected m_fast_ ≈ 30 modified heptads and a_fast_ ≈ 5 as a conservative effective-state value within the restricted kinetically distinguishable configurations per active heptad. The Shannon information is:

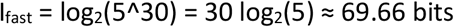

Therefore, the physiological information capacity of this layer is approximately 70 bits per Pol II molecule (Figure 2), which is equivalent to 5^30 ≈ 9.31 × 10^20 distinguishable rapid-response CTD states (Reproducibility Module 16, Supplementary Information S4). This short transient information enables Pol II to give dynamically fast responses to extracellular cues and numerous intracellular variables, coordinating transcription initiation, elongation, and promoter-proximal pausing in real time.

### VII-II. Intermediate layer (minutes to hours)

Biochemical time-course kinetic assays indicated a number of 21 Pro6 isomerization processes along the Ser5-Pro6 bonds of consensus heptad sequences (YSPT**S-P**S), in the human CTD^37^, which spans the scale of ~30 seconds (catalyzed) to up to ~30 min (uncatalyzed)^47,48^. Therefore we have m_Pro6_ ≈ 21 and a_Pro6_ = 2. Likewise, kinetic and functional studies show that O-GlcNAcylation addition and removal cycles occur within < 2min to ~30 min^49–51^. PTM databases reported nine O-GlcNAc marks in the human CTD (eight by GlyGen, and one by PhosphoSitePlus) and two more are reported by experimental works with human CTD using LC-MS/MS^49^. Therefore, m_O-GlcNAc_ ≈ 11, with a_O-GlcNAc_ = 2.

Thus, for Pro6 isomerization and O-GlcNAcylation processes we obtain a number of effectively modified sites of m_Intermediate_ = 32, and a_Intermediate_ = 2 kinetically distinguishable states per site. The information contained in both processes according to Shannon information theory is:

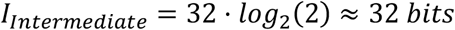

This mesoscale physiological information corresponds to N_intermediate_ ≈ 2^32^ ≈ 4.29 × 10^9^ possible functional configuration states (Reproducibility Module 16, Supplementary Information S4). The Pol II mesoscale information represents a notable reservoir of mid-term regulatory history (Figure 2). Intermediate PTM modifications integrate metabolic signals, chromatin context, and cofactor interactions, providing a transient regulatory layer that sustains transcriptional adaptations over tens of minutes, bridging rapid signaling events and mesoscale-term regulatory processes.

### VII-III. Slow layer (hours–days)

Rpb1 ubiquitination, including K48- and K63-linked ubiquitin chains on Rpb1 K1268, coordinates Pol II degradation, repair, and transcription recovery after DNA damage^24,72^. Ubiquitination establishes a slower transient process that can persist over longer windows, from ~15 min to ~24 hours, and influences the available Pol II pool^24,52,53^. This slower timescale encodes information about long-term cellular perturbations such as DNA damage, oxidative stress, and arrested or defective Pol II states, influencing the pool of available Pol II molecules and thereby shaping transcriptional output over extended periods^24,73,74^. Experimentally, across the entire 12-subunit human RNA Pol II complex proteomics/PTM resources report on the order of **~**120 distinct ubiquitinated residues in total^37–41^, but only a biologically constrained subset is kinetically accessible at any given time under physiological conditions. Assuming a very conservative estimation for ubiquitination, of m_slow_ ~10 slow sites/processes, with an average a_slow_ ~2.5 distinguishable states per site (capturing long-lived outcomes such as stable ubiquitination-linked states) the Shannon information is:

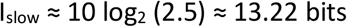

This physiological slow process encodes roughly 13.2 bits per Pol II molecule, corresponding to a potential space of ~9.54 × 10^3 (2.5^10) of functional configuration states. This layer is smaller than the rapid one (~70 bits) but extremely powerful because it persists for up to ~24 h influencing Pol II stability and genome-wide transcriptional readiness (Reproducibility Module 16, Supplementary Information S4).

Timescale marks such as acetylation, SUMOylation, and methylation are not explicitly parameterized here because current condition-resolved kinetic occupancy experimental data are insufficient to define a defensible effective state model.

### VII-IV. Integrative Pol II physiological multi-timescale architecture

Combining these timescales, Pol II PTMs establish a biochemical dynamic structure, from transient second-scale responses to longer-lived hours-to-days regulatory states. This hierarchy enables the cell to integrate recent transcriptional events with transient physiological trends. Thus, the conservative kinetically addressable estimate is:

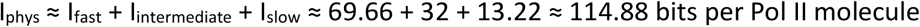

This physiological value (Reproducibility Module 16, Supplementary Information S4) is a conservative estimate of kinetically addressable information, distinct from the theoretical upper bound derived from the full **~**707.98 bits residue-level state space (Results: Section V). The remaining **~**593.10 bits (707.98 – 114.88 bits) reflect static combinatorial potential not fully expressed simultaneously due to kinetic constraints (Figure 2). Only a subset of PTMs operates in any given temporal window, therefore, an individual Pol II molecule cannot biologically sample all theoretical states in real time. Nevertheless, 114.88 bits of kinetically accessible, timescale-separated information is extraordinarily notable for a single protein.

At M = 80,200 Pol II molecules per nucleus (BioNumbers 112321^63^):

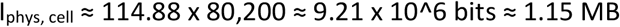

This physiological bound is kinetically addressable information, which represents a remarkable amount of physiological transcriptional value at the cellular level, distinct from the theoretical upper bound derived from the full residue-level state space. Together, these experimentally motivated estimates provide a quantitative framework that reframes Pol II PTM patterning as a high-dimensional, dynamically reconfigurable, multi-timescale information-processing system.

## Discussion

Pol II activity is central to human cellular regulation, helping ensure that only the specific genomic information required for optimal physiological processes is transcribed at any given moment of the cell cycle.

This macromolecular enzyme operates within a complex regulatory network that is tuned at multiple stages by transcription factors, activators and repressors, coactivators, and chromatin-modifying enzymes. These regulators interact in complex ways with Pol II to ensure that specific genomic information is expressed at the right time, in the appropriate amounts, and in any particular cell type, in response to diverse environmental and intracellular signals^36^.

A fundamental feature of Pol II is its dense layer of more than 775 PTMs, which together form a dynamic molecular marking system that integrates a great number of cellular signaling pathways and physiological cues modulating transcription initiation, elongation, and termination. These PTMs, most prominently phosphorylation, acetylation, ubiquitination, glycosylation, methylation, isomerization, citrullination, and SUMOylation, act as a tightly regulated, reversible control layer that coordinates transcription through combinatorial patterns^36,75^. Although the genomic information content of the cell has been well-documented (ENCODE Project Consortium 2012^42^), the potential information capacity associated with covalent marks of Pol II remains largely unexplored.

In this work, we provide a quantitative framework for understanding Pol II PTM patterning as a distributed regulatory information layer across the full enzyme (Rpb1–Rpb12). Under a discrete-state model and Shannon state-entropy assumptions (independence and equal state probabilities), we first estimate a theoretical maximum information capacity of ~707.98 bits per Pol II molecule. At the cellular level, assuming ~80,200 Pol II molecules per nucleus, this corresponds to ~5.68 × 10^7 bits (~7.10 MB). These values are explicit upper bounds and do not imply that all states are sampled, occupied, or biologically accessible simultaneously in vivo. PTM occupancies are constrained by transcriptional stage, enzymatic wiring (writers/erasers/readers), and structural coupling among sites, so only a subset of PTMs is active within any given temporal window.

Consistent with these constraints we estimated a conservative kinetically addressable value of ~114.88 bits per molecule (~1.15 MB per nucleus). This distinction is essential since the theoretical capacity reflects the size of the state space defined by known covalent marks, whereas the physiological capacity reflects only what can plausibly be written, maintained, and read given kinetic and structural constraints.

The results also emphasize the vast combinatorial potential of Pol II PTM patterning. For the CTD alone, the modeled PTM pattern space is in the order of ~10^64^ possible configurations, illustrating the scale of the regulatory landscape even under conservative mutual-exclusivity assumptions. While cells sample only a small and biologically meaningful subset of this space, the size of the underlying state space helps contextualize why PTM patterning can support rich and adaptable regulation.

Importantly, our physiological and kinetic analysis also suggests that Pol II encodes regulatory information not only through the combinatorial diversity of PTMs forming a flexible mark pattern code^76^. Enzymes responsible for covalent modification marks operate over distinct timescales constituting a multilayered molecular system that enables Pol II to encode the recent transcriptional and signaling history in a temporally ordered manner through distinct kinetic activities. In fact, the PTMs involved in regulatory transcriptional processes exhibit distinct lifetimes, forming fast (~5 seconds– minutes), intermediate (~1–30 min) and slow (~15 min – ~24h) biochemical transient processes. Rapid phosphorylation cycling supports second-to-minute responsiveness, whereas intermediate marks such as proline isomerization and O-GlcNAcylation integrate signals over longer windows and slower ubiquitination-linked processes can encode more persistent regulatory history through polymerase recycling and turnover. This kinetic-layer analysis yields a conservative cell-level estimate of ~1.15 MB of kinetically addressable regulatory state capacity, distinct from the theoretical maximum.

Finally, we addressed how non-uniform state probabilities and coupling reduce achieved entropy relative to state-space capacity (Supplementary Information S2). We also performed sensitivity analyses (Figure 3) showing that the conclusion of high-dimensional PTM pattern capacity is robust to plausible alternative discrete-state levels for specific PTM classes (Supplementary Information S3). A more precise estimation of achieved entropy would require (i) condition-resolved occupancy distributions, (ii) joint correlations among sites on the same Pol II molecule, and (iii) sufficient single-molecule resolution to distinguish combinatorial patterns rather than population averages. These requirements remain experimentally challenging; therefore, a more precise entropy estimation of Pol II is outside the scope of the present work. All numerical results shown here are fully reproducible using the “Python Reproducibility Modules” provided in Supplementary Information S4.

**Figure 3.**
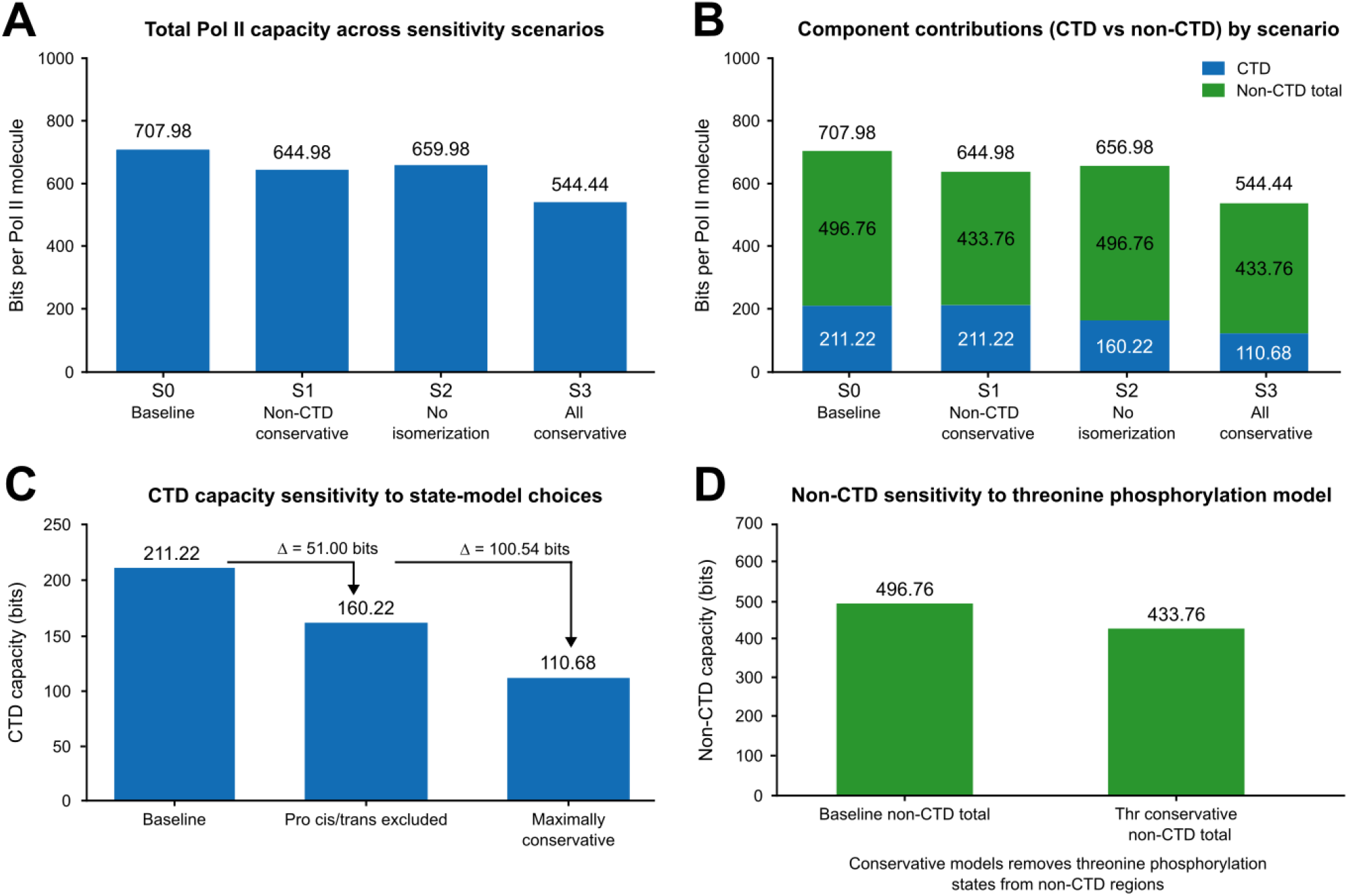
Sensitivity analysis of Pol II PTM information-capacity estimates. A, Total Pol II theoretical information capacity across four sensitivity scenarios. The baseline model (S0) yields 707.98 bits per Pol II molecule. A non-CTD conservative model removing threonine phosphorylation states from non-CTD regions (S1) yields 644.98 bits. A CTD model excluding Pro cis/trans isomerization (S2) yields 656.98 bits. The fully conservative model combining non-CTD threonine-phosphorylation removal with maximal CTD conservative assumptions (S3) yields 544.44 bits. B, Component contributions of CTD and non-CTD regions across the same scenarios. CTD contributions are shown separately from the combined non-CTD total, corresponding to Rpb1 core non-CTD plus Rpb2–Rpb12. C, Sensitivity of CTD capacity to alternative state-model choices. Excluding Pro cis/trans isomerization reduces CTD capacity from 211.22 to 160.22 bits, a reduction of 51.00 bits. The maximally conservative CTD model reduces CTD capacity to 110.68 bits, a reduction of 100.54 bits relative to baseline. D, Sensitivity of the non-CTD capacity to a conservative threonine-phosphorylation model. Removing threonine phosphorylation states from non-CTD regions reduces the combined non-CTD capacity from 496.76 to 433.76 bits. Together, these sensitivity analyses show that the conclusion of a high-dimensional Pol II PTM information landscape remains robust under conservative alternative state-model assumptions.

Overall, this quantification, grounded in Shannon’s information theory, reframes Pol II PTM landscapes as a distributed, dynamically reconfigurable regulatory layer whose capacity can be estimated transparently and refined as richer occupancy and correlation data become available. Similarly, our study highlights the quantitative scale of the functional coupling between genetic information and cellular physiological processes. In a broader sense, casting PTM landscapes in information-theoretic terms provides an analytical language to connect biochemical complexity with regulatory potential, and offers a quantitative basis for considering how dysregulation of PTM writers, erasers, and readers could alter transcriptional control and contribute to profound cellular consequences.

## Acknowledgments

The author I.M. was supported by Basque Government funding, grant GIC21/04, and by the Spanish Government, grants PID2024-156800OB-I00 and PID2024-156173OA-I00. JMC acknowledges financial support from Ikerbasque: The Basque Foundation for Science, and from Spanish Ministry of Science (PID 2023-148012OB), Spanish Ministry of Health (PI22/01118), and Basque Ministry of Health (2023111002 & 2022111031).

## Author contributions

IMDF: conceived, designed and directed the investigation, writing – original draft. MF, IM, LM, JC-P, BC-P, CB, GPY, JMC and JIL: investigation, research mapping, writing – review and editing. JC-P and BC-P Python Modules. All authors wrote the manuscript and agreed with its submission.

## Data availability

The version-pinned UniProt sequences used for residue mapping are provided as Supplementary Data File 3 (uniprot_polII_seqs_2026_05_22.tsv). The integrated RNA Polymerase II PTM atlas used in this study is provided as Supplementary Data File 4 (ptms_polII_2026_05_22.tsv). Both supplementary data files are also deposited at https://doi.org/10.5281/zenodo.20345702. All source data were retrieved from six publicly available databases: UniProt UniSave (https://www.ebi.ac.uk/uniprot/unisave/app/#/), UniProtKB (https://www.uniprot.org/uniprotkb/), EBI Proteins (https://www.ebi.ac.uk/proteins/api/), dbPTM (https://biomics.lab.nycu.edu.tw/dbPTM/), iPTMnet (https://research.bioinformatics.udel.edu/iptmnet/), PhosphoSitePlus (https://www.phosphosite.org/), and GlyGen (https://glygen.org/). The estimate of 80,200 RNA Polymerase II copies per mammalian nucleus was taken from BioNumbers (ID 112321; Zhao et al. 2014^63^; https://bionumbers.hms.harvard.edu/bionumber.aspx?s=n&v=0&id=112321). Literature-mined PTM entries were incorporated from Harding et al. (1992)^59^, Noble et al. (2005)^60^, Xiang et al. (2010)^56^, Werner-Allen et al. (2010)^47^, Zhang et al. (2012)^48^, Mayfield et al. (2015)^57^, Lewis et al. (2016)^49^, Gibbs et al. (2017)^58^ and Sharma et al. (2019)^61^.

## Code availability

The Jupyter notebooks required to fully reproduce all tables and numeric results from these source data are described in Supplementary Information S4 and provided as Supplementary Data File 1 and Supplementary Data File 2, and have also been deposited on GitHub (https://github.com/JoseCarrascoPujante/Multi-timescale_PTM_architecture_in_human_RNA_polymerase_II) and Zenodo (https://doi.org/10.5281/zenodo.20345702).

## STAR★Methods

We quantified the information capacity encoded by PTM patterns in human Pol II using an explicit discrete-state model at the residue level and computing Shannon maximal entropy under a uniform, independent-site assumption. We mainly report (I) a theoretical upper bound based on the full PTM state space and (II) a physiological/kinetically accessible estimate derived from coarse-graining PTM dynamics into multi-timescale layers.

### Pol II subunit definitions and sequence sources

In our study, we considered the canonical 12-subunit human complex of Pol II (Rpb1–Rpb12; gene products POLR2A–POLR2L and associated subunits). Protein sequences, isoform identifiers, and residue coordinates were obtained from UniProt entries corresponding to the reviewed human proteins. Residue numbering was standardized to UniProt canonical sequences. For Rpb1, the CTD region and repeat numbering were defined using the annotated human CTD architecture (UniProt P24928^37^). PTM sites were compiled from curated PTM resources (UniProt, dbPTM, iPTMnet, PhosphoSitePlus and GlyGen; supplementary cross-checking with UniProt feature annotations and PhosphoSitePlus where relevant). For each subunit, all the supported PTM sites were extracted with residue identity, position, and PTM type.

### Input data

PTM sites were imported from the free access resources of said online PTM databases considering their residue position, residue identity, PTM type(s), and UniProt accession. Residue coordinates were standardized to UniProt canonical sequences; for Rpb1, the CTD region and repeat boundaries were defined using the annotated POLR2A CTD architecture (52 heptads, UniProt P24928^37^).

### Data cleaning and normalization

PTM entries for each subunit were mapped to a canonical site key from the UniProt protein sequence (protein accession, residue letter, residue index). Duplicate entries across sources (same site and PTM type) were merged. For residues annotated with multiple PTM types, all PTM types were retained for that site. Ambiguous entries (e.g., unspecified residue within a peptide) and sites not mappable to the canonical sequence were excluded. CTD (Rpb1) heptad-repeat sites were modeled explicitly using mammal-relevant PTM classes (Figure 1). PTMs on residues outside the CTD were included in the Rpb1 core non-CTD set to avoid double counting.

### Discrete PTM state models (per residue)

Each modifiable residue *i* was assigned a discrete number of mutually exclusive states *s*_i_ reflecting the PTM chemistry used in the model. Unless otherwise stated, “unmodified” was treated as one state, and modified forms were treated as additional mutually exclusive states.

### CTD PTM model

For CTD heptads, residues were assigned:

- Tyr1: unmodified/phosphorylated (22 sites)
- Ser2: unmodified/phosphorylated (27 sites)
- Ser2: unmodified/O-GlcNAcylated/phosphorylated (3 sites)
- Ser4: unmodified/phosphorylated (3 sites)
- Ser5: unmodified/phosphorylated (36 sites)
- Ser5: unmodified/O-GlcNAcylated/phosphorylated (4 sites)
- Ser7 (1621): unmodified/O-GlcNAcylated/phosphorylated (1 site)
- Ser7: unmodified/phosphorylated (4 sites)
- Ser9 (heptad 49) (1938): unmodified/phosphorylated (1 site)
- Pro3 (YSPTSPS, YSPSSPS & YSPTSPN): cis/trans isomerization (26 sites)
- Pro6 (YSPTSPS & YSPTSPN): cis/trans isomerization (25 sites)
- Thr2: unmodified/phosphorylated (5 sites)
- Thr4: unmodified/O-GlcNAcylated/phosphorylated (2 sites)
- Thr4: unmodified/phosphorylated (15 sites)
- Thr5 (1885): unmodified/phosphorylated (1 site)
- Thr7: unmodified/phosphorylated (5 sites)
- Thr7 (1894): unmodified/O-GlcNAcylated/phosphorylated (1 site)
- Thr10 (heptad 49) (1939): unmodified/phosphorylated (1 site)
- Lys7: unmodified/acetylated/dimethylated/monomethylated (5 sites)
- Lys7: unmodified/acetylated/dimethylated/monomethylated/ SUMOylated/trimethylated/ubiquitinated (2 sites)
- Lys7 (1908): unmodified/acetylated/dimethylated (1 site)
- Arg4 (1603): unmodified/monomethylated (1 site)
- Arg7 (1810): unmodified/asymmetrically, dimethylated/citrullinated/dimethylated/monomethylated/symmetrically demethylated (1 site)
- Ser2, Ser6 C-terminal tail: unmodified/phosphorylated (2 sites)

The CTD was modeled as 21 canonical repeats with consensus **YSPTSPS** and 31 non-consensus repeats. Residues present in the CTD sequence but not assigned a curated PTM state in this model were treated as having s_i_ = 1 and therefore contributed 0 bits to the information capacity.

### Non-CTD PTM model (Rpb1 core and Rpb2–Rpb12)

Outside the CTD, PTMs were modeled using the following state sets:

- Acetylation: unmodified / acetylated (s = 2).
- Glutathionylation: unmodified / glutathionylated (s = 2).
- Methylation: unmodified / methylated (s = 2).
- O-GlcNAcylation: unmodified / O-GlcNAcylated (s = 2).
- Phosphorylation: unmodified / phosphorylated (s =2).
- Sulfoxidation: unmodified / sulfoxidized (s = 2).
- SUMOylation: unmodified / SUMOylated (s = 2).
- Ubiquitination: unmodified / ubiquitinated (s = 2).

### Multi-PTM residues and composite state counts

If a residue was annotated with more than one PTM type (e.g., a serine that can be phosphorylated or O-GlcNAcylated), we treated PTM types as **mutually exclusive** at that residue and defined an effective composite state count:

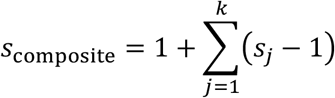

where *s*_*j*_ is the state count for PTM type j acting at that residue and the “1” represents the unmodified state shared across PTM types.

This choice avoids double-counting the unmodified state and should be interpreted as a conservative residue-level state model, not as a claim that simultaneous multi-mark occupancy cannot occur in all biological contexts.

### Information calculations

Per-site information was computed as *I*_*i*_ = log_2_(*s*_*i*_). For a set of *n* modeled sites:

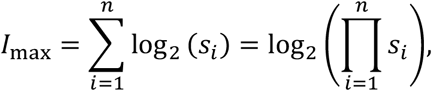

and the pattern-space size is 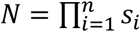 with *I*_max_ = log_2_(*N*). Total Pol II capacity was computed as:

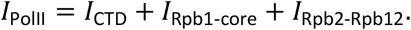

Cellular capacity was computed as *I*_cell_ = *M* ⋅ *I*_PolII_, with *M* = 80,200 Pol II molecules per nucleus.

### Physiological/kinetically accessible estimate

To estimate a conservative physiologically accessible information capacity, PTMs were coarse-grained into three explicitly parameterized kinetic layers: a fast CTD phosphorylation layer, an intermediate Pro6 isomerization/O-GlcNAcylation layer, and a slow ubiquitination-linked layer. Acetylation, SUMOylation, and methylation were acknowledged qualitatively but not explicitly parameterized because current condition-resolved kinetic occupancy data are insufficient to define a defensible effective-state model.This physiological information bound can be computed as:

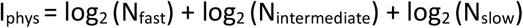

For each layer k, N_k_ = a_k_^m_k_ where m_k_ is the effective number of sites/processes and a_k_ is the effective number of kinetically distinguishable states per site/process. Therefore I_k_ = log_2_ (N_k_) = m_k_ log_2_ (a_k_)

### Reproducibility

Supplementary Information S4 provides a Python implementation that reproduces all numerical calculations and tables. The pipeline requires Python ≥3.10 together with pandas, requests, IPython/Jupyter utilities, and standard-library modules.

### Limitations

The reported maximum theoretical capacities are upper bounds derived from a discrete-state model under independence and equal-probability assumptions. In vivo, many marks are rare, transient, or stage-restricted, and sites are not independent due to enzymatic coupling, reader-mediated cooperativity, steric/structural constraints, and transcription-cycle ordering, all of which reduce achieved entropy relative to the theoretical maximum (Supplementary Information S2). PTM catalogs are incomplete and biased toward abundant proteins and detectable chemistries, so the enumerated state space may omit true sites or include sites with negligible occupancy. Partial observability is also a major limitation: many measurements are population-averaged and do not resolve correlated site occupancies on individual Pol II molecules. More kinetic lifetime measurements and single-molecule PTM occupancy data will be required to estimate achieved entropy more precisely.

## Supplementary Information S1. Pol II architecture

Human RNA polymerase II is organized as a 12-subunit complex, Rpb1–Rpb12, built around a catalytic core whose structural and mechanistic principles are conserved from archaea to humans^54^. The largest subunit, Rpb1, carries the extended C-terminal domain (CTD), a repetitive regulatory platform that is densely targeted by reversible PTMs. These CTD modifications contribute to the temporal coordination of Pol II activity across the transcription cycle, from initiation and promoter escape to productive elongation, termination, and co-transcriptional RNA processing^44^. Structural and functional studies of the Pol II pre-initiation complex (PIC) show that general transcription factors bind Pol II surfaces including Rpb2 domains (especially the protrusion and lobe domains) and the core initiation module which includes the Rpb3/Rpb11 heterodimer^77,78^. The Rpb4/Rpb7 heterodimer forms a specialized RNA-binding module^79,80^ whose phosphorylation appears to regulate its engagement with RNA polymerase II^81–83^, linking elongation and termination^81,84^ to mRNA imprinting^81,83^ and decay^80,83^. Rpb5 stability and its assembly into RNA Pol II, which influences nuclear Pol II levels^85^, is promoted by URI^86^, a transcriptional repressor^87,88^ whose multiple phosphorylation and acetylation states position it as a potential regulatory hub translating extracellular signalling into specific patterns of RNA Pol II transcriptional activity^86^. Separately, phosphorylation of the Rpb6 N-terminal domain has been postulated to confer it with regulatory functions^89^.

The small subunits Rpb8–Rpb12 also participate in PTM-dependent processes essential for Pol II function. It has been suggested that phosphorylation of the small subunits Rpb8, Rpb9, Rpb10 and Rpb12 may influence transcriptional activity^90^. Notably, Rpb9 has documented roles in initiation-factor interactions^91^, transcriptional fidelity^92,93^, transcription elongation rate^94^ and transcription-coupled DNA repair^95^. In the cellular response to UV-induced DNA damage, non-proteolytic polyubiquitination of Rpb8 approximately doubles cell survival rates^96^, while Rpb9-promoted ubiquitination and proteasomal degradation of subunit Rpb1 disassembles RNA Pol II molecules arrested at DNA lesions^97^. Rpb11 and Rpb12 are small Pol II subunits associated with the Rpb3-containing assembly subcomplex^98–100^. Rpb11 supports Pol II assembly through its heterodimerization with Rpb3^98,101^ and participates in the Mediator-bound preinitiation complex^78^. It also carries four acetylation sites, two of which have been predicted to stabilize the Rpb3/Rpb11 interface^102^. In turn, Rpb12 contributes to Pol II biogenesis^99^ and, based on ortholog evidence from *Archaea*, also fosters promoter opening^103^. It carries surface-exposed phosphosites with the potential to influence its interactions with transcription regulators^90^. Collectively, PTMs across the entire Pol II subunit ensemble (Rpb1–Rpb12), with over 630 amino acid residues annotated as undergoing over 775 covalent modifications (Reproducibility Module 17, Supplementary Information S4), are centered on Rpb1, whose CTD PTMs connect RNA Pol II to co-transcriptional RNA processing, termination and transcription-coupled DNA repair^44,104,105^. Beyond Rpb1, phosphorylation of the Rpb4/7 heterodimer modulates stalk stoichiometry and links transcription termination to mRNA imprinting^80,81^, and Rpb8 ubiquitination regulates the DNA-damage response^96^. For the remaining subunits, evidence is still dominated by PTM mapping and indirect pathway effects, leaving RNA Pol II full-ensemble coordination unresolved^106,107^.

## Supplementary Information S2. From state-space capacity to achieved entropy under non-uniform and correlated PTM probabilities

### S2.1. Purpose and interpretation

Our first results report a state-space capacity upper bound for Pol II PTM patterns under a discrete-state model in which each site *i* has *s*_*i*_ mutually exclusive states. Under site independence and uniform state probabilities, the maximal information capacity is:

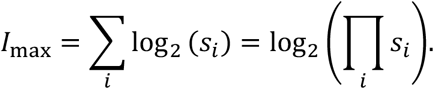

This quantity equals the Shannon entropy only when state probabilities are uniform at each site and sites are independent. The purpose of this Supplementary Information is to make this distinction explicit and to show, with simple examples, how non-uniform state probabilities and site coupling reduce achieved entropy relative to maximal state-space capacity.

Accordingly, throughout the manuscript, capacity refers to the logarithm of the modeled state-space size, whereas ‘achieved entropy’ refers to the entropy of an empirical occupancy distribution

### S2.2. Site-wise entropy under non-uniform occupancies

If site *i* has state probabilities 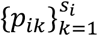, the achieved (site-wise) entropy is:

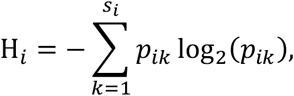

and under independence the achieved entropy is H = ∑_i_ H_i_. Because biological PTM occupancies are typically sparse, stage-dependent, and often dominated by the unmodified state, H ≪ I_max_ in most regimes.

To show how non-uniform priors reduce achieved entropy, we consider simple distributions in which the unmodified state dominates. These distributions serve to demonstrate both scale and directionality.

#### S2.2.1. Binary sites (2 states): unmodified vs modified

For a binary PTM site with states unmodified and modified, let p be the probability that the site is modified. The state probabilities are therefore 1 − p and p. Then, the achieved entropy is:

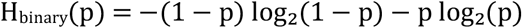

Examples are shown in Table S2.1.

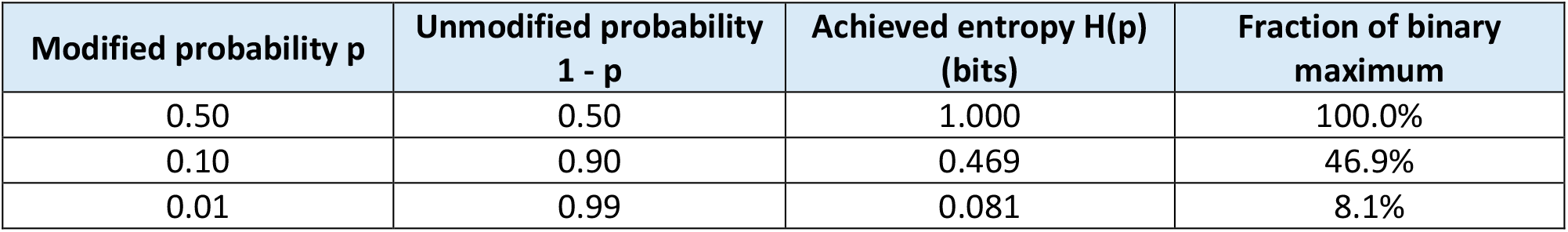

Even for binary PTMs, sparse occupancy can reduce effective bits per site by an order of magnitude relative to the maximum.

#### S2.2.2. Three-state sites (3 states): unmodified / phospho / O-GlcNAcylated

For a three-state site such as a serine or threonine modeled as unmodified, phosphorylated, or O-GlcNAcylated, the maximal state-space capacity is log_2_(3) = 1.585 bits. However, a non-uniform occupancy distribution gives a smaller achieved entropy.

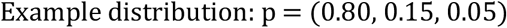

then:

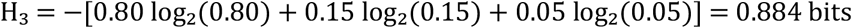

In this illustrative case, the achieved entropy is approximately 56% of the maximal three-state capacity. This example is not intended as a measured occupancy distribution; it simply demonstrates how non-uniform probabilities reduce entropy relative to state-space capacity.

#### S2.2.3. Four-state sites: arginine modification example

For a four-state site, such as an arginine modeled as unmodified, asymmetrically dimethylated, symmetrically dimethylated, or citrullinated, the maximal capacity is log_2_(4) = 2 bits. If the unmodified state dominates, the achieved entropy is again much lower.

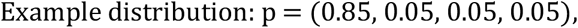

then:

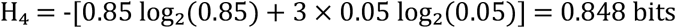

This illustrates that high nominal state granularity does not imply high achieved entropy unless the alternative states are substantially populated. The numerical occupancy examples above are illustrative and are not used to infer in vivo Pol II entropy.

### S2.3. State-space capacity versus achieved entropy

The residue-level capacities reported in the main text quantify the size of the modeled PTM state space under a uniform independent-site assumption. This is useful because it provides an auditable upper bound and allows direct comparison among CTD, Rpb1-core, and Rpb2–Rpb12 contributions.

However, achieved in vivo entropy would require condition-resolved probabilities for every PTM state at every site. If the sites were independent and if all marginal probabilities were known, the achieved entropy would be:

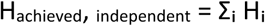

In practice, PTM state probabilities are likely to be non-uniform because many marks are rare, transient, condition-specific, or transcription-stage-specific. Therefore:

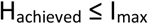

The inequality is expected to be strict for most biological contexts.

### S2.4. Reduction of entropy by site coupling and correlations

Pol II PTMs are not generally independent. Shared writers and erasers, CTD-reader recruitment, steric constraints, structural coupling, transcription-cycle ordering, and chromatin context can generate statistical dependence among sites.

For two sites X and Y, the joint entropy satisfies:

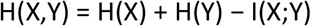

where I(X;Y) is the mutual information between the two sites. If the two sites are independent, I(X;Y) = 0. If they are coupled, I(X;Y) > 0 and the joint entropy is lower than the sum of the marginal entropies. For more than two sites, it is generally not sufficient to subtract a simple sum of pairwise mutual information terms, because higher-order dependencies may exist. The safe general expression is:

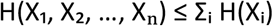

The difference between the sum of marginal entropies and the joint entropy is the total correlation, also called multi-information:

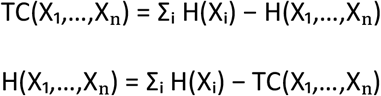

Total correlation is non-negative and captures the aggregate dependence among sites, including higher-order dependencies that are not reducible to pairwise mutual information.

### S2.5. Scope and data requirements for estimating achieved entropy

The examples above show that achieved entropy can be smaller than state-space capacity when the unmodified state is dominant or when specific modified states are rarely attained. A precise estimation of achieved in vivo entropy would require condition-resolved occupancy distributions and joint correlations among sites on the same Pol II molecule, and, for information transfer to transcriptional output, coupling between PTM patterns and transcriptional states.

These requirements remain experimentally challenging; therefore, more precise entropy estimation of Pol II, at present, is outside the scope of our work. Instead, the manuscript reports theoretical state-space capacity and a conservative kinetically addressable estimate.

A practical way to express the impact of coupling is to define an effective number of independently addressable variables. If a group of sites behaves as a coupled module (e.g., CTD phosphorylation patterns constrained by kinase/phosphatase dynamics and repeat-level coupling), then the achieved entropy scales with the number of effectively independent degrees of freedom, not with the raw number of chemically modifiable residues. Thus, this Supplementary Information defines the experimental data required for future conversion of the present capacity framework into condition-specific entropy estimates.

## Supplementary Information S3. Sensitivity analyses of Pol II PTM information-capacity estimates

Here, we evaluate how the estimated **maximum information capacity** of Pol II PTM patterns depends on discrete-state modeling choices. The main text reports an **upper-bound capacity** under a residue-level discrete-state model with mutually exclusive states at each site.

For each site *i* with *s*_*i*_ mutually exclusive states, the maximal capacity is:

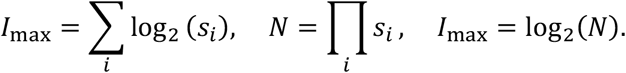

When multiple PTM types are annotated on the same residue, PTMs are treated as mutually exclusive and combined using:

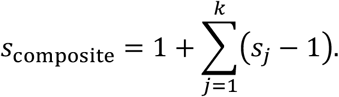

These sensitivity analyses quantify how I_max_ changes under alternative plausible assignments of s_i_ for PTM classes.

### S3.1. Alternative state models tested

A. CTD Proline cis / trans isomerization Baseline: *s*_Pro_ = 2 (cis / trans isomerization) Conservative: *s*_Pro_ = 1 (unmodified)
B. CTD Arginine state granularity Baseline: site-specific Arg state models as shown in Table 2, with different sets of methylation states for each Arg residue, ranging from s=2 to s=6 Intermediate: *s*_Arg_ = 2 (unmodified / modified) Conservative: *s*_Arg_ = 1 (unmodified) The intermediate Arg/Lys models are included to define possible alternative granularities but are not used as separate rows in the representative S0–S3 scenario table.
C. CTD Lysine state granularity Baseline: site-specific Lys7 state models as used in Table 2, with individual Lys7 residues assigned to curated acetylation/methylation state sets ranging from s=4 to s=7 rather than to a universal Lys7 state model. Intermediate: *s*_Lys_ = 2 (unmodified/modified) Conservative: *s*_Lys_ = 1 (unmodified)
D. Threonine phosphorylation Baseline: threonine sites were modeled according to their curated state sets, including unmodified/phosphorylated or unmodified/O-GlcNAcylated/phosphorylated where applicable. Conservative Thr model: threonine phosphorylation states were removed from the model; threonine sites lacking other PTM states were assigned s_i_ = 1 and therefore contributed 0 bits.

### S3.2. Sensitivity of Rpb2–Rpb12 capacity (based on Supplementary Tables S2– S12)

Using Supplementary Tables S2–S12, and after consolidating multi-PTM residues into composite sites, the maximal information capacity for Rpb2–Rpb12 under the baseline model is:

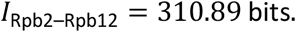

We then recompute *I*_Rpb2–Rpb12_ under the alternative Thr granularity state model while leaving all other PTMs unchanged:

Conservative (Thr = 1)

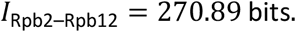

Therefore, even in the absence of Thr phosphorylation, the Rpb2–Rpb12 subsystem retains a capacity in the hundreds-of-bits range.

### S3.3. Sensitivity of Rpb1 core (non-CTD; Supplementary Table S1)

For the non-CTD Rpb1 core (159 sites), the baseline model yields:

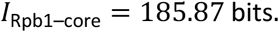

Under alternative non-CTD state models:

Conservative (Thr *s* = 1):

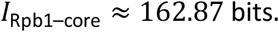

### S3.4. Sensitivity of CTD capacity to state definitions

Baseline CTD capacity (main text):

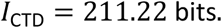

#### S3.4.1 CTD without isomerization in Pro sites

Replacing Pro cis/trans from *s* = 2 to *s* = 1 (and keeping all other CTD assumptions unchanged):

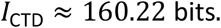

Thus, the explicit 2-state Pro cis/trans model contributes approximately:

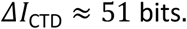

#### S3.4.2. CTD “maximally conservative” illustrative variant

If (i) Prolines are considered as unmodified (s=1), (ii) Thr phosphorylation is removed (s=1), (iii) Lys7 PTM states are collapsed to the unmodified state (s=1), and (iv) Arg residues are pooled as s=1 (unmodified), then:

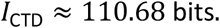

### S3.5. Scenario table: combined sensitivity for total Pol II capacity

The total Pol II capacity is:

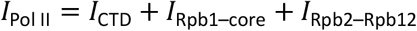

Assuming *M* = 80,200 Pol II molecules per nucleus:

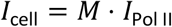

## Supplementary Information S3-Table S1. Representative sensitivity scenarios for total Pol II maximal information capacity

| Scenario | CTD state model | non-CTD state model (Rpb1-core + Rpb2-Rpb12) | $I_{\text{Pol II}}$ (bits per molecule) | Bytes per molecule ($I \div 8$) | $I_{\text{cell}}$ (bits per nucleus) | Bytes per nucleus ($I \div 8$) | MB per nucleus ($10^6$ bytes) |
| --- | --- | --- | --- | --- | --- | --- | --- |
| **S0 (baseline)** | Baseline | Baseline | 707.98 | 88.50 | $5.68 \times 10^7$ | $7.10 \times 10^6$ | 7.10 |
| **S1 (non-CTD conservative)** | Baseline | Thr conservative | 644.98 | 80.62 | $5.17 \times 10^7$ | $6.47 \times 10^6$ | 6.47 |
| **S2 (no isomerization)** | Pro cis/trans excluded | Baseline | 656.98 | 82.12 | $5.27 \times 10^7$ | $6.59 \times 10^6$ | 6.59 |
| **S3 (all conservative)** | Maximally conservative | Thr conservative | 544.44 | 68.05 | $4.37 \times 10^7$ | $5.46 \times 10^6$ | 5.46 |

These scenario definitions correspond to the sections of Supplementary Information S3.1–S3.4. $I_{\text{Pol II}} = I_{\text{CTD}} + I_{\text{Rpb1-core}} + I_{\text{Rpb2-Rpb12}}$ . Cellular totals use $I_{\text{cell}} = M \cdot I_{\text{Pol II}}$ with $M = 80,200$ . MB is reported in decimal units (1 MB = $10^6$ bytes). This table and all calculations in the text can be regenerated using the Reproducibility Module 18 (Supplementary Information S4)

In brief, across plausible alternative state-model definitions, I_Pol II_ remains in the order of hundreds of bits per molecule, and the implied nuclear-level capacity remains multi-megabyte scale under the same independence/uniform assumptions. These sensitivity analyses therefore support the robustness of the qualitative conclusion (high-dimensional PTM pattern capacity), while clarifying the quantitative dependence on discrete-state granularity choices for specific PTM classes (see Supplementary Information S3-Table S1. Representative sensitivity scenarios for total Pol II maximal information capacity).

### S3.6. Link to Supplementary Information S2 (capacity vs achieved entropy)

The sensitivity analyses above quantify how the maximal capacity *I*_max_ = ∑_*i*_ log_2_ (*s*_*i*_) depends on discrete-state granularity choices. Importantly, these results concern state-space capacity under uniform, independent-site assumptions. As discussed in Supplementary Information S2, biological PTM occupancies are typically non-uniform and sites are often coupled, so the achieved entropy *H* is generally substantially smaller than *I*_max_ and further reduced by correlations. Accordingly, S3 evaluates robustness of the theoretical capacity estimate to alternative discrete-state definitions. By contrast, S2 addresses how non-uniform occupancies and site coupling reduce achieved entropy relative to this capacity. Thus, S2 and S3 address complementary sources of uncertainty: biological occupancy/correlation effects and discrete-state model granularity. The numerical occupancy examples are illustrative and are not used to infer in vivo Pol II entropy.

## Supplementary Information S4. Reproducibility package for Pol II PTM information-capacity calculations

### Overview

This supplement documents two interactive Python (Jupyter) notebooks: Notebook 1 (*notebook_1*.*ipynb*, Supplementary Data File 1) and Notebook 2 (*notebook_2*.*ipynb*, Supplementary Data File 2) that together reproduce every numeric value, table and figure reported in the main text and in Supplementary Information S1 and S3. Notebook 1 contains ten Reproducibility Modules (1–10) that retrieve the archival UniProt^37^ UniSave canonical sequences for human RNA Pol II subunits 1-12 and build the integrated post-translational modification (PTM) atlas of human RNA polymerase II from public databases and literature-mined entries, respectively saving these two databases as files *uniprot_polII_seqs_{CURRENT_DATE}*.*tsv* (Supplementary Data File 3) and *ptms_polII_{CURRENT_DATE}*.*tsv* (Supplementary Data File 4) in the notebook’s working directory. Notebook 2 contains eight Reproducibility Modules (11–18) that load the frozen PTM atlas and the version-pinned UniProt sequences and compute the manuscript’s information-theoretic results. Modules are intended to run sequentially within each notebook; their numbering matches the order of the calculations in the manuscript and in Supplementary Information S1 and S3.

For exact, network-independent reproducibility, the frozen archival UniProt UniSave sequences *uniprot_polII_seqs_2026_05_22*.*tsv* and PTM atlas *ptms_polII_2026_05_22*.*tsv* snapshots are bundled as Supplementary Data File 3 and Supplementary Data File 4. When both files are present in the working directory, every download module in Notebook 1 detects the cache and skips its network call, and Notebook 2 reads the snapshots directly. The Notebooks’ audit asserts are pinned to these snapshots and will fail against new ones regenerated from updated upstream databases.

The package regenerates: (1) the integrated PTM atlas and the version-pinned protein sequence reference (Notebook 1, Modules 1–10); (2) a colour-coded map of the 52 heptads and 10-residue tail of the Rpb1 CTD (Table 1; Module 11); (3) the multiplicative CTD state-space size *N_CTD*, the information capacity *I_CTD*, and the residue-class decomposition of *I_CTD* in Table 2 (Module 12); (5) the information capacity of the Rpb1 non-CTD core region (residues 1–1592) in Supplementary Table S1 (Module 13); (6) per-subunit capacities for Rpb2–Rpb12 in Supplementary Tables S2–S12 (Module 14); (7) the per-molecule and per-nucleus totals *I_total* and *I_nucleus* (Module 15); (8) the multi-timescale physiological capacity *I_physio* with its fast, intermediate and slow layers (Module 16); (9) total unique PTM sites and modified residues in the integrated atlas (Module 17); and (10) the four S0–S3 sensitivity scenarios (Supplementary Information S3.1–S3.4 and Supplementary Information S3-Table S1; Module 18).

### Notebook 1. PTM and protein sequence data retrieval

#### Environment check (optional)

Verifies that the execution environment meets the minimum software dependencies for this notebook (Python ≥ 3.10, pandas ≥ 1.3, requests ≥ 2.20, beautifulsoup4 ≥ 4.9, IPython ≥ 7.0) and aborts with a diagnostic message if any requirement is missing or below the required version, preventing execution errors downstream.

#### Reproducibility Module 1. Canonical UniProt sequence retrieval

Loads the 12 human Pol II protein sequences from a saved cache file (named *uniprot_polII_seqs_\**.*tsv*) if one exists; otherwise fetches them directly from the UniProt database using fixed, pinned version numbers for each protein. The specific versions are locked in to ensure the output is always identical no matter when or where the code is run. The sequences are stored in a simple lookup table (protein ID → sequence string) that is passed to Module 2 for residue-letter lookups and to Module 8, which breaks the main subunit (P24928) into 52 heptads plus 10-residue C-terminal tail. If no cache was found at startup, the freshly downloaded data is saved to a new cache file for next time.

#### Reproducibility Module 2. UniProt manually curated PTM annotations

Fetches manually reviewed modification records for each of the twelve Pol II subunits directly from UniProt, keeping only entries that point to a single, specific position in the protein. The modification name is cleaned up by dropping any extra detail after the first semicolon (e.g. “Phosphoserine; by kinase X” becomes just “Phosphoserine”), and the corresponding amino acid letter is looked up from the sequences loaded in Module 1. The output is a tidy table with one row per (Protein, Residue, Modification) combination — the shared format used by all six download modules.

#### Reproducibility Module 3. UniProt large-scale proteomics PTMs

Retrieves modification data assembled from large-scale mass-spectrometry experiments via the EBI Proteins API. Each entry reports a short peptide sequence and the position of the modified residue within it; the code uses those two pieces of information to figure out the residue letter and its position in the full protein. Results are emitted in the shared format.

#### Reproducibility Module 4. dbPTM experimental PTMs

Downloads all experiment archives from the dbPTM database, decompresses them and reads them into a standard six-column layout. Only rows belonging to the twelve Pol II accessions are kept. The modified residue letter is taken from the central position (index 10) of the 21-amino-acid flanking window surrounding each modification site. All individual per-archive tables are combined and de-duplicated into a single output in the shared format.

#### Reproducibility Module 5. iPTMnet integrated PTMs

Loads the iPTMnet integrated scoring file directly as a tab-separated table, keeping only the Substrate, Residue and PTM columns. Rows are filtered to the twelve canonical Pol II accessions plus the alias ‘P24928-1’ that iPTMnet stores separately despite sequence identity to canonical P24928 — the suffix is stripped so these records merge cleanly with the other sources downstream.

#### Reproducibility Module 6. PhosphoSitePlus PTMs

Because PhosphoSitePlus does not offer a free data API, this module scrapes the information directly from its web pages. A built-in lookup table maps each Pol II UniProt accession to its internal PhosphoSitePlus identifier. For each subunit, the module first reads the abbreviation glossary from the protein overview page (e.g. “p” → “Phosphorylation”), then works through the site table page and records every human modification site that is not flagged as belonging to an isoform.

#### Reproducibility Module 7. GlyGen glycosylation and phosphorylation annotations

Queries the GlyGen protein database using “{*UniProtKB accession number*}-1” identifiers for each Pol II subunit studied, since that is how GlyGen indexes its entries for those UniProtKB canonical protein isoforms. The response is processed in two passes: first collecting glycosylation entries with a meaningful subtype (such as O-GlcNAcylation), then collecting phosphorylation entries. Three-letter amino acid codes returned by the API are converted to single-letter codes, and the “-1” isoform suffix is stripped from the accession before passing results downstream.

#### Reproducibility Module 8. CTD heptad position map

Cuts out the CTD region of the main Pol II subunit (Rpb1, residues 1593–1970) and divides the heptad-bearing stretch into repeating units by splitting on every tyrosine (every CTD heptad starts with a tyrosine residue), producing 52 CTD heptapeptides and a lookup table mapping each residue position to its heptad number and position within that heptad for use in Modules 9 and 10. The remaining 10 residues form the C-terminal tail. Two built-in checks confirm exactly 52 heptads covering exactly 368 residues, catching any accidental changes to the boundary constants. The module also defines two lookup tables used by later modules: one converting single-letter to three-letter amino acid codes, and one translating modification names into descriptive adjectives (e.g. “Phosphorylation” → “phosphorylated”) for use in human-readable labels in Module 10.

#### Reproducibility Module 9. Cross-database alignment and literature-mined additions

Takes the six harmonised tables from Modules 2–7 and adds a seventh table of modifications not covered by any of those databases, drawn from the literature: proline isomerization at positions 3 and 6 of heptads matching specific sequences (26 + 25 = 51 sites); O-GlcNAcylation at S1829 and S1896; and citrullination at R1810. A built-in check confirms the literature table contains exactly 54 entries. Each source is then collapsed so there is one modification string per (Substrate, Residue) pair, and all seven sources are horizontally joined into a wide audit table (*ptm_comparison*) with one column per data source, so every site reported by any database is visible in one place. The module also counts how many residues carry each unique combination of modifications, for downstream auditing.

#### Reproducibility Module 10. PTM data curation and integration

Standardises the modification vocabulary across all sources by collapsing synonyms and inconsistent naming into a single controlled set (e.g. “Ubiquitinylation” and “Ubiquitylation” both become “Ubiquitination”; various phosphoresidue names all become “Phosphorylation”). Two irreversible modification types that do not fit the reversible site model underlying the information-capacity calculation — N-Glycosylation and Caspase cleavage — are removed. Where both a specific O-GlcNAcylation and a generic O-Glycosylation call appear at the same residue, the generic one is absorbed into the specific one to avoid double-counting. The cleaned records are combined and de-duplicated into *all_databases_merge*, the single master table used by all downstream modules. Finally, the master table is written to disk as ptms_polII_{CURRENT DATE}.tsv; on future runs all download and curation steps detect this file and skip all network access entirely, preserving reproducibility without re-querying any database.

### Notebook 2. Information-theoretic calculations

#### Environment check and Analysis setup (required)

The environment check confirms that the required software versions are present and halts with a clear message if anything is missing. The analysis setup loads the most recent sequence cache and PTM master table from disk, reconstructs the 52-heptad position map for Rpb1 using the same boundary constants as Module 8 in Notebook 1, creates the *ctd_subclass* dictionary, and builds the amino acid and modification name lookup tables in the same way as Module 8 in Notebook 1. It then splits *all_databases_merge* into three non-overlapping pandas dataframes used by all downstream modules: *ctd_df* for Rpb1 CTD (positions 1593–1970), *core_df* for Rpb1 non-CTD core (positions 1–1592) and *r212_df* for the eleven non-Rpb1 subunits. It also defines the shared helper function *_I* that computes information capacity in bits as a sum of log2(*number of modification states + 1*), where +1 accounts for the baseline unmodified state, over all modified residues. Running this cell first is mandatory, as everything else in Notebook 2 depends on it.

#### Reproducibility Module 11. Colour-coded CTD heptad map (Table 1)

Produces Table 1 from the main text: an HTML rendering of all 52 CTD heptapeptides plus the 10-residue C-terminal tail, with each residue coloured according to which PTMs have been reported on it. The colour palette is built automatically from the combinations observed in the data, and a legend is included. Two non-standard CTD heptads carrying extra residues are marked with an asterisk.

#### Reproducibility Module 12. CTD combinatorial state space, information capacity, and Table 2 of the manuscript

Calculates how many distinct modifications states the CTD can theoretically encode by multiplying together the number of possible states at each site (including the unmodified state), then takes the log_2_ of that product to express the result in bits. Both numbers are checked against the values reported in the manuscript (log_10_ of *N_CTD* = 63.58; *N_CTD* = 211.22 bits) and will raise an error if recalculated from a different data snapshot. The module then produces Table 2, breaking the total capacity down by residue class and modification type, with the heptad region’s residues listed first in biological order and the C-terminal tail appended at the end. A final check confirms the per-row bit contributions sum exactly to *I_CTD*.

#### Reproducibility Module 13. Information capacity of the Rpb1 non-CTD core (Supplementary Table S1)

Applies the same capacity calculation to the non-CTD (core) portion of Rpb1 (residues 1–1592) and renders Supplementary Table S1, with one row per modified residue sorted by position. Columns show the residue, PTM type(s), UniProt accession, number of possible states, and bits per site. Summary counts of unique PTM types, modified residues, and (residue, PTM type) pairs are reported, with a check that the total calculated information capacity matches the manuscript value and the per-site bits sum to the same correct total.

#### Reproducibility Module 14. Information capacity encoded by PTMs in subunits Rpb2–Rpb12 (Supplementary Tables S2–S12)

Repeats the same procedure for the eleven non-Rpb1 subunits and produces one supplementary table per subunit in canonical order (Rpb2 through Rpb12). Each table lists every modified residue, its PTM combination, its UniProt accession, its per-site number of possible states, and its per-site bit contribution. Per-subunit capacities are printed and audited against their respective values reported in the manuscript, and final checks confirm that the computed total information capacity for Rpb1-Rpb12 matches both the manuscript value and the sum of the per-site bit contribution column across all tables.

#### Reproducibility Module 15. Per-molecule and per-nucleus information capacity

Adds the three compartment capacities (CTD + Rpb1 core + Rpb2–12) to get the total per-molecule *I_total*, then scales up by *N_POLII* = 80,200 — the estimated number of Pol II complexes in a mammalian nucleus (BioNumbers ID 112321; Zhao et al., 2014)^63^ — to get an upper-bound nuclear capacity. Results are reported in bits, bytes, and megabytes. A built-in check guards computed per-molecule and per-nucleus values against drifting from the manuscript value.

#### Reproducibility Module 16. Multi-timescale PTM biochemical architecture

Divides the per-molecule capacity into three layers defined by their characteristic regulatory timescales: a fast layer of 30 phosphorylation sites cycling on a ~5 s–15 min timescale with 5 effective states each; an intermediate layer of 32 proline isomerization and O-GlcNAcylation sites cycling over ~1–30 min with 2 states each; and a slow layer of 10 ubiquitination-related sites turning over on ~15 min–~24 h with 2.5 effective states each. The three-layer capacities are summed into a physiological per-molecule total, also reported at nuclear scale and compared to the theoretical upper bound. Checks confirm the physiological information capacity per molecule and per nucleus match manuscript values.

#### Reproducibility Module 17. Total unique PTM sites and modified residues (Supplementary Information S1)

Reports the total number of unique modified residues, counted as distinct (Accession, Residue) pairs, and the total number of unique (Accession, Residue, PTM) triplets across the entire integrated PTM atlas. These two scalars are then checked against the totals quoted in Supplementary Information S1.

#### Reproducibility Module 18. Sensitivity analyses (Supplementary Information S3 and Supplementary Information S3-Table S1)

Tests how sensitive the main results are to modelling assumptions by recalculating total Pol II information capacity under four scenarios: the manuscript baseline (S0); a conservative version that removes threonine phosphorylation outside the CTD (S1); a version that keeps the non-CTD model but removes proline isomerization from the CTD (S2); and a maximally conservative version that removes proline isomerization, threonine phosphorylation everywhere, and all lysine-7 and arginine modifications in the CTD (S3). Per-molecule and per-nucleus information capacities are reported for each scenario and assembled into Supplementary Table S3. Checks pin each scenario total to its manuscript value.

## Supplementary Tables S1–S12, note

PTM sites were compiled from UniProt, dbPTM, iPTMnet, PhosphoSitePlus and GlyGen. For each residue, we assign a discrete state count *s*_*i*_, with s=2 for every PTM type (phosphorylation *s*=2, acetylation *s*=2, etc.). When multiple PTM types map to the same residue, PTMs are treated as mutually exclusive and combined using: 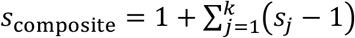. Bits per site are *log*_2_(*s*_*i*_).

**Supplementary Table S1.** POLR2A (RPB1 core, non-CTD)

| Residue | PTM Type(s) | UniProt accession | States ($s_i$) | Bits per site $\log_2(s_i)$ |
| --- | --- | --- | --- | --- |
| **M1** | Acetylation | P24928 | 2 | 1.00 |
| **S8** | Phosphorylation | P24928 | 2 | 1.00 |
| **S27** | Phosphorylation | P24928 | 2 | 1.00 |
| **K32** | SUMOylation / Ubiquitination | P24928 | 3 | 1.58 |
| **S35** | Phosphorylation | P24928 | 2 | 1.00 |
| **T37** | Phosphorylation | P24928 | 2 | 1.00 |
| **K42** | Acetylation / SUMOylation / Ubiquitination | P24928 | 4 | 2.00 |
| **T46** | Phosphorylation | P24928 | 2 | 1.00 |
| **T47** | Phosphorylation | P24928 | 2 | 1.00 |
| **K53** | Ubiquitination | P24928 | 2 | 1.00 |
| **K92** | Ubiquitination | P24928 | 2 | 1.00 |
| **K116** | Ubiquitination | P24928 | 2 | 1.00 |

| Residue | PTM Type(s) | UniProt accession | States (s<sub>i</sub>) | Bits per site log<sub>2</sub>(s<sub>i</sub>) |
| --- | --- | --- | --- | --- |
| S121 | Phosphorylation | P24928 | 2 | 1.00 |
| K125 | Ubiquitination | P24928 | 2 | 1.00 |
| K127 | Ubiquitination | P24928 | 2 | 1.00 |
| K132 | Ubiquitination | P24928 | 2 | 1.00 |
| K134 | Acetylation | P24928 | 2 | 1.00 |
| K139 | Acetylation | P24928 | 2 | 1.00 |
| Y145 | Phosphorylation | P24928 | 2 | 1.00 |
| K149 | SUMOylation / Ubiquitination | P24928 | 3 | 1.58 |
| K151 | SUMOylation / Ubiquitination | P24928 | 3 | 1.58 |
| K163 | SUMOylation / Ubiquitination | P24928 | 3 | 1.58 |
| T176 | Phosphorylation | P24928 | 2 | 1.00 |
| K177 | Acetylation / Ubiquitination | P24928 | 3 | 1.58 |
| K179 | Ubiquitination | P24928 | 2 | 1.00 |
| Y199 | Phosphorylation | P24928 | 2 | 1.00 |
| K203 | Ubiquitination | P24928 | 2 | 1.00 |
| S209 | Phosphorylation | P24928 | 2 | 1.00 |
| K212 | Ubiquitination | P24928 | 2 | 1.00 |
| K213 | SUMOylation / Ubiquitination | P24928 | 3 | 1.58 |
| S217 | Phosphorylation | P24928 | 2 | 1.00 |
| K226 | Ubiquitination | P24928 | 2 | 1.00 |
| C233 | Glutathionylation / Methylation | P24928 | 3 | 1.58 |
| S269 | Phosphorylation | P24928 | 2 | 1.00 |
| K279 | SUMOylation / Ubiquitination | P24928 | 3 | 1.58 |
| K285 | Acetylation / SUMOylation / Ubiquitination | P24928 | 4 | 2.00 |
| K308 | SUMOylation | P24928 | 2 | 1.00 |
| K331 | Ubiquitination | P24928 | 2 | 1.00 |
| K337 | Ubiquitination | P24928 | 2 | 1.00 |
| K340 | Ubiquitination | P24928 | 2 | 1.00 |
| K357 | Acetylation / Ubiquitination | P24928 | 3 | 1.58 |
| T368 | Phosphorylation | P24928 | 2 | 1.00 |
| S374 | Phosphorylation | P24928 | 2 | 1.00 |
| K417 | Ubiquitination | P24928 | 2 | 1.00 |
| R426 | Methylation | P24928 | 2 | 1.00 |
| K434 | Ubiquitination | P24928 | 2 | 1.00 |
| K445 | SUMOylation / Ubiquitination | P24928 | 3 | 1.58 |
| C451 | Glutathionylation | P24928 | 2 | 1.00 |
| K466 | SUMOylation / Ubiquitination | P24928 | 3 | 1.58 |
| T481 | Phosphorylation | P24928 | 2 | 1.00 |
| M520 | Sulfoxidation | P24928 | 2 | 1.00 |
| S530 | Phosphorylation | P24928 | 2 | 1.00 |
| T543 | Phosphorylation | P24928 | 2 | 1.00 |
| T549 | Phosphorylation | P24928 | 2 | 1.00 |
| K581 | SUMOylation / Ubiquitination | P24928 | 3 | 1.58 |
| T587 | Phosphorylation | P24928 | 2 | 1.00 |
| S607 | Phosphorylation | P24928 | 2 | 1.00 |
| T608 | Phosphorylation | P24928 | 2 | 1.00 |
| Y618 | Phosphorylation | P24928 | 2 | 1.00 |
| K619 | Acetylation / SUMOylation / Ubiquitination | P24928 | 4 | 2.00 |
| K627 | SUMOylation / Ubiquitination | P24928 | 3 | 1.58 |
| K642 | Acetylation / Ubiquitination | P24928 | 3 | 1.58 |
| K643 | Ubiquitination | P24928 | 2 | 1.00 |
| K697 | Ubiquitination | P24928 | 2 | 1.00 |
| T698 | Phosphorylation | P24928 | 2 | 1.00 |
| Y699 | Phosphorylation | P24928 | 2 | 1.00 |
| K707 | SUMOylation / Ubiquitination | P24928 | 3 | 1.58 |
| K708 | Methylation / Ubiquitination | P24928 | 3 | 1.58 |
| K710 | Acetylation / SUMOylation / Ubiquitination | P24928 | 4 | 2.00 |
| K719 | Ubiquitination | P24928 | 2 | 1.00 |
| T732 | Phosphorylation | P24928 | 2 | 1.00 |
| T736 | Phosphorylation | P24928 | 2 | 1.00 |
| K751 | Ubiquitination | P24928 | 2 | 1.00 |
| K758 | Ubiquitination | P24928 | 2 | 1.00 |
| S759 | Phosphorylation | P24928 | 2 | 1.00 |
| S761 | Phosphorylation | P24928 | 2 | 1.00 |
| Y763 | Phosphorylation | P24928 | 2 | 1.00 |
| K767 | Methylation / SUMOylation / Ubiquitination | P24928 | 4 | 2.00 |
| S768 | Phosphorylation | P24928 | 2 | 1.00 |
| S772 | Phosphorylation | P24928 | 2 | 1.00 |
| K775 | Ubiquitination | P24928 | 2 | 1.00 |
| S777 | Phosphorylation | P24928 | 2 | 1.00 |
| K778 | SUMOylation / Ubiquitination | P24928 | 3 | 1.58 |
| K796 | SUMOylation / Ubiquitination | P24928 | 3 | 1.58 |
| K803 | Acetylation / Methylation / SUMOylation / Ubiquitination | P24928 | 5 | 2.32 |
| K812 | Acetylation / SUMOylation / Ubiquitination | P24928 | 4 | 2.00 |
| K853 | SUMOylation / Ubiquitination | P24928 | 3 | 1.58 |
| T857 | Phosphorylation | P24928 | 2 | 1.00 |
| Y859 | Phosphorylation | P24928 | 2 | 1.00 |
| K866 | SUMOylation / Ubiquitination | P24928 | 3 | 1.58 |
| S867 | Phosphorylation | P24928 | 2 | 1.00 |
| S870 | Phosphorylation | P24928 | 2 | 1.00 |
| K874 | Ubiquitination | P24928 | 2 | 1.00 |
| Y875 | Phosphorylation | P24928 | 2 | 1.00 |
| T878 | Phosphorylation | P24928 | 2 | 1.00 |
| **S882** | Phosphorylation | P24928 | 2 | 1.00 |
| **K910** | Acetylation / SUMOylation / Ubiquitination | P24928 | 4 | 2.00 |
| **K914** | Ubiquitination | P24928 | 2 | 1.00 |
| **K918** | Ubiquitination | P24928 | 2 | 1.00 |
| **K919** | Ubiquitination | P24928 | 2 | 1.00 |
| **K940** | Acetylation / SUMOylation / Ubiquitination | P24928 | 4 | 2.00 |
| **T972** | Phosphorylation | P24928 | 2 | 1.00 |
| **S975** | Phosphorylation | P24928 | 2 | 1.00 |
| **K976** | SUMOylation / Ubiquitination | P24928 | 3 | 1.58 |
| **K992** | Ubiquitination | P24928 | 2 | 1.00 |
| **K1008** | Acetylation / Ubiquitination | P24928 | 3 | 1.58 |
| **K1014** | Acetylation / Ubiquitination | P24928 | 3 | 1.58 |
| **K1018** | Ubiquitination | P24928 | 2 | 1.00 |
| **K1019** | Ubiquitination | P24928 | 2 | 1.00 |
| **S1030** | Phosphorylation | P24928 | 2 | 1.00 |
| **C1050** | Methylation | P24928 | 2 | 1.00 |
| **Y1109** | Phosphorylation | P24928 | 2 | 1.00 |
| **K1125** | SUMOylation / Ubiquitination | P24928 | 3 | 1.58 |
| **K1132** | Ubiquitination | P24928 | 2 | 1.00 |
| **K1133** | Ubiquitination | P24928 | 2 | 1.00 |
| **K1135** | SUMOylation / Ubiquitination | P24928 | 3 | 1.58 |
| **S1138** | Phosphorylation | P24928 | 2 | 1.00 |
| **S1147** | Phosphorylation | P24928 | 2 | 1.00 |
| **K1155** | Ubiquitination | P24928 | 2 | 1.00 |
| **C1159** | Methylation | P24928 | 2 | 1.00 |
| **K1219** | Ubiquitination | P24928 | 2 | 1.00 |
| **K1225** | SUMOylation / Ubiquitination | P24928 | 3 | 1.58 |
| **K1234** | Ubiquitination | P24928 | 2 | 1.00 |
| K1254 | SUMOylation / Ubiquitination | P24928 | 3 | 1.58 |
| K1268 | SUMOylation / Ubiquitination | P24928 | 3 | 1.58 |
| K1278 | Ubiquitination | P24928 | 2 | 1.00 |
| C1287 | Glutathionylation | P24928 | 2 | 1.00 |
| K1306 | Ubiquitination | P24928 | 2 | 1.00 |
| Y1308 | Phosphorylation | P24928 | 2 | 1.00 |
| K1317 | Ubiquitination | P24928 | 2 | 1.00 |
| K1318 | Ubiquitination | P24928 | 2 | 1.00 |
| K1319 | Ubiquitination | P24928 | 2 | 1.00 |
| K1329 | Ubiquitination | P24928 | 2 | 1.00 |
| S1348 | Phosphorylation | P24928 | 2 | 1.00 |
| K1350 | Acetylation / Ubiquitination | P24928 | 3 | 1.58 |
| Y1383 | Phosphorylation | P24928 | 2 | 1.00 |
| S1387 | Phosphorylation | P24928 | 2 | 1.00 |
| S1391 | Phosphorylation | P24928 | 2 | 1.00 |
| Y1392 | Phosphorylation | P24928 | 2 | 1.00 |
| Y1395 | Phosphorylation | P24928 | 2 | 1.00 |
| T1406 | Phosphorylation | P24928 | 2 | 1.00 |
| T1415 | Phosphorylation | P24928 | 2 | 1.00 |
| T1424 | Phosphorylation | P24928 | 2 | 1.00 |
| K1429 | Ubiquitination | P24928 | 2 | 1.00 |
| K1479 | Ubiquitination | P24928 | 2 | 1.00 |
| K1481 | Ubiquitination | P24928 | 2 | 1.00 |
| S1505 | Phosphorylation | P24928 | 2 | 1.00 |
| S1508 | Phosphorylation | P24928 | 2 | 1.00 |
| S1514 | Phosphorylation | P24928 | 2 | 1.00 |
| T1518 | Phosphorylation | P24928 | 2 | 1.00 |
| T1525 | Phosphorylation | P24928 | 2 | 1.00 |
| **S1532** | Phosphorylation | P24928 | 2 | 1.00 |
| **S1534** | O-GlcNAcylation / Phosphorylation | P24928 | 3 | 1.58 |
| **T1540** | Phosphorylation | P24928 | 2 | 1.00 |
| **T1568** | Phosphorylation | P24928 | 2 | 1.00 |
| **S1571** | Phosphorylation | P24928 | 2 | 1.00 |
| **S1574** | Phosphorylation | P24928 | 2 | 1.00 |
| **S1590** | Phosphorylation | P24928 | 2 | 1.00 |
| **S1592** | Phosphorylation | P24928 | 2 | 1.00 |

**Supplementary Table S2.**
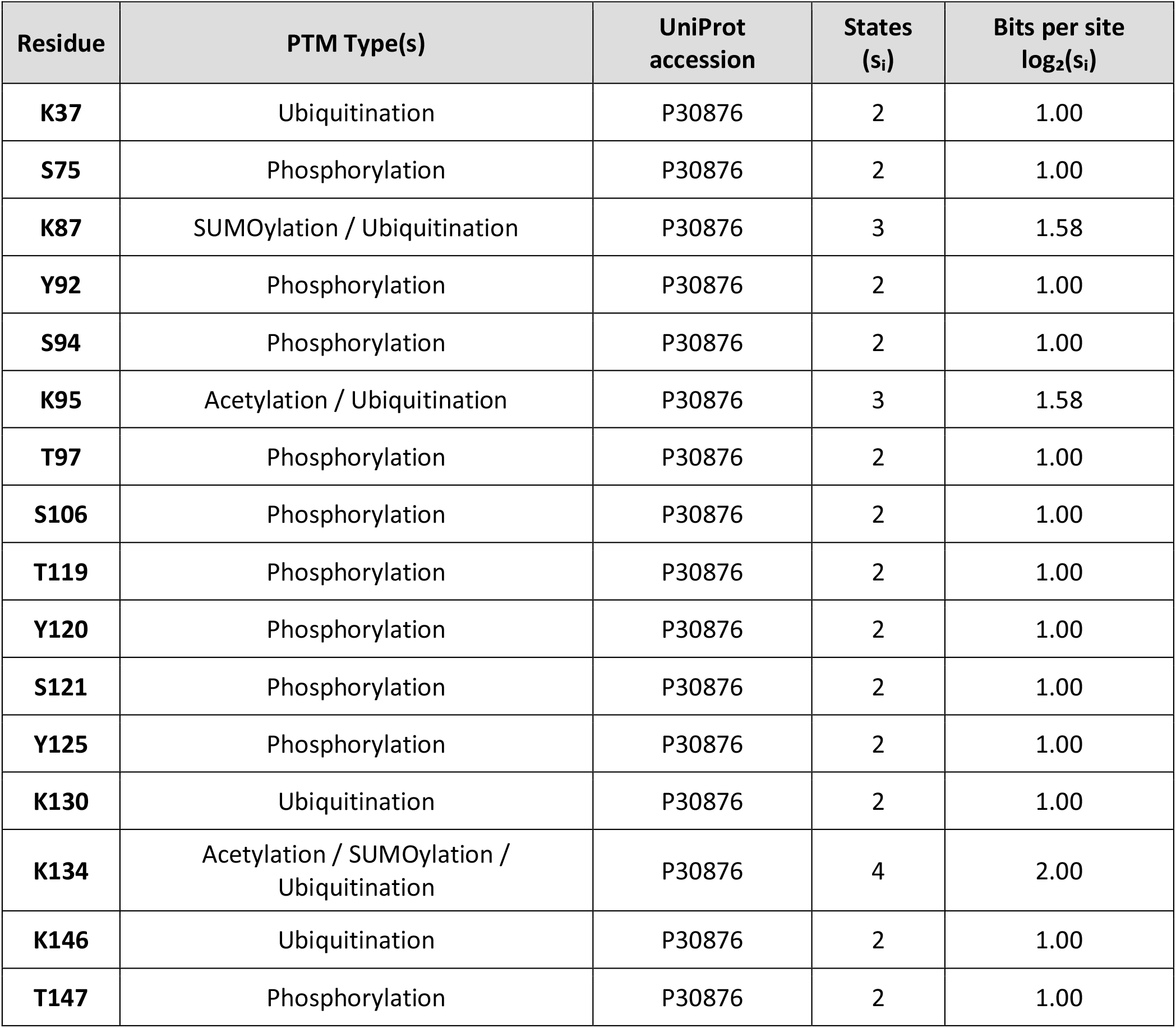
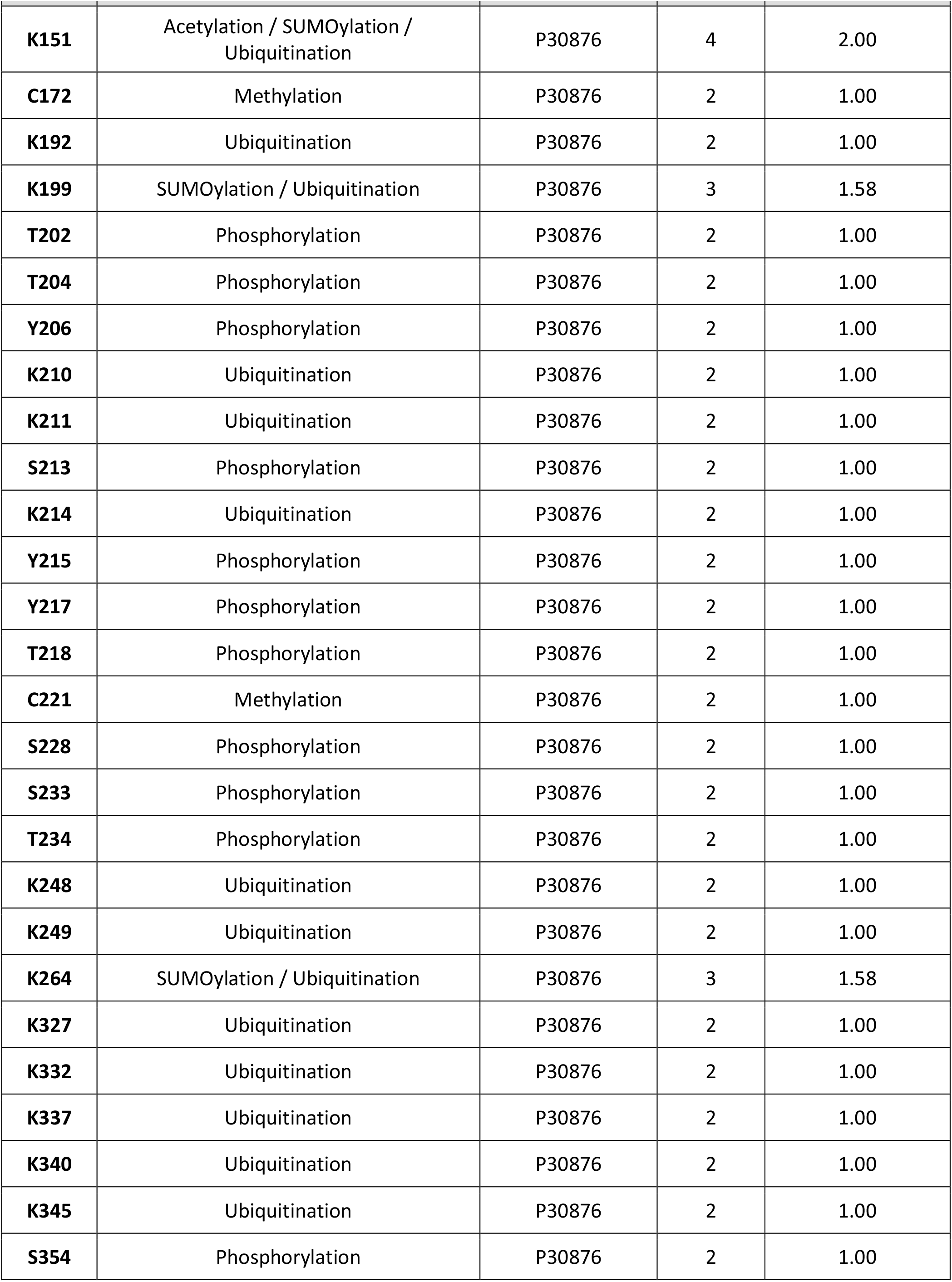
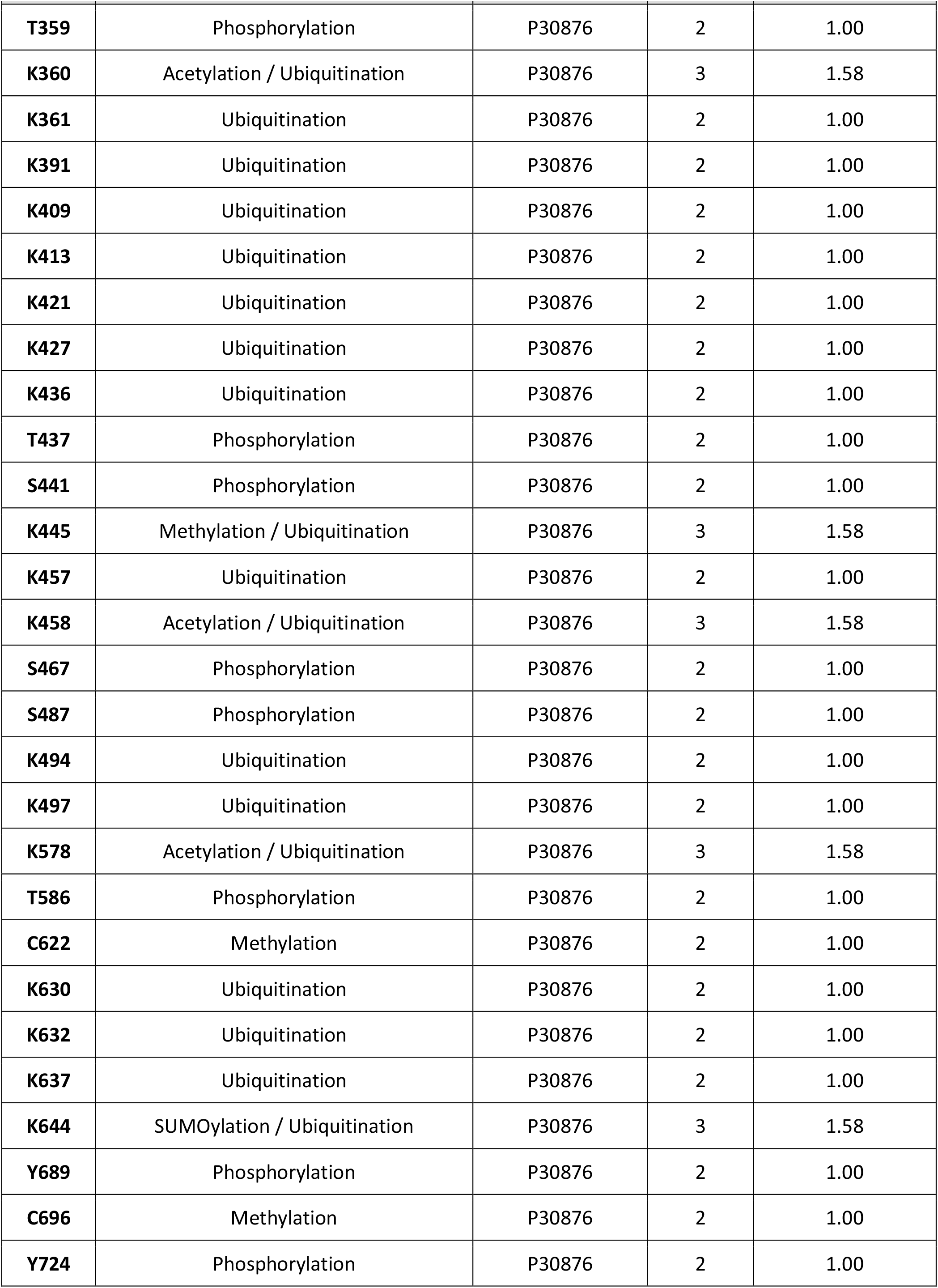
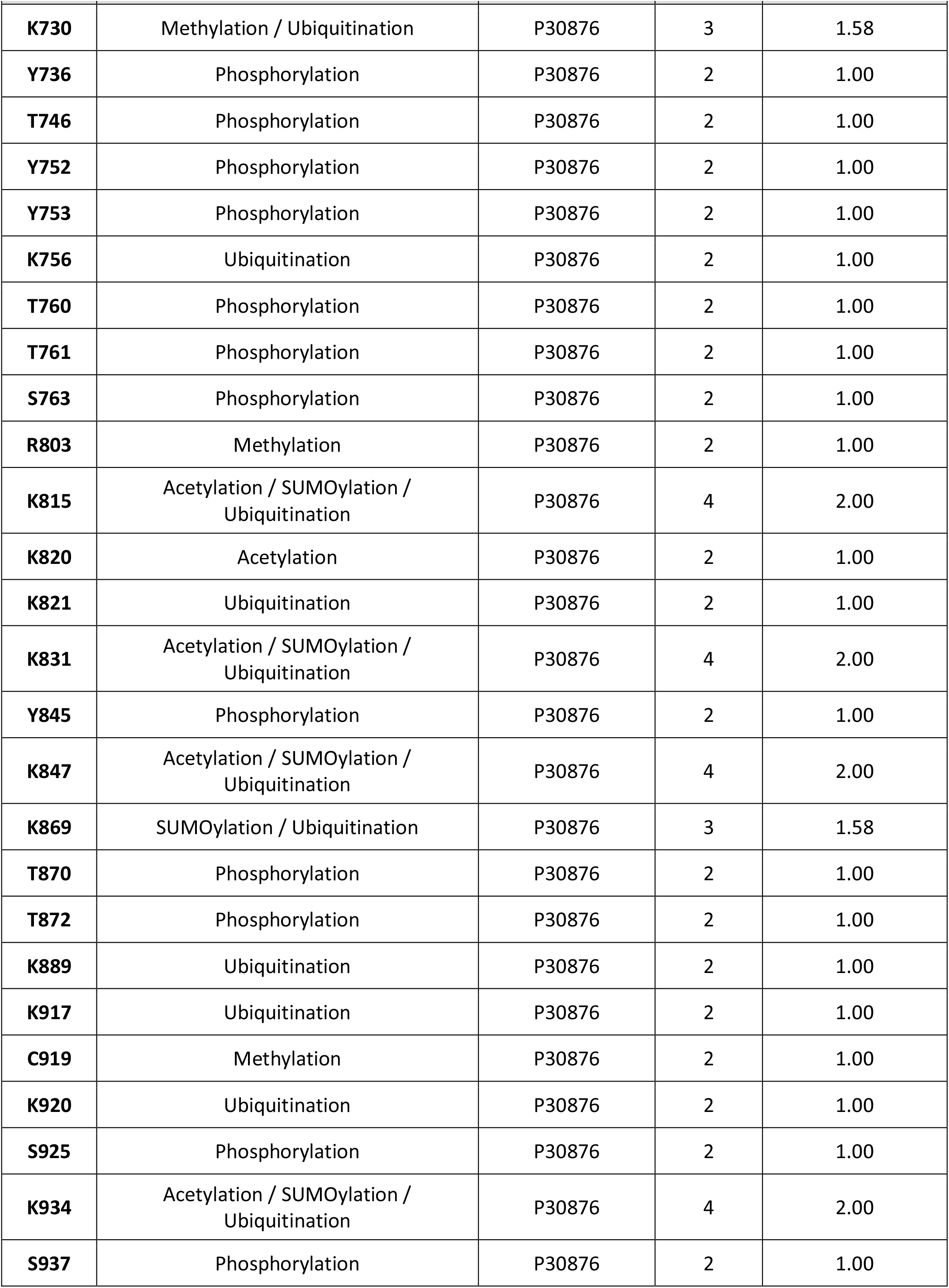
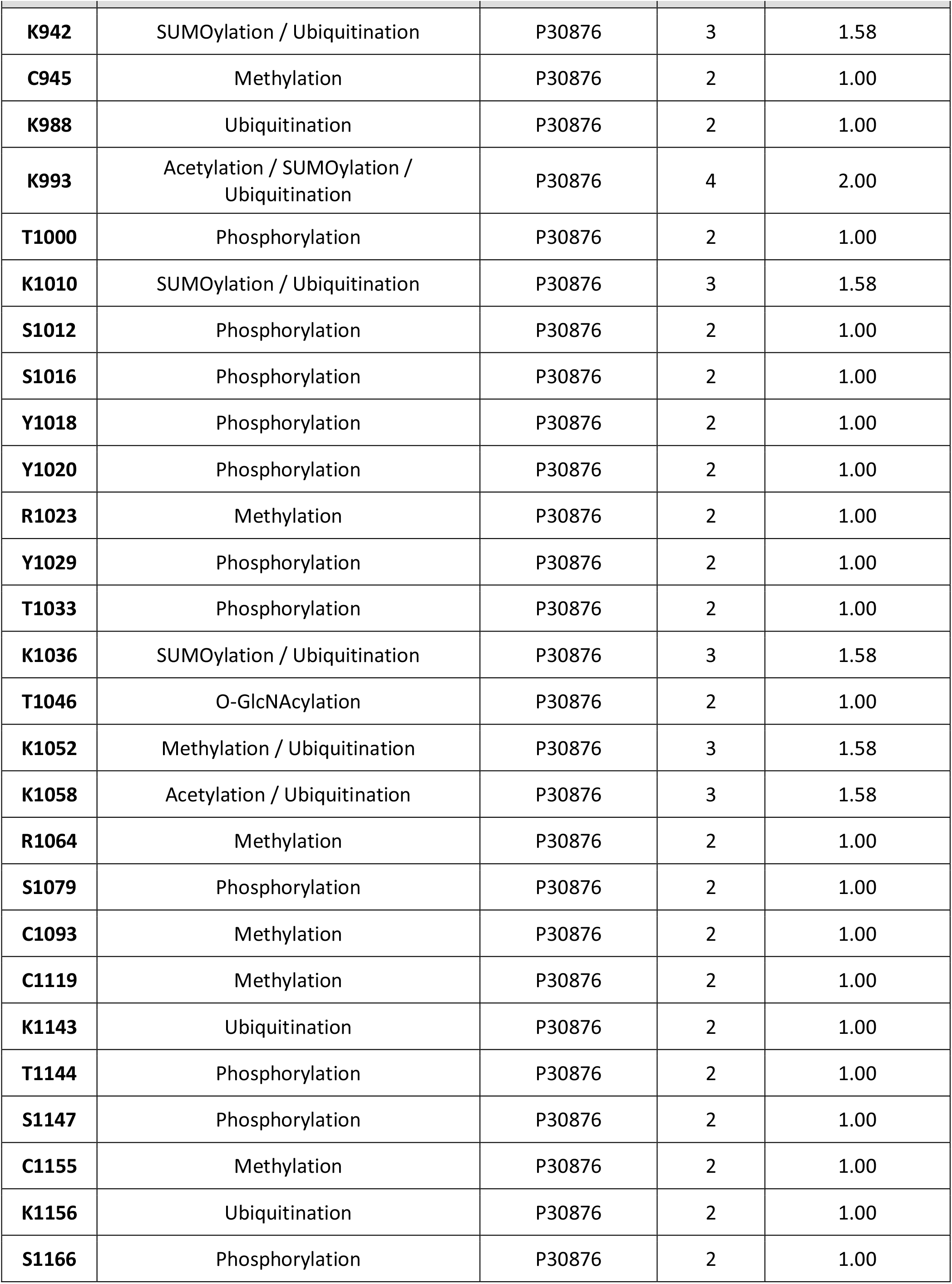
POLR2B (RPB2 / Rpb2)

| Residue | PTM Type(s) | UniProt accession | States (s<sub>i</sub>) | Bits per site log<sub>2</sub>(s<sub>i</sub>) |
| --- | --- | --- | --- | --- |
| **K37** | Ubiquitination | P30876 | 2 | 1.00 |
| **S75** | Phosphorylation | P30876 | 2 | 1.00 |
| **K87** | SUMOylation / Ubiquitination | P30876 | 3 | 1.58 |
| **Y92** | Phosphorylation | P30876 | 2 | 1.00 |
| **S94** | Phosphorylation | P30876 | 2 | 1.00 |
| **K95** | Acetylation / Ubiquitination | P30876 | 3 | 1.58 |
| **T97** | Phosphorylation | P30876 | 2 | 1.00 |
| **S106** | Phosphorylation | P30876 | 2 | 1.00 |
| **T119** | Phosphorylation | P30876 | 2 | 1.00 |
| **Y120** | Phosphorylation | P30876 | 2 | 1.00 |
| **S121** | Phosphorylation | P30876 | 2 | 1.00 |
| **Y125** | Phosphorylation | P30876 | 2 | 1.00 |
| **K130** | Ubiquitination | P30876 | 2 | 1.00 |
| **K134** | Acetylation / SUMOylation / Ubiquitination | P30876 | 4 | 2.00 |
| **K146** | Ubiquitination | P30876 | 2 | 1.00 |
| **T147** | Phosphorylation | P30876 | 2 | 1.00 |
| K151 | Acetylation / SUMOylation / Ubiquitination | P30876 | 4 | 2.00 |
| C172 | Methylation | P30876 | 2 | 1.00 |
| K192 | Ubiquitination | P30876 | 2 | 1.00 |
| K199 | SUMOylation / Ubiquitination | P30876 | 3 | 1.58 |
| T202 | Phosphorylation | P30876 | 2 | 1.00 |
| T204 | Phosphorylation | P30876 | 2 | 1.00 |
| Y206 | Phosphorylation | P30876 | 2 | 1.00 |
| K210 | Ubiquitination | P30876 | 2 | 1.00 |
| K211 | Ubiquitination | P30876 | 2 | 1.00 |
| S213 | Phosphorylation | P30876 | 2 | 1.00 |
| K214 | Ubiquitination | P30876 | 2 | 1.00 |
| Y215 | Phosphorylation | P30876 | 2 | 1.00 |
| Y217 | Phosphorylation | P30876 | 2 | 1.00 |
| T218 | Phosphorylation | P30876 | 2 | 1.00 |
| C221 | Methylation | P30876 | 2 | 1.00 |
| S228 | Phosphorylation | P30876 | 2 | 1.00 |
| S233 | Phosphorylation | P30876 | 2 | 1.00 |
| T234 | Phosphorylation | P30876 | 2 | 1.00 |
| K248 | Ubiquitination | P30876 | 2 | 1.00 |
| K249 | Ubiquitination | P30876 | 2 | 1.00 |
| K264 | SUMOylation / Ubiquitination | P30876 | 3 | 1.58 |
| K327 | Ubiquitination | P30876 | 2 | 1.00 |
| K332 | Ubiquitination | P30876 | 2 | 1.00 |
| K337 | Ubiquitination | P30876 | 2 | 1.00 |
| K340 | Ubiquitination | P30876 | 2 | 1.00 |
| K345 | Ubiquitination | P30876 | 2 | 1.00 |
| S354 | Phosphorylation | P30876 | 2 | 1.00 |
| T359 | Phosphorylation | P30876 | 2 | 1.00 |
| K360 | Acetylation / Ubiquitination | P30876 | 3 | 1.58 |
| K361 | Ubiquitination | P30876 | 2 | 1.00 |
| K391 | Ubiquitination | P30876 | 2 | 1.00 |
| K409 | Ubiquitination | P30876 | 2 | 1.00 |
| K413 | Ubiquitination | P30876 | 2 | 1.00 |
| K421 | Ubiquitination | P30876 | 2 | 1.00 |
| K427 | Ubiquitination | P30876 | 2 | 1.00 |
| K436 | Ubiquitination | P30876 | 2 | 1.00 |
| T437 | Phosphorylation | P30876 | 2 | 1.00 |
| S441 | Phosphorylation | P30876 | 2 | 1.00 |
| K445 | Methylation / Ubiquitination | P30876 | 3 | 1.58 |
| K457 | Ubiquitination | P30876 | 2 | 1.00 |
| K458 | Acetylation / Ubiquitination | P30876 | 3 | 1.58 |
| S467 | Phosphorylation | P30876 | 2 | 1.00 |
| S487 | Phosphorylation | P30876 | 2 | 1.00 |
| K494 | Ubiquitination | P30876 | 2 | 1.00 |
| K497 | Ubiquitination | P30876 | 2 | 1.00 |
| K578 | Acetylation / Ubiquitination | P30876 | 3 | 1.58 |
| T586 | Phosphorylation | P30876 | 2 | 1.00 |
| C622 | Methylation | P30876 | 2 | 1.00 |
| K630 | Ubiquitination | P30876 | 2 | 1.00 |
| K632 | Ubiquitination | P30876 | 2 | 1.00 |
| K637 | Ubiquitination | P30876 | 2 | 1.00 |
| K644 | SUMOylation / Ubiquitination | P30876 | 3 | 1.58 |
| Y689 | Phosphorylation | P30876 | 2 | 1.00 |
| C696 | Methylation | P30876 | 2 | 1.00 |
| Y724 | Phosphorylation | P30876 | 2 | 1.00 |
| K730 | Methylation / Ubiquitination | P30876 | 3 | 1.58 |
| Y736 | Phosphorylation | P30876 | 2 | 1.00 |
| T746 | Phosphorylation | P30876 | 2 | 1.00 |
| Y752 | Phosphorylation | P30876 | 2 | 1.00 |
| Y753 | Phosphorylation | P30876 | 2 | 1.00 |
| K756 | Ubiquitination | P30876 | 2 | 1.00 |
| T760 | Phosphorylation | P30876 | 2 | 1.00 |
| T761 | Phosphorylation | P30876 | 2 | 1.00 |
| S763 | Phosphorylation | P30876 | 2 | 1.00 |
| R803 | Methylation | P30876 | 2 | 1.00 |
| K815 | Acetylation / SUMOylation / Ubiquitination | P30876 | 4 | 2.00 |
| K820 | Acetylation | P30876 | 2 | 1.00 |
| K821 | Ubiquitination | P30876 | 2 | 1.00 |
| K831 | Acetylation / SUMOylation / Ubiquitination | P30876 | 4 | 2.00 |
| Y845 | Phosphorylation | P30876 | 2 | 1.00 |
| K847 | Acetylation / SUMOylation / Ubiquitination | P30876 | 4 | 2.00 |
| K869 | SUMOylation / Ubiquitination | P30876 | 3 | 1.58 |
| T870 | Phosphorylation | P30876 | 2 | 1.00 |
| T872 | Phosphorylation | P30876 | 2 | 1.00 |
| K889 | Ubiquitination | P30876 | 2 | 1.00 |
| K917 | Ubiquitination | P30876 | 2 | 1.00 |
| C919 | Methylation | P30876 | 2 | 1.00 |
| K920 | Ubiquitination | P30876 | 2 | 1.00 |
| S925 | Phosphorylation | P30876 | 2 | 1.00 |
| K934 | Acetylation / SUMOylation / Ubiquitination | P30876 | 4 | 2.00 |
| S937 | Phosphorylation | P30876 | 2 | 1.00 |
| K942 | SUMOylation / Ubiquitination | P30876 | 3 | 1.58 |
| C945 | Methylation | P30876 | 2 | 1.00 |
| K988 | Ubiquitination | P30876 | 2 | 1.00 |
| K993 | Acetylation / SUMOylation / Ubiquitination | P30876 | 4 | 2.00 |
| T1000 | Phosphorylation | P30876 | 2 | 1.00 |
| K1010 | SUMOylation / Ubiquitination | P30876 | 3 | 1.58 |
| S1012 | Phosphorylation | P30876 | 2 | 1.00 |
| S1016 | Phosphorylation | P30876 | 2 | 1.00 |
| Y1018 | Phosphorylation | P30876 | 2 | 1.00 |
| Y1020 | Phosphorylation | P30876 | 2 | 1.00 |
| R1023 | Methylation | P30876 | 2 | 1.00 |
| Y1029 | Phosphorylation | P30876 | 2 | 1.00 |
| T1033 | Phosphorylation | P30876 | 2 | 1.00 |
| K1036 | SUMOylation / Ubiquitination | P30876 | 3 | 1.58 |
| T1046 | O-GlcNAcylation | P30876 | 2 | 1.00 |
| K1052 | Methylation / Ubiquitination | P30876 | 3 | 1.58 |
| K1058 | Acetylation / Ubiquitination | P30876 | 3 | 1.58 |
| R1064 | Methylation | P30876 | 2 | 1.00 |
| S1079 | Phosphorylation | P30876 | 2 | 1.00 |
| C1093 | Methylation | P30876 | 2 | 1.00 |
| C1119 | Methylation | P30876 | 2 | 1.00 |
| K1143 | Ubiquitination | P30876 | 2 | 1.00 |
| T1144 | Phosphorylation | P30876 | 2 | 1.00 |
| S1147 | Phosphorylation | P30876 | 2 | 1.00 |
| C1155 | Methylation | P30876 | 2 | 1.00 |
| K1156 | Ubiquitination | P30876 | 2 | 1.00 |
| S1166 | Phosphorylation | P30876 | 2 | 1.00 |

**Supplementary Table S3.** POLR2C (RPB3 / Rpb3)

| **Residue** | **PTM Type(s)** | **UniProt accession** | **States (s<sub>i</sub>)** | **Bits per site log<sub>2</sub>(s<sub>i</sub>)** |
| --- | --- | --- | --- | --- |
| **Y3** | Phosphorylation | P19387 | 2 | 1.00 |
| **T8** | Phosphorylation | P19387 | 2 | 1.00 |
| **T12** | Phosphorylation | P19387 | 2 | 1.00 |
| **T15** | Phosphorylation | P19387 | 2 | 1.00 |
| **K20** | SUMOylation / Ubiquitination | P19387 | 3 | 1.58 |
| **T26** | Phosphorylation | P19387 | 2 | 1.00 |
| **S33** | Phosphorylation | P19387 | 2 | 1.00 |
| **K81** | SUMOylation / Ubiquitination | P19387 | 3 | 1.58 |
| **S122** | Phosphorylation | P19387 | 2 | 1.00 |
| **S124** | Phosphorylation | P19387 | 2 | 1.00 |
| **Y142** | Phosphorylation | P19387 | 2 | 1.00 |
| **K152** | Ubiquitination | P19387 | 2 | 1.00 |
| **K171** | SUMOylation | P19387 | 2 | 1.00 |
| **K175** | SUMOylation / Ubiquitination | P19387 | 3 | 1.58 |
| **K199** | SUMOylation / Ubiquitination | P19387 | 3 | 1.58 |
| **K205** | SUMOylation / Ubiquitination | P19387 | 3 | 1.58 |
| **S206** | Phosphorylation | P19387 | 2 | 1.00 |
| **Y208** | Phosphorylation | P19387 | 2 | 1.00 |
| **S209** | Phosphorylation | P19387 | 2 | 1.00 |
| **S216** | Phosphorylation | P19387 | 2 | 1.00 |
| **Y220** | Phosphorylation | P19387 | 2 | 1.00 |
| **K225** | Acetylation / SUMOylation / Ubiquitination | P19387 | 4 | 2.00 |
| **S235** | Phosphorylation | P19387 | 2 | 1.00 |
| **K253** | Ubiquitination | P19387 | 2 | 1.00 |
| **K254** | Ubiquitination | P19387 | 2 | 1.00 |
| **K255** | SUMOylation / Ubiquitination | P19387 | 3 | 1.58 |

| Residue | PTM Type(s) | UniProt accession | States (s<sub>i</sub>) | Bits per site log<sub>2</sub>(s<sub>i</sub>) |
| --- | --- | --- | --- | --- |
| **S257** | Phosphorylation | P19387 | 2 | 1.00 |
| **T261** | Phosphorylation | P19387 | 2 | 1.00 |
| **S264** | Phosphorylation | P19387 | 2 | 1.00 |

**Supplementary Table S4.** POLR2D (RPB4 / Rpb4)

| Residue | PTM Type(s) | UniProt accession | States (s<sub>i</sub>) | Bits per site log<sub>2</sub>(s<sub>i</sub>) |
| --- | --- | --- | --- | --- |
| **S18** | Phosphorylation | O15514 | 2 | 1.00 |
| **T28** | Phosphorylation | O15514 | 2 | 1.00 |
| **S35** | Phosphorylation | O15514 | 2 | 1.00 |
| **K45** | Ubiquitination | O15514 | 2 | 1.00 |
| **Y67** | Phosphorylation | O15514 | 2 | 1.00 |
| **S72** | Phosphorylation | O15514 | 2 | 1.00 |
| **K91** | Ubiquitination | O15514 | 2 | 1.00 |
| **K94** | Acetylation | O15514 | 2 | 1.00 |
| **K112** | Ubiquitination | O15514 | 2 | 1.00 |
| **K137** | Ubiquitination | O15514 | 2 | 1.00 |

**Supplementary Table S5.** POLR2E (RPABC1 / Rpb5)

| Residue | PTM Type(s) | UniProt accession | States (s<sub>i</sub>) | Bits per site log<sub>2</sub>(s<sub>i</sub>) |
| --- | --- | --- | --- | --- |
| **M1** | Acetylation | P19388 | 2 | 1.00 |
| **T7** | Phosphorylation | P19388 | 2 | 1.00 |
| **Y8** | Phosphorylation | P19388 | 2 | 1.00 |
| **R9** | Methylation | P19388 | 2 | 1.00 |
| **K15** | Ubiquitination | P19388 | 2 | 1.00 |
| **T29** | Phosphorylation | P19388 | 2 | 1.00 |
| **T56** | Phosphorylation | P19388 | 2 | 1.00 |
| **T59** | Phosphorylation | P19388 | 2 | 1.00 |
| K81 | SUMOylation / Ubiquitination | P19388 | 3 | 1.58 |
| K85 | Ubiquitination | P19388 | 2 | 1.00 |
| K88 | Acetylation / Ubiquitination | P19388 | 3 | 1.58 |
| Y90 | Phosphorylation | P19388 | 2 | 1.00 |
| M110 | Sulfoxidation | P19388 | 2 | 1.00 |
| T111 | Phosphorylation | P19388 | 2 | 1.00 |
| S113 | Phosphorylation | P19388 | 2 | 1.00 |
| K115 | SUMOylation / Ubiquitination | P19388 | 3 | 1.58 |
| K153 | SUMOylation / Ubiquitination | P19388 | 3 | 1.58 |
| T157 | Phosphorylation | P19388 | 2 | 1.00 |
| K186 | Ubiquitination | P19388 | 2 | 1.00 |
| K192 | Acetylation / SUMOylation / Ubiquitination | P19388 | 4 | 2.00 |
| S197 | Phosphorylation | P19388 | 2 | 1.00 |
| T199 | Phosphorylation | P19388 | 2 | 1.00 |

**Supplementary Table S6.**
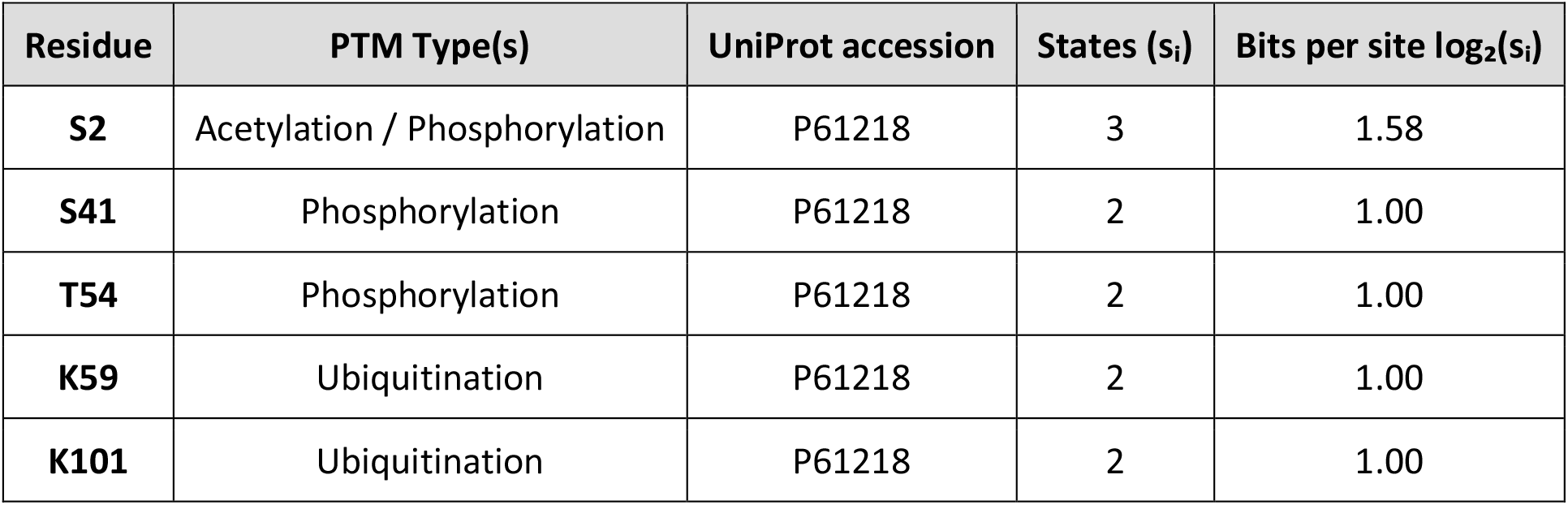
POLR2F (RPABC2 / Rpb6)

**Supplementary Table S7.** POLR2G (RPB7 / Rpb7)

| Residue | PTM Type(s) | UniProt accession | States (s<sub>i</sub>) | Bits per site log<sub>2</sub>(s<sub>i</sub>) |
| --- | --- | --- | --- | --- |
| Y3 | Phosphorylation | P62487 | 2 | 1.00 |
| S6 | Phosphorylation | P62487 | 2 | 1.00 |
| **Y17** | Phosphorylation | P62487 | 2 | 1.00 |
| **K27** | Ubiquitination | P62487 | 2 | 1.00 |
| **K29** | Acetylation / SUMOylation / Ubiquitination | P62487 | 4 | 2.00 |
| **C38** | Methylation | P62487 | 2 | 1.00 |
| **K71** | Ubiquitination | P62487 | 2 | 1.00 |
| **K73** | SUMOylation / Ubiquitination | P62487 | 3 | 1.58 |
| **K81** | Acetylation / SUMOylation / Ubiquitination | P62487 | 4 | 2.00 |
| **K94** | Ubiquitination | P62487 | 2 | 1.00 |
| **C106** | Methylation | P62487 | 2 | 1.00 |
| **K129** | SUMOylation / Ubiquitination | P62487 | 3 | 1.58 |
| **M131** | Sulfoxidation | P62487 | 2 | 1.00 |
| **K146** | Ubiquitination | P62487 | 2 | 1.00 |
| **K154** | Ubiquitination | P62487 | 2 | 1.00 |

**Supplementary Table S8.** POLR2H (RPABC3 / Rpb8)

| Residue | PTM Type(s) | UniProt accession | States (s<sub>i</sub>) | Bits per site log<sub>2</sub>(s<sub>i</sub>) |
| --- | --- | --- | --- | --- |
| **A2** | Acetylation | P52434 | 2 | 1.00 |
| **K13** | SUMOylation / Ubiquitination | P52434 | 3 | 1.58 |
| **K20** | Acetylation / SUMOylation / Ubiquitination | P52434 | 4 | 2.00 |
| **K21** | Ubiquitination | P52434 | 2 | 1.00 |
| **C30** | Methylation | P52434 | 2 | 1.00 |
| **K36** | Acetylation | P52434 | 2 | 1.00 |
| **K59** | Ubiquitination | P52434 | 2 | 1.00 |
| **Y65** | Phosphorylation | P52434 | 2 | 1.00 |
| **Y75** | Phosphorylation | P52434 | 2 | 1.00 |
| **T78** | Phosphorylation | P52434 | 2 | 1.00 |
| K82 | Ubiquitination | P52434 | 2 | 1.00 |
| K83 | Ubiquitination | P52434 | 2 | 1.00 |
| Y90 | Phosphorylation | P52434 | 2 | 1.00 |
| M92 | Sulfoxidation | P52434 | 2 | 1.00 |
| K95 | Ubiquitination | P52434 | 2 | 1.00 |
| T104 | Phosphorylation | P52434 | 2 | 1.00 |
| S105 | Phosphorylation | P52434 | 2 | 1.00 |
| K110 | Ubiquitination | P52434 | 2 | 1.00 |
| K111 | Ubiquitination | P52434 | 2 | 1.00 |
| K118 | Ubiquitination | P52434 | 2 | 1.00 |
| Y118 | Phosphorylation | P52434 | 2 | 1.00 |
| K119 | Ubiquitination | P52434 | 2 | 1.00 |
| Y142 | Phosphorylation | P52434 | 2 | 1.00 |
| K146 | Ubiquitination | P52434 | 2 | 1.00 |
| K147 | Ubiquitination | P52434 | 2 | 1.00 |

**Supplementary Table S9.** POLR2I (RPB9 / Rpb9)

| Residue | PTM Type(s) | UniProt accession | States (s<sub>i</sub>) | Bits per site log<sub>2</sub>(s<sub>i</sub>) |
| --- | --- | --- | --- | --- |
| M1 | Acetylation | P36954 | 2 | 1.00 |
| Y7 | Phosphorylation | P36954 | 2 | 1.00 |
| K27 | Ubiquitination | P36954 | 2 | 1.00 |
| Y37 | Phosphorylation | P36954 | 2 | 1.00 |
| S51 | Phosphorylation | P36954 | 2 | 1.00 |
| Y54 | Phosphorylation | P36954 | 2 | 1.00 |
| T59 | Phosphorylation | P36954 | 2 | 1.00 |
| T66 | Phosphorylation | P36954 | 2 | 1.00 |
| S73 | Phosphorylation | P36954 | 2 | 1.00 |
| T77 | Phosphorylation | P36954 | 2 | 1.00 |

| Residue | PTM Type(s) | UniProt accession | States ($s_i$) | Bits per site $\log_2(s_i)$ |
| --- | --- | --- | --- | --- |
| **T81** | Phosphorylation | P36954 | 2 | 1.00 |
| **S101** | Phosphorylation | P36954 | 2 | 1.00 |
| **Y111** | Phosphorylation | P36954 | 2 | 1.00 |
| **Y112** | Phosphorylation | P36954 | 2 | 1.00 |
| **T115** | Phosphorylation | P36954 | 2 | 1.00 |

**Supplementary Table S10.** POLR2L (RPABC5 / Rpb10)

| Residue | PTM Type(s) | UniProt accession | States ($s_i$) | Bits per site $\log_2(s_i)$ |
| --- | --- | --- | --- | --- |
| **C10** | Methylation | P62875 | 2 | 1.00 |
| **K12** | Ubiquitination | P62875 | 2 | 1.00 |
| **K17** | Ubiquitination | P62875 | 2 | 1.00 |
| **K41** | Ubiquitination | P62875 | 2 | 1.00 |
| **M48** | Methylation | P62875 | 2 | 1.00 |
| **K58** | Ubiquitination | P62875 | 2 | 1.00 |
| **Y62** | Phosphorylation | P62875 | 2 | 1.00 |
| **K67** | Ubiquitination | P62875 | 2 | 1.00 |

**Supplementary Table S11.**
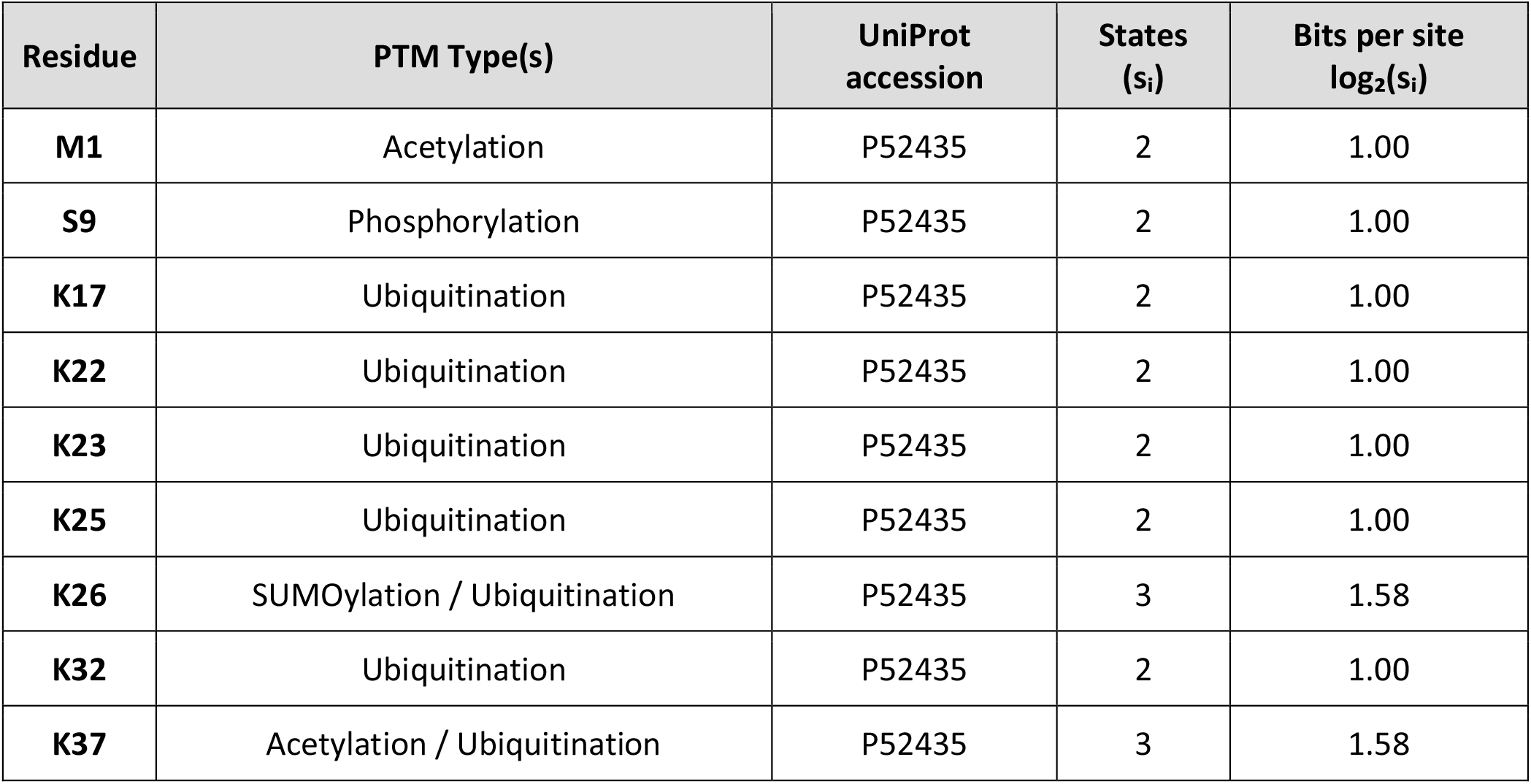
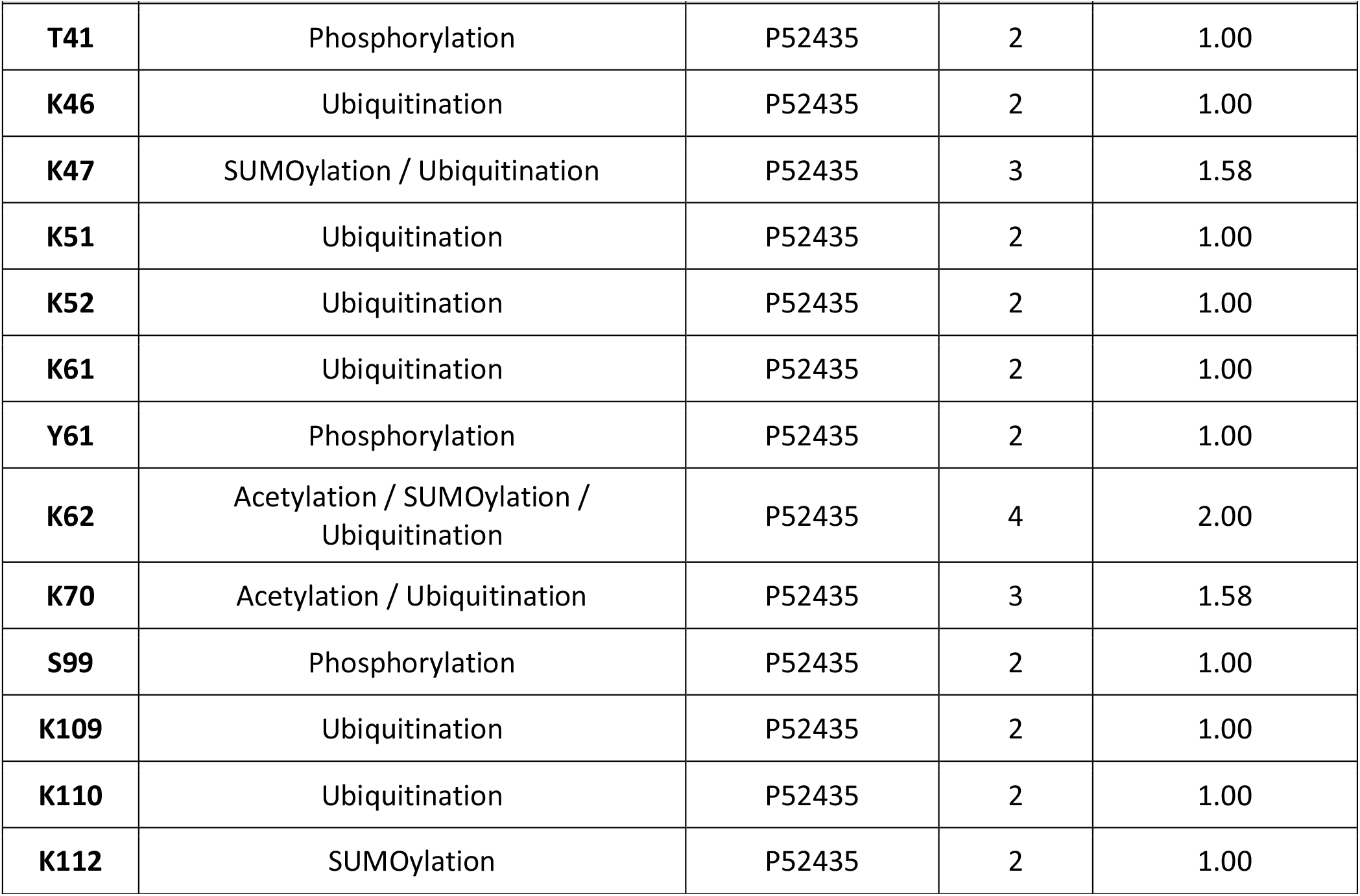
POLR2J (RPB11-a / Rpb11)

**Supplementary Table S12.** POLR2K (RPABC4 / Rpb12)

| Residue | PTM Type(s) | UniProt accession | States (s<sub>i</sub>) | Bits per site log<sub>2</sub>(s<sub>i</sub>) |
| --- | --- | --- | --- | --- |
| **M1** | Acetylation | P53803 | 2 | 1.00 |
| **K5** | Ubiquitination | P53803 | 2 | 1.00 |
| **C22** | Methylation | P53803 | 2 | 1.00 |

