## Supplementary material for "Multi-timescale PTM architecture in human RNA polymerase II": notebook_1.ipynb

```
# %% [markdown]
# Multi-timescale PTM architecture in human RNA polymerase II 
## Notebook 1 — PTM and protein sequence data retrieval
────────────────────────────────────────────────────────

This notebook (Supplementary Data File 1) generates all input datasets required for downstream quantitative analyses performed in `notebook_2.ipynb` (Supplementary Data File 2). Execution is required only to rebuild these files from current databases or to audit their provenance: when the frozen `uniprot_polII_seqs_2026_05_22.tsv` (Supplementary Data File 3) and `ptms_polII_2026_05_22.tsv` (Supplementary Data File 4) are present in the working directory, every download cell detects these cached files and returns without network access. If those two files are missing from the working directory this notebook will produce two frozen output files:

| File | Content |
|------|--------|
| `uniprot_polII_seqs_{CURRENT_DATE}.tsv` | Canonical UniProt sequences (UniSave version-pinned per subunit) |
| `ptms_polII_{CURRENT_DATE}.tsv` | Curated PTM atlas integrating UniProt, dbPTM, iPTMnet, PhosphoSitePlus, GlyGen, and literature mining across all RNA Pol II subunits |

These outputs define the reproducible reference state of the system and are required for all subsequent information-theoretic and structural analyses.

All database queries are version-controlled when possible. If live endpoints are used, results are explicitly cached to ensure reproducibility of downstream computations.

If the provided frozen output files are missing or overwritten with updated values, downstream notebooks will not reproduce manuscript values.

**Software requirements:** Python ≥ 3.10 | pandas ≥ 1.3 | requests ≥ 2.20 | IPython ≥ 7.0 | beautifulsoup4 ≥ 4.9

---

## Reproducibility workflow overview

Full descriptions are provided in Supplementary Information S4. The table below is an informative index only.

The first cell in this notebook (***Environment check***) checks minimum software requirements and raises an error if unmet. It is convenient to run it before the rest of the analysis to prevent errors downstream.

| Reproducibility Module | Stage | Contents |
|------|-------|----------|
| **1** | Sequence data (input) | Retrieves canonical UniProt protein sequences using versioned identifiers to ensure reproducibility. Writes a local sequence snapshot `uniprot_polII_seqs_{CURRENT_DATE}.tsv`. |
| **2–7** | PTM data acquisition | Downloads post-translational modification (PTM) annotations from multiple databases (UniProt curated, EBI proteomics, dbPTM, iPTMnet, PhosphoSitePlus, GlyGen). |
| **8** | CTD heptad position map | Reconstructs the CTD heptad architecture of Rpb1 (P24928), mapping each residue to its structural position within heptad repeats. |
| **9** | Alignment of PTMs across data sources | Integrates PTM annotations across all sources into a residue-level side-by-side wide-format comparison to assess agreement, redundancy, and coverage. |
| **10** | PTM data curation and integration | Harmonizes PTM nomenclature, removes inconsistencies, removes irreversible PTMs, and builds the final unified long-form dataset of unique (Accession, Residue, PTM) entries across all databases. Generates the reproducible snapshot `ptms_polII_{CURRENT_DATE}.tsv` file. |

## Key outputs

The pipeline produces two persistent files:

- **Protein sequence snapshot** (Supplementary Data File 3)
  `uniprot_polII_seqs_{CURRENT_DATE}.tsv`  
  Saved in: **project root directory (same folder as the notebook execution path)**  
  → Contains canonical UniProt sequences used for residue mapping.

- **Final PTM atlas (main manuscript dataset)** (Supplementary Data File 4)
  `ptms_polII_{CURRENT_DATE}.tsv`  
  Saved in: **project root directory (same folder as the notebook execution path)**  
  → Fully curated, harmonized, and integrated PTM dataset across all sources.

---

## Reproducibility note

All intermediate datasets are generated in-memory and can be fully reconstructed from the original database queries. Only the final curated snapshots are written to disk to ensure computational reproducibility while avoiding unnecessary external dependencies.

# %% [code]
## Environment check.
# Verifies that the Python environment meets the minimum requirements for running the notebook.

import sys, importlib, re

# ============================================================
# Python
# ============================================================

_py = sys.version_info
assert _py >= (3, 10), f"Python ≥ 3.10 required, got {_py.major}.{_py.minor}"

# ============================================================
# Third-party requirements
# ============================================================

_REQUIRED = {
    "pandas":   (1, 3),   # data manipulation
    "requests": (2, 20),  # timeout parameter
    "bs4":      (4, 9),   # BeautifulSoup HTML parsing (dbPTM, PhosphoSitePlus)
    "IPython":  (7, 0),   # display(), HTML()
}

def _parse(ver_str):
    """Numeric version tuple, ignoring pre-release suffixes (e.g. '1.3.0rc1')."""
    return tuple(int(x) for x in re.findall(r'\d+', ver_str.split('+')[0])[:3])

_ok, _fail = [], []
for _pkg, _min in _REQUIRED.items():
    try:
        _ver_str = importlib.import_module(_pkg).__version__
        _ver = _parse(_ver_str)
        _min_str = ".".join(map(str, _min))
        if _ver >= _min:
            _ok.append(f"  \u2713  {_pkg:<10} {_ver_str}")
        else:
            _fail.append(f"  \u2717  {_pkg:<10} {_ver_str}  (need \u2265 {_min_str})")
    except ImportError:
        _fail.append(f"  \u2717  {_pkg:<10} NOT INSTALLED  (need \u2265 {'.'.join(map(str, _min))})")

# ============================================================
# Report 
# ============================================================

print(f"  \u2713  {'Python':<10} {_py.major}.{_py.minor}.{_py.micro}")
print(*_ok, sep="\n")
if _fail:
    print("\n  FAILED:")
    print(*_fail, sep="\n")
    raise EnvironmentError("Environment check failed; install missing/outdated packages before running.")
else:
    print("\n  All requirements satisfied. Ready to run.")

# %% [code]
## Reproducibility Module 1. Canonical UniProt Sequence Validation

# ============================================================
# Imports
# ============================================================

from datetime import date
import os, requests
import pandas as pd

# ============================================================
# Configuration
# ============================================================

import pathlib as _pl
SNAPSHOT_DATE = date.today().strftime("%Y_%m_%d")  # ISO date format
_existing_seq = sorted(_pl.Path('.').glob('uniprot_polII_seqs_*.tsv'))  # Select the most recent existing cache file
seq_cache = str(_existing_seq[-1]) if _existing_seq \
            else f"uniprot_polII_seqs_{SNAPSHOT_DATE}.tsv"  # dated name only when writing new
acc_versions = {   # pinned to the versions cited in the manuscript
    'P24928': 248, 'P30876': 231, 'P19387': 235, 'O15514': 211,
    'P19388': 237, 'P61218': 187, 'P62487': 178, 'P52434': 230,
    'P36954': 221, 'P62875': 187, 'P52435': 215, 'P53803': 209,
}

# ============================================================
# Download from UniProt
# ============================================================

if os.path.exists(seq_cache):
    _seq_df  = pd.read_csv(seq_cache, sep='\t')
    sequences = dict(zip(_seq_df['Accession'], _seq_df['Sequence']))
    print(f"Loaded {len(sequences)} sequences from '{seq_cache}'.")
else:
    print(
        "Protein sequences not found locally. Downloading the required data from UniProt...\n"
        "This may take a few seconds on the first run, after which the data will be saved for future use.\n"
    )

# ============================================================
# Fetch sequences and build DataFrame
# ============================================================
    
    _rows = []
    for acc, ver in acc_versions.items():
        _r = requests.get(
            f"https://rest.uniprot.org/unisave/{acc}?format=fasta&versions={ver}",
            timeout=30,
        )
        _r.raise_for_status()
        _fa = _r.text
        _seq = ''.join(line for line in _fa.splitlines() if not line.startswith('>'))
        _rows.append({'Accession': acc, 'Sequence': _seq})
        print(f"  Fetched {acc} v{ver} ({len(_seq)} aa)")

# ============================================================
# Save to cache
# ============================================================

    _seq_df   = pd.DataFrame(_rows)
    _seq_df.to_csv(seq_cache, sep='\t', index=False)
    sequences = dict(zip(_seq_df['Accession'], _seq_df['Sequence']))
    print(f"\nSaved '{seq_cache}' with {len(sequences)} sequences.")


# %% [code]
## Reproducibility Module 2. Download UniProt manually curated PTM data for all subunits of RNA Pol II

# ============================================================
# Imports
# ============================================================

import pandas as pd
import pathlib as _pl
import requests

# ============================================================
# Download from UniProt
# ============================================================

_ptm_cache = sorted(_pl.Path('.').glob('ptms_polII_*.tsv'))
if _ptm_cache:
    uniprot_curated_all = pd.DataFrame(columns=['Substrate', 'Residue', 'PTM'])
    print(f"Cache found: {_ptm_cache[-1].name}: skipping UniProt curated download.")
else:

# ============================================================
# Configuration
# ============================================================

    uniprot_acs = [
        'P24928', 'P30876', 'P19387', 'O15514', 
        'P19388', 'P61218', 'P62487', 'P52434', 
        'P36954', 'P62875', 'P52435', 'P53803'
    ]

# ============================================================
# Fetch PTM features per accession
# ============================================================

    all_uniprot_ptms = []

    for ac in uniprot_acs:
        url = f"https://rest.uniprot.org/uniprotkb/{ac}.json?fields=ft_mod_res"
        response = requests.get(url, timeout=30)

        if response.status_code == 200:
            data = response.json()

            features = data.get('features', [])

            extracted_data = []
            for feature in features:
                ft_type = feature.get('type')
                if ft_type == 'Modified residue':
                    location = feature.get('location', {})
                    start = location.get('start', {}).get('value')

                    # Check if it's a single exact position (start == end usually for point mods)
                    if start and isinstance(start, int):
                        # Description contains the modification name
                        description = feature.get('description', '')

                        # Some descriptions have extra details separated by semicolon, e.g. "Phosphoserine; by..."
                        mod_name = description.split(';')[0].strip()

                        residue_letter = sequences[ac][start - 1] if 0 < start <= len(sequences[ac]) else ''

                        extracted_data.append({
                            'Substrate': ac,
                            'Residue': f"{residue_letter}{start}",
                            'PTM': mod_name
                        })

            if extracted_data:
                df = pd.DataFrame(extracted_data)
                df.drop_duplicates(inplace=True)
                all_uniprot_ptms.append(df)
                print(f"[{ac}] Successfully extracted {len(df)} modifications.")
            else:
                print(f"[{ac}] No PTM features found.")
        else:
            print(f"[{ac}] Failed to fetch. Status code: {response.status_code}")

# ============================================================
# Combine and report
# ============================================================

    if all_uniprot_ptms:
        uniprot_curated_all = pd.concat(all_uniprot_ptms, ignore_index=True)
        print(f"\nSuccessfully extracted {len(uniprot_curated_all)} total PTMs from UniProt manually curated PTM database.")
    else:
        print("No data was extracted from UniProt manually curated PTM database.")

# %% [code]
## Reproducibility Module 3. Download UniProt large-scale proteomics PTM data (EBI Proteins API) for all subunits of RNA Pol II

# ============================================================
# Imports
# ============================================================

import pandas as pd
import pathlib as _pl
import requests

# ============================================================
# Download from EBI Proteins API
# ============================================================

_ptm_cache = sorted(_pl.Path('.').glob('ptms_polII_*.tsv'))
if _ptm_cache:
    uniprot_large_scale_proteomics_all = pd.DataFrame(columns=['Substrate', 'Residue', 'PTM'])
    print(f"Cache found: {_ptm_cache[-1].name}: skipping UniProt large-scale download.")
else:

# ============================================================
# Configuration
# ============================================================

    uniprot_acs = [
        'P24928', 'P30876', 'P19387', 'O15514', 
        'P19388', 'P61218', 'P62487', 'P52434', 
        'P36954', 'P62875', 'P52435', 'P53803'
    ]

# ============================================================
# Fetch large-scale proteomics PTM features per accession
# ============================================================

    all_large_scale_proteomics_ptms = []

    for ac in uniprot_acs:
        url = f"https://www.ebi.ac.uk/proteins/api/proteomics/ptm/{ac}?format=json"
        response = requests.get(url, headers={"Accept": "application/json"}, timeout=30)

        if response.status_code == 200:
            data = response.json()

            extracted_data = []
            # Support both 'features' inside dict or inside the first element of a list
            if isinstance(data, list) and len(data) > 0:
                features = data[0].get('features', [])
            elif isinstance(data, dict):
                features = data.get('features', [])
            else:
                features = []

            for feature in features:
                if feature.get('type') == 'PROTEOMICS_PTM':
                    begin = int(feature.get('begin'))
                    peptide = feature.get('peptide', '')

                    ptms = feature.get('ptms', [])
                    for ptm in ptms:
                        name = ptm.get('name')
                        # 'position' is 1-based relative to the peptide string!
                        pos_in_peptide = int(ptm.get('position'))

                        if 1 <= pos_in_peptide <= len(peptide):
                            residue_letter = peptide[pos_in_peptide - 1].upper()
                            global_pos = begin + pos_in_peptide - 1

                            extracted_data.append({
                                'Substrate': ac,
                                'Residue': f"{residue_letter}{global_pos}",
                                'PTM': name
                            })

            if extracted_data:
                df = pd.DataFrame(extracted_data)
                df.drop_duplicates(inplace=True)
                all_large_scale_proteomics_ptms.append(df)
                print(f"[{ac}] Successfully extracted {len(df)} unique modifications.")
            else:
                print(f"[{ac}] No large-scale PTM features found.")
        else:
            print(f"[{ac}] No PTM data available in proteomics dataset.")
            
# ============================================================
# Combine and report
# ============================================================

    if all_large_scale_proteomics_ptms:
        uniprot_large_scale_proteomics_all = pd.concat(all_large_scale_proteomics_ptms, ignore_index=True)
        print(f"\nSuccessfully extracted {len(uniprot_large_scale_proteomics_all)} total PTMs from UniProt large-scale proteomics PTM database.")
    else:
        print("No data was extracted from EBI Proteins API.")

# %% [code]
## Reproducibility Module 4. Download dbPTM PTM data for all subunits of RNA Pol II

# ============================================================
# Imports
# ============================================================

import pandas as pd
import pathlib as _pl
import io
import requests
from bs4 import BeautifulSoup

# ============================================================
# Download from dbPTM
# ============================================================

_ptm_cache = sorted(_pl.Path('.').glob('ptms_polII_*.tsv'))
if _ptm_cache:
    dbPTM_all = pd.DataFrame(columns=['Substrate', 'Residue', 'PTM'])
    print(f"Cache found: {_ptm_cache[-1].name}: skipping dbPTM download.")
else:

# ============================================================
# Configuration
# ============================================================

    uniprot_acs = [
        'P24928', 'P30876', 'P19387', 'O15514', 
        'P19388', 'P61218', 'P62487', 'P52434', 
        'P36954', 'P62875', 'P52435', 'P53803'
    ]

    base_url = "https://biomics.lab.nycu.edu.tw/dbPTM/download/experiment"

# ============================================================
# Discover and fetch .gz files from directory index
# ============================================================

    index_response = requests.get(f"{base_url}/", timeout=30)
    index_response.raise_for_status()
    index_soup = BeautifulSoup(index_response.text, 'html.parser')
    gz_files = [a['href'] for a in index_soup.find_all('a', href=True) if a['href'].endswith('.gz')]
    print(f"Found {len(gz_files)} .gz files to process.")

    all_dfs = []

    for filename in gz_files:
        url = f"{base_url}/{filename}"
        try:
            _g = requests.get(url, timeout=60)
            _g.raise_for_status()
            df = pd.read_csv(
                io.BytesIO(_g.content), sep='\t', compression='gzip',
                header=None, on_bad_lines='skip',
                names=['dbptm_id', 'Substrate', 'Position', 'PTM', 'PubMed', 'Peptide']
            )
            filtered = df[df['Substrate'].isin(uniprot_acs)].copy()

            if not filtered.empty:
                # The flanking sequence is a 21-aa window; the modified residue is the middle character (index 10)
                filtered['Residue'] = (
                    filtered['Peptide'].str[10].str.upper() +
                    filtered['Position'].astype(int).astype(str)
                )
                all_dfs.append(filtered[['Substrate', 'Residue', 'PTM']])
                print(f"[{filename}] Found {len(filtered)} matching rows.")
            else:
                print(f"[{filename}] No matching rows for canonical RNA Pol II subunits.")
        except Exception as e:
            print(f"[{filename}] Failed: {e}")

# ============================================================
# Combine and report
# ============================================================

    if all_dfs:
        dbPTM_all = pd.concat(all_dfs, ignore_index=True)
        dbPTM_all.drop_duplicates(inplace=True)
        dbPTM_all.reset_index(drop=True, inplace=True)
        print(f"\nSuccessfully extracted {len(dbPTM_all)} total PTMs from dbPTM.")
    else:
        print("No data was extracted from dbPTM.")


# %% [code]
## Reproducibility Module 5. Download iPTMnet PTM data for all subunits of RNA Pol II

# ============================================================
# Imports
# ============================================================

import pandas as pd
import pathlib as _pl

# ============================================================
# Download from iPTMnet
# ============================================================

_ptm_cache = sorted(_pl.Path('.').glob('ptms_polII_*.tsv'))
if _ptm_cache:
    iptmnet_all = pd.DataFrame(columns=['Substrate', 'Residue', 'PTM'])
    print(f"Cache found: {_ptm_cache[-1].name}: skipping iPTMnet download.")
else:

# ============================================================
# Configuration
# ============================================================

    uniprot_acs = [
        'P24928', 'P30876', 'P19387', 'O15514', 
        'P19388', 'P61218', 'P62487', 'P52434', 
        'P36954', 'P62875', 'P52435', 'P53803'
    ]

    source = "https://research.bioinformatics.udel.edu/iptmnet_data/files/current/score.txt"

# ============================================================
# Fetch, filter and normalize
# ============================================================

    # Load without headers. Column 0 is Accession, 1 is Residue, 3 is PTM.
    iptmnet_df = pd.read_csv(source, sep='\t', header=None, usecols=[0, 1, 3], names=['Substrate', 'Residue', 'PTM'])

    # Filter only our subunits. We also include P24928-1 as iPTMnet stores it separately
    # even though its sequence is identical to the canonical sequence. Other subunits
    # do not have this issue as of the current data release.
    search_acs = uniprot_acs + ['P24928-1']
    iptmnet_all = iptmnet_df[iptmnet_df['Substrate'].isin(search_acs)].copy()

    # Normalize P24928-1 to P24928 to ensure uniformity downstream
    iptmnet_all['Substrate'] = iptmnet_all['Substrate'].replace('P24928-1', 'P24928')
    iptmnet_all = iptmnet_all[iptmnet_all['Residue'].astype(str).str.strip() != ''].copy()

    # Remove duplicates
    iptmnet_all.drop_duplicates(inplace=True)
    iptmnet_all.reset_index(drop=True, inplace=True)

# ============================================================
# Report
# ============================================================

    for ac in uniprot_acs:
        count = len(iptmnet_all[iptmnet_all['Substrate'] == ac])
        if count > 0:
            print(f"[{ac}] Successfully extracted {count} modifications.")
        else:
            print(f"[{ac}] No PTM features found.")
            
    print(f"\nSuccessfully extracted {len(iptmnet_all)} total PTMs from iPTMnet.")


# %% [code]
## Reproducibility Module 6. Download PhosphoSitePlus PTM data for all subunits of RNA Pol II

# ============================================================
# Imports
# ============================================================

import pandas as pd
import pathlib as _pl
import requests
from bs4 import BeautifulSoup
import re

# ============================================================
# Download from PhosphoSitePlus
# ============================================================

_ptm_cache = sorted(_pl.Path('.').glob('ptms_polII_*.tsv'))
if _ptm_cache:
    phosphosite_all = pd.DataFrame(columns=['Substrate', 'Residue', 'PTM'])
    print(f"Cache found: {_ptm_cache[-1].name}: skipping PhosphoSitePlus download.")
else:

# ============================================================
# Configuration
# ============================================================

    uniprot_map = {
        'P24928': 1039,
        'P30876': 14272,
        'P19387': 20249,
        'O15514': 14927,
        'P19388': 10246402,
        'P61218': 15349,
        'P62487': 3619563,
        'P52434': 13013,
        'P36954': 13186,
        'P62875': 9454,
        'P52435': 11965530,
        'P53803': 14553755
    }

    headers = {
        'User-Agent': 'Mozilla/5.0 (Windows NT 10.0; Win64; x64)'
    }

# ============================================================
# Fetch PTM sites per accession
# ============================================================

    all_dfs_psp = []

    for target_uniprot, target_id in uniprot_map.items():
        # 1. Fetch the main protein page to get the abbreviations
        main_url = f"https://www.phosphosite.org/proteinAction.action?id={target_id}&showAllSites=true"
        main_response = requests.get(main_url, headers=headers, timeout=30)

        abbreviation_map = {}
        if main_response.status_code == 200:
            main_soup = BeautifulSoup(main_response.text, 'html.parser')
            legend_div = main_soup.find('div', id='Show_Modification_Legend')
            if legend_div:
                legend_table = legend_div.find('table')
                if legend_table:
                    for row in legend_table.find_all('tr'):
                        spans = row.find_all('span', class_='popupInfoItem')
                        if len(spans) >= 2:
                            abbr = spans[0].text.strip()
                            full_name = spans[1].text.strip()
                            if abbr and full_name:
                                abbreviation_map[abbr] = full_name
        else:
            print(f"[{target_uniprot}] Failed to fetch main page {main_url}. Message: {main_response.text}")
            continue

        # 2. Fetch the site table
        site_url = f"https://www.phosphosite.org/siteTableNewAction?id={target_id}&showAllSites=true"
        site_response = requests.get(site_url, headers=headers, timeout=30)

        if site_response.status_code == 200:
            site_soup = BeautifulSoup(site_response.text, 'html.parser')
            tables = site_soup.find_all('table')

            if tables:
                extracted_data = []
                current_site = None

                for row in tables[0].find_all('tr'):
                    tds = row.find_all('td')
                    if not tds:
                        continue

                    first_col_text = tds[0].text.strip()

                    # Check if this row is a site header (e.g. S8-p)
                    match = re.match(r'^([A-Za-z]\d+)\-([a-z0-9]+)$', first_col_text)
                    if match:
                        residue = match.group(1).upper()
                        mod_abbr = match.group(2).lower()
                        current_site = {
                            'Residue': residue,
                            'PTM': abbreviation_map.get(mod_abbr, mod_abbr)
                        }
                    # Otherwise check if this row contains species info for the `current_site`
                    elif current_site and '(human)' in first_col_text.lower():
                        if 'iso' not in first_col_text.lower():
                            extracted_data.append({
                                'Substrate': target_uniprot,
                                'Residue': current_site['Residue'],
                                'PTM': current_site['PTM']
                            })

                result_df = pd.DataFrame(extracted_data)

                if not result_df.empty:
                    result_df.drop_duplicates(inplace=True)
                    all_dfs_psp.append(result_df)
                    print(f"[{target_uniprot}] Successfully extracted {len(result_df)} modifications.")
                else:
                    print(f"[{target_uniprot}] No matching '(human)' (non-iso) entries found.")

            else:
                print(f"[{target_uniprot}] Could not find the site table on PhosphoSitePlus.")
        else:
            print(f"[{target_uniprot}] Failed to fetch site table. Status code: {site_response.status_code}")

# ============================================================
# Combine and report
# ============================================================

    if all_dfs_psp:
        phosphosite_all = pd.concat(all_dfs_psp, ignore_index=True)
        print(f"\nSuccessfully extracted {len(phosphosite_all)} total PTMs from PhosphoSitePlus.")
    else:
        print("No data was extracted from PhosphoSitePlus.")

# %% [code]
## Reproducibility Module 7. Download GlyGen PTM data for all subunits of RNA Pol II

# ============================================================
# Imports
# ============================================================

import pandas as pd
import pathlib as _pl
import requests

# ============================================================
# Download from GlyGen
# ============================================================

_ptm_cache = sorted(_pl.Path('.').glob('ptms_polII_*.tsv'))
if _ptm_cache:
    glygen_all = pd.DataFrame(columns=['Substrate', 'Residue', 'PTM'])
    print(f"Cache found: {_ptm_cache[-1].name}: skipping GlyGen download.")
else:

# ============================================================
# Configuration
# ============================================================

    # The 12 exact accession codes (primarily isoform-1 where applicable)
    glygen_acs = [
        'P24928-1', 'P30876-1', 'P19387-1', 'O15514-1', 
        'P19388-1', 'P61218-1', 'P62487-1', 'P52434-1', 
        'P36954-1', 'P62875-1', 'P52435-1', 'P53803-1'
    ]

    # Standard 3-letter to 1-letter amino acid mapping to conform to the formatting ("S8", "K32", etc)
    aa_map = {
        'Ala': 'A', 'Arg': 'R', 'Asn': 'N', 'Asp': 'D', 'Cys': 'C', 
        'Gln': 'Q', 'Glu': 'E', 'Gly': 'G', 'His': 'H', 'Ile': 'I', 
        'Leu': 'L', 'Lys': 'K', 'Met': 'M', 'Phe': 'F', 'Pro': 'P', 
        'Ser': 'S', 'Thr': 'T', 'Trp': 'W', 'Tyr': 'Y', 'Val': 'V'
    }

# ============================================================
# Fetch glycosylation and phosphorylation features per accession
# ============================================================

    extracted_data_glygen = []

    for ac in glygen_acs:
        # Use the GlyGen protein detail endpoint
        url = f"https://api.glygen.org/protein/detail/{ac}/"
        response = requests.get(url, headers={"Accept": "application/json"}, timeout=30)

        if response.status_code == 200:
            data = response.json()

            # We normalize back the Accession removing the "-1" for downstream comparisons
            canonical_ac = ac.split("-")[0]
            rows_this_ac = []

            # 1. Look for Glycosylation data
            for item in data.get("glycosylation", []):
                pos = item.get("start_pos")
                residue = item.get("start_aa", "")
                subtype = item.get("subtype", "")

                if not subtype or subtype.lower() == "other":
                    continue

                if pos:
                    aa_letter = aa_map.get(residue, residue)
                    rows_this_ac.append((canonical_ac, f"{aa_letter}{pos}", subtype))

            # 2. Look for Phosphorylation data
            for item in data.get("phosphorylation", []):
                if item.get("site_category", "").lower() == "predicted":
                    continue

                pos = item.get("start_pos")
                residue = item.get("residue", "")

                if pos:
                    aa_letter = aa_map.get(residue, residue)
                    rows_this_ac.append((canonical_ac, f"{aa_letter}{pos}", "Phosphorylation"))

            unique_rows = set(rows_this_ac)
            extracted_data_glygen.extend(
                {'Substrate': s, 'Residue': r, 'PTM': p} for s, r, p in unique_rows
            )
            count = len(unique_rows)
            if count:
                print(f"[{canonical_ac}] Successfully extracted {count} modifications.")
            else:
                print(f"[{canonical_ac}] No PTM features found.")
        else:
            print(f"[{ac}] Failed to fetch. Status code: {response.status_code}")

# ============================================================
# Combine and report
# ============================================================

    if extracted_data_glygen:
        glygen_all = pd.DataFrame(extracted_data_glygen)
        glygen_all.drop_duplicates(inplace=True)
        glygen_all.reset_index(drop=True, inplace=True)
        print(f"\nSuccessfully extracted {len(glygen_all)} total PTMs from GlyGen.")
    else:
        print("No data was extracted from GlyGen.")

# %% [code]
## Reproducibility Module 8. CTD heptad position map (Rpb1, P24928)

# ============================================================
# Imports
# ============================================================

import re

# ============================================================
# Configuration
# ============================================================

# CTD region boundaries for Rpb1 (P24928)
CTD_START, CTD_END = 1593, 1970
TAIL_START = 1961

# 1-letter to 3-letter amino acid map for human-readable labels
aa3 = {
    'A': 'Ala', 'R': 'Arg', 'N': 'Asn', 'D': 'Asp', 'C': 'Cys',
    'Q': 'Gln', 'E': 'Glu', 'G': 'Gly', 'H': 'His', 'I': 'Ile',
    'L': 'Leu', 'K': 'Lys', 'M': 'Met', 'F': 'Phe', 'P': 'Pro',
    'S': 'Ser', 'T': 'Thr', 'W': 'Trp', 'Y': 'Tyr', 'V': 'Val'
}

# Map database PTM names to past-tense adjective form (for the state model string)
ptm_adjectives = {
    'Phosphorylation':           'phosphorylated',
    'Acetylation':               'acetylated',
    'Methylation':               'monomethylated',
    'Citrullination':            'citrullinated',
    'Dimethylation':             'dimethylated',
    'Trimethylation':            'trimethylated',
    'Asymmetric dimethylation':  'asymmetrically dimethylated',
    'Symmetric dimethylation':   'symmetrically dimethylated',
    'SUMOylation':               'SUMOylated',
    'Ubiquitination':            'ubiquitinated',
    'O-GlcNAcylation':           'O-GlcNAcylated',
    'O-Glycosylation':           'O-glycosylated',
    'N-Glycosylation':           'N-glycosylated',
    'Glutathionylation':         'glutathionylated',
    'Sulfoxidation':             'sulfoxidated',
    'Caspase cleavage':          'cleaved',
}

# ============================================================
# Validate CTD sequence start position
# ============================================================

sequence = sequences['P24928']
ctd_sequence = sequence[CTD_START - 1:]

assert ctd_sequence[0] == 'Y', (
    f"CTD position {CTD_START} should be Y, got '{ctd_sequence[0]}'; "
    f"check CTD_START."
)

# ============================================================
# Parse heptads and build position map
# ============================================================

# Heptad region only: slice up to the tail boundary so the tail (which has no
# leading Y) doesn't get glued onto heptad 52 by the Y-delimited split.
heptad_region = sequence[CTD_START - 1:TAIL_START - 1]   # protein positions 1593–1960, Python indexing is 0-based and end-exclusive
heptads = re.findall(r'Y[^Y]*', heptad_region)

# Build the position -> (heptad_index, position_within_heptad) map for the
# 52 canonical heptads. Residues beyond heptad 52 belongs to the C-terminal
# tail and are handled separately by the TAIL_START boundary downstream.
heptad_position_map = {}
running_pos = CTD_START
for h_idx, heptad in enumerate(heptads, start=1):
    for p_in_heptad in range(1, len(heptad) + 1):
        heptad_position_map[running_pos] = (h_idx, p_in_heptad)
        running_pos += 1

# ============================================================
# Validate and report
# ============================================================

assert len(heptads) == 52, f"Expected 52 heptads, got {len(heptads)}."
assert len(heptad_position_map) == 368, (
    f"Heptad map covers {len(heptad_position_map)} residues (expected 368). "
    f"Check CTD_START / TAIL_START."
)
print(f"Parsed {len(heptads)} Y-delimited heptads covering protein positions "
      f"{CTD_START}–{TAIL_START - 1}.")


# %% [code]
## Reproducibility Module 9. Build a side-by-side comparison of PTM names across all seven
# PTM data sources (UniProt curated, UniProt large-scale proteomics, dbPTM,
# iPTMnet, PhosphoSitePlus, GlyGen, and literature-mined)

# ============================================================
# Imports
# ============================================================

import pandas as pd
import pathlib as _pl
from collections import Counter
from functools import reduce

# ============================================================
# Build comparison
# ============================================================

_ptm_cache = sorted(_pl.Path('.').glob('ptms_polII_*.tsv'))
if _ptm_cache:
    ptm_comparison = pd.DataFrame()
    print(f"Cache found: {_ptm_cache[-1].name}: skipping database comparison.")
else:

# ============================================================
# Literature-mined PTMs
# ============================================================

    # ── Literature-mined PTMs not reported by the six databases ───────────────
    # (1) Pro cis/trans isomerization at:
    #       Pro3 in heptads YSPTSPS (Harding et al., 1992, doi:10.1021/jm00103a002 | Noble et al., 2005, doi:10.1038/nsmb887 | Gibbs et al., 2017, doi:10.1038/ncomms15233)
    #                   and YSPSSPS (Gibbs et al., 2017, doi:10.1038/ncomms15233)
    #                   and YSPTSPN (Gibbs et al., 2017, doi:10.1038/ncomms15233)
    #       Pro6 in heptads YSPTSPS (Xiang et al., 2010, doi:10.1038/nature09391 | Werner-Allen et al., 2011, doi:10.1074/jbc.M110.197129 | Zhang et al., 2012, doi:10.1021/cb3000887 | Mayfield et al., 2015, doi:10.1021/acschembio.5b00296 | Gibbs et al., 2017, doi:10.1038/ncomms15233)
    #                   and YSPTSPN (Gibbs et al., 2017, doi:10.1038/ncomms15233)
    # (2) O-GlcNAcylation at S1829 and S1896 (Lewis et al., 2016, doi:10.1074/jbc.M115.684365).
    # (3) Citrullination at R1810 (Sharma et al., 2019, doi:10.1016/j.molcel.2018.10.016).
    CONSENSUS_PRO3 = {'YSPTSPS', 'YSPSSPS', 'YSPTSPN'}
    CONSENSUS_PRO6 = {'YSPTSPS', 'YSPTSPN'}

    literature_rows = [
        {'Substrate': 'P24928', 'Residue': f'P{pos}', 'PTM': 'cis / trans isomerization'}
        for pos, (h_idx, hp) in heptad_position_map.items()
        if sequence[pos - 1] == 'P' and (
            (hp == 3 and heptads[h_idx - 1] in CONSENSUS_PRO3) or
            (hp == 6 and heptads[h_idx - 1] in CONSENSUS_PRO6)
        )
    ]

    literature_rows.extend([
        {'Substrate': 'P24928', 'Residue': 'S1829', 'PTM': 'O-GlcNAcylation'},
        {'Substrate': 'P24928', 'Residue': 'S1896', 'PTM': 'O-GlcNAcylation'},
        {'Substrate': 'P24928', 'Residue': 'R1810', 'PTM': 'Citrullination'},

    ])

    literature_mining_all = pd.DataFrame(literature_rows)

    assert len(literature_mining_all) == 54, (
        f"Expected 26 Pro3 (YSPTSPS|YSPSSPS|YSPTSPN) + 25 Pro6 (YSPTSPS|YSPTSPN) "
        f"+ 2 O-GlcNAc + 1 Arg1810 citrullination (Sharma et al. 2019) = 54 "
        f"literature-mined entries; got {len(literature_mining_all)}."
    )

# ============================================================
# Merge all sources into a single comparison table
# ============================================================

    sources = {
        'uniprot_curated':                 uniprot_curated_all,
        'uniprot_large_scale_proteomics':  uniprot_large_scale_proteomics_all,
        'dbPTM':                           dbPTM_all,
        'iptmnet':                         iptmnet_all,
        'phosphosite':                     phosphosite_all,
        'glygen':                          glygen_all,
        'literature_mining':               literature_mining_all
    }

    # For each source, collapse all PTM names that share the same (Substrate, Residue)
    # into a single "; "-joined string so the outer merge does not explode rows.
    agg_dfs = []
    for source_name, df in sources.items():
        grouped = (
            df.dropna(subset=['Substrate', 'Residue'])
              .astype({'Substrate': str, 'Residue': str, 'PTM': str})
              .groupby(['Substrate', 'Residue'], as_index=False)['PTM']
              .agg(lambda s: '; '.join(sorted(set(s))))
              .rename(columns={'Substrate': 'Accession', 'PTM': f'PTM_{source_name}'})
        )
        agg_dfs.append(grouped)

    # Outer-join on (Accession, Residue) so every site reported by ANY database
    # appears, with empty cells where a database did not annotate it.
    ptm_comparison = reduce(
        lambda left, right: pd.merge(left, right, on=['Accession', 'Residue'], how='outer'),
        agg_dfs
    ).fillna('')

    # Sort by Accession, then by the numeric position parsed out of the Residue label
    ptm_comparison['_pos'] = (
        ptm_comparison['Residue'].str.extract(r'(\d+)', expand=False).astype(int)
    )
    ptm_comparison = (
        ptm_comparison.sort_values(['Accession', '_pos'])
                      .drop(columns='_pos')
                      .reset_index(drop=True)
    )

# ============================================================
# Tally and report
# ============================================================

    # Tally the number of residues that have each unique combination of
    # PTM annotations across databases, and also the total number of residues
    # annotated with each individual PTM type.
    ptm_cols = ptm_comparison.columns[2:]  # all seven PTM_<source> columns

    combination_counts = Counter()   # residues per unique PTM combination
    ptm_site_counts    = Counter()   # residues per individual PTM type

    for _, row in ptm_comparison.iterrows():
        site_ptms = set()
        for cell_val in row[ptm_cols].dropna():
            if str(cell_val).strip():
                parts = [p.strip() for p in str(cell_val).split(';') if p.strip()]
                site_ptms.update(parts)
        if site_ptms:
            combination_counts['; '.join(sorted(site_ptms))] += 1
            for ptm in site_ptms:
                ptm_site_counts[ptm] += 1

    print("="*50)
    print(f"Found {len(combination_counts)} unique PTM combinations per residue "
          f"(sorted by descending instance count):\n")
    for combo, count in sorted(combination_counts.items(), key=lambda kv: (-kv[1], kv[0])):
        print(f" - {combo}: {count} residue{'s' if count != 1 else ''}")

    print("\n" + "="*50)
    print(f"Found {len(ptm_site_counts)} global, unique individual PTM types "
          f"(sorted by descending instance count):\n")
    for ptm, count in sorted(ptm_site_counts.items(), key=lambda kv: (-kv[1], kv[0])):
        print(f" - {ptm}: {count} residue{'s' if count != 1 else ''}")


# %% [code]
## Reproducibility Module 10. Database curation step.
# Create a merged long-form table of all unique (Accession, Residue, PTM) triples
# across all databases.

# ============================================================
# Imports
# ============================================================

import pandas as pd
from collections import defaultdict
import re
from collections import Counter
from datetime import date
import pathlib as _pl

# ============================================================
# Cached PTM atlas loading
# ============================================================

_ptm_cache = sorted(_pl.Path('.').glob('ptms_polII_*.tsv'))
if _ptm_cache:
    # Load from cache and skip full download+harmonization pipeline
    all_databases_merge = pd.read_csv(_ptm_cache[-1], sep='	')
    print(f"Loaded {len(all_databases_merge)} rows from cache: {_ptm_cache[-1].name}")
else:

# ============================================================
# Full PTM harmonization pipeline
# ============================================================

    # PTMs that are not reversible in a writer/eraser sense are excluded from
    # the residue-state model so that information capacity reflects only
    # bistable/regulatable sites.
    IRREVERSIBLE_PTMS = {'N-Glycosylation', 'Caspase cleavage'}

    ptm_comparison = (
        ptm_comparison
            .replace('Ubiquitinylation', 'Ubiquitination', regex=True)
            .replace('Ubiquitylation',   'Ubiquitination', regex=True)
            .replace('Sumoylation',   'SUMOylation', regex=True)
            .replace('Mono-methylation', 'Methylation',    regex=True)
            .replace('Di-methylation',   'Dimethylation',  regex=True)
            .replace(r'O-GlcNAc(?!ylation)', 'O-GlcNAcylation', regex=True)
            .replace('O-linked Glycosylation', 'O-Glycosylation', regex=True)
            # UniProt curated raw-name harmonization:
            .replace(r'Phospho(serine|threonine|tyrosine)', 'Phosphorylation', regex=True)
            .replace(r'N-acetyl(methionine|alanine|serine)', 'Acetylation',    regex=True)
            .replace('N6-acetyllysine',           'Acetylation',              regex=True)
            .replace('N6-methyllysine',           'Methylation',              regex=True)
            .replace('N6,N6-dimethyllysine',      'Dimethylation',            regex=True)
            .replace('N6,N6,N6-trimethyllysine',  'Trimethylation',           regex=True)
            .replace('Omega-N-methylarginine',    'Methylation',              regex=True)
            .replace('Omega-N-methylated arginine','Methylation',             regex=True)
            .replace('Asymmetric dimethylarginine','Asymmetric dimethylation',regex=True)
            .replace('Symmetric dimethylarginine','Symmetric dimethylation',  regex=True)
    )

    # Extract unique values across the PTM database columns (indices 2:end)
    ptm_cols = ptm_comparison.columns[2:]

    # Build the merged long-form table (one row per Accession, Residue, PTM) and tally statistics
    merged_rows = []

    for _, row in ptm_comparison.iterrows():
        accession = row['Accession']
        residue = row['Residue']

        site_ptms = set()
        for cell_val in row[ptm_cols].dropna():
            if str(cell_val).strip():
                # Split by semicolon because some single cells already contain
                # multiple PTMs
                parts = [p.strip() for p in str(cell_val).split(';') if p.strip()]
                site_ptms.update(parts)

        # Collapse O-Glycosylation into O-GlcNAcylation when both are present
        # at the same residue across databases and no other subtypes of O-Glycosylation
        # are annotated.
        if 'O-GlcNAcylation' in site_ptms and 'O-Glycosylation' in site_ptms:
            site_ptms.discard('O-Glycosylation')

        # Drop irreversible PTMs (no writer/eraser cycle) since they do not contribute
        # to the residue-state model used in the information-capacity calculation.
        site_ptms -= IRREVERSIBLE_PTMS

        if site_ptms:
            for ptm in sorted(site_ptms):
                merged_rows.append({'Accession': accession, 'Residue': residue, 'PTM': ptm})

    # Save the merged long-form TSV (Accession, Residue, PTM)
    all_databases_merge = pd.DataFrame(merged_rows)
    all_databases_merge = all_databases_merge.drop_duplicates().reset_index(drop=True)
    SNAPSHOT_DATE = date.today().strftime("%Y_%m_%d")   # ISO date format
    all_databases_merge.to_csv(f'ptms_polII_{SNAPSHOT_DATE}.tsv',
                               sep='\t', index=False)
    print(f"Frozen snapshot written: ptms_polII_{SNAPSHOT_DATE}.tsv")

# Post-curation: unique PTM combinations per residue and individual PTM type counts
# (mirrors the pre-curation output of the database comparison cell, for auditability)

_combo_counts = Counter()
for (_acc, _res), _grp in all_databases_merge.groupby(['Accession', 'Residue']):
    _combo_counts['; '.join(sorted(_grp['PTM'].unique()))] += 1

_ptm_site_counts = Counter(all_databases_merge['PTM'])  # already 1 row per (acc,res,ptm)

print("=" * 50)
print(f"Found {len(_combo_counts)} unique PTM combinations per residue "
      f"(sorted by descending instance count):\n")
for _combo, _cnt in sorted(_combo_counts.items(), key=lambda kv: (-kv[1], kv[0])):
    print(f"  - {_combo}: {_cnt} residue{'s' if _cnt != 1 else ''}")

print("\n" + "=" * 50)
print(f"Found {len(_ptm_site_counts)} global, unique individual PTM types "
      f"(sorted by descending instance count):\n")
for _ptm, _cnt in sorted(_ptm_site_counts.items(), key=lambda kv: (-kv[1], kv[0])):
    print(f"  - {_ptm}: {_cnt} residue{'s' if _cnt != 1 else ''}")
```
