## Supplementary material for "Multi-timescale PTM architecture in human RNA polymerase II": notebook_2.ipynb

```
# %% [markdown]
# Multi-timescale PTM architecture in human RNA polymerase II

## Notebook 2 — Information-theoretic calculations
---

This notebook (Supplementary Data File 2) reproduces all quantitative results of the manuscript and of Supplementary Information S1 and S3 from the frozen `uniprot_polII_seqs_2026_05_22.tsv` (Supplementary Data File 3) and `ptms_polII_2026_05_22.tsv` (Supplementary Data File 4) files provided in this repository. The manuscript-value asserts in this notebook are pinned to these two snapshots and will fail against new ones regenerated from updated upstream resources.
> [!NOTE]
> **The notebook `notebook_1.ipynb` (Supplementary Data File 1) need not be run beforehand if these two snapshot files are present in the working directory; it is required only to regenerate the snapshots or to audit their provenance**.

| File | Source | Role |
|------|--------|------|
| `uniprot_polII_seqs_2026_05_22.tsv` | UniProt UniSave (version-pinned canonical sequences) | Protein sequence reference for structural mapping |
| `ptms_polII_2026_05_22.tsv` | Integrated PTM atlas (UniProt, dbPTM, iPTMnet, PhosphoSitePlus, GlyGen, literature mining) | Combined residue-level PTM dataset |

If `ptms_polII_2026_05_22.tsv` is absent, this notebook cannot reconstruct the PTM atlas and downstream analyses will fail.

If `uniprot_polII_seqs_2026_05_22.tsv` is absent, structural partitioning (CTD/core/heptad mapping) cannot be performed and all region-specific analyses will be invalid.

This notebook derives all region partitions (CTD, core, Rpb2–Rpb12), structural mappings (heptads and tail), and information-theoretic quantities (I_CTD, I_core, I_r212, I_total, I_nucleus).

**Software requirements:** Python ≥ 3.10 | pandas ≥ 1.3 | IPython ≥ 7.0

---

## Reproducibility workflow overview

Full descriptions are provided in Supplementary Information S4. The table below is an informative index only.

The first cell in this notebook (***Environment check and analysis setup***) checks minimum software requirements (raises an error if unmet), loads frozen PTM atlas and UniProt sequences from the two .tsv supplemental files, and generates assets essential for downstream analysis.
> [!WARNING]
> **It is required to run this cell before the rest of the analysis**.

| Reproducibility Module | Stage | Contents |
|------|-------|----------|
| **11** | CTD PTM heptad map (Table 1, Results I section) | Constructs a color-coded map of Rpb1 (P24928) CTD PTM states across the 52 heptads and the C-terminal tail. |
| **12** | CTD state space, information capacity, and residue-class decomposition (Table 2, Results II section) | Computes total combinatorial state space size (N<sub>CTD</sub>) and information capacity (I<sub>CTD</sub> in bits), then structures PTM combinations into ordered residue-type classes, and computes per-class combinatorial state space size and information contribution (Table 2). |
| **13** | Information capacity encoded by PTMs in the Rpb1 non-CTD (core) sequence (Results III section) | Builds Supplementary Table S1 for the Rpb1 non-CTD region (residues 1–1592). Reports modified residues, PTM composition, number of accessible states (sᵢ), and information contribution (log₂(sᵢ)) for each modified residue. |
| **14** | Information capacity encoded by PTMs in subunits Rpb2–Rpb12 (Results IV section) | Builds supplementary tables S2-S12 for RNA polymerase II subunits Rpb2–Rpb12 (one per subunit), listing modified residues, PTM combinations, state-space size (sᵢ), and per-site information contribution (log₂(sᵢ)). It also reports information capacity for each subunit. |
| **15** | Upper bound theoretical information capacity per Pol II molecule and per nucleus (Results V and VI sections)| Combines the information capacities of the CTD, Rpb1 core, and Rpb2–Rpb12 subunits to estimate the total PTM information capacity of a single RNA polymerase II molecule (I<sub>total</sub>). Using an abundance of 80,200 Pol II molecules per nucleus (obtained from the literature), it further estimates the theoretical upper-bound PTM information capacity at the nuclear scale (I<sub>nucleus</sub>). |
| **16** | Multi-timescale PTM biochemical architecture (Results VII section) | Estimates a physiologically constrained PTM information capacity for RNA polymerase II by partitioning PTM regulation into literature-derived fast, intermediate, and slow biochemical layers. Computes layer-wise and total physiological information capacity (I<sub>physio</sub>) at the molecular and nuclear scales and compares these estimates against the theoretical maximum PTM capacity (I<sub>total</sub>). |
| **17** | Total unique PTM sites and modified residues in the integrated Pol II PTM database (Supplementary Information S1) | Computes total unique PTMs and total unique modified residues across the RNA Pol II molecule. |
| **18** | Sensitivity analyses (Supplementary Information S3) | Reproduces Supplementary Information S3 (Sections S3.1–S3.4) and Supplementary Information S3-Table S1. Quantifies the effects of applying four alternative PTM-state models (sensitivity scenarios S0–S3) at the molecular and nuclear scales: S0 (baseline), S1 (non-CTD Thr conservative), S2 (CTD Pro cis/trans excluded), S3 (maximally conservative). |

# %% [code]
## Environment check and analysis setup.
# Verifies that the Python environment meets the minimum requirements for running the notebook.
# Loads the reproducible input files (UniProt sequences and PTM atlas) and constructs the derived CTD state-space object for downstream analysis.

import sys, importlib, re
import pandas as pd
from IPython.display import display
from math import log2, log10, isclose
import pathlib
from collections import defaultdict

# ============================================================
#  ENVIRONMENT CHECK
# ============================================================
#  Python
# ============================================================

_py = sys.version_info
assert _py >= (3, 10), f"Python ≥ 3.10 required, got {_py.major}.{_py.minor}"

# ============================================================
# Third-party requirements
# ============================================================

_REQUIRED = {
    "pandas":   (1, 3),   # data manipulation
    "IPython":  (7, 0),   # display(), HTML()
}

def _parse(ver_str):
    """Numeric version tuple, ignoring pre-release suffixes (e.g. '1.3.0rc1')."""
    return tuple(int(x) for x in re.findall(r'\d+', ver_str.split('+')[0])[:3])

_ok, _fail = [], []
for _pkg, _min in _REQUIRED.items():
    try:
        _ver_str = importlib.import_module(_pkg).__version__
        _ver = _parse(_ver_str)
        _min_str = ".".join(map(str, _min))
        if _ver >= _min:
            _ok.append(f"  \u2713  {_pkg:<10} {_ver_str}")
        else:
            _fail.append(f"  \u2717  {_pkg:<10} {_ver_str}  (need \u2265 {_min_str})")
    except ImportError:
        _fail.append(f"  \u2717  {_pkg:<10} NOT INSTALLED  (need \u2265 {'.'.join(map(str, _min))})")

# ============================================================
# Report 
# ============================================================

print(f"  \u2713  {'Python':<10} {_py.major}.{_py.minor}.{_py.micro}")
print(*_ok, sep="\n")
if _fail:
    print("\n  FAILED:")
    print(*_fail, sep="\n")
    raise EnvironmentError("Environment check failed; install missing/outdated packages before running.")
else:
    print("\n  All requirements satisfied. Ready to run.")

# ============================================================
# ANALYSIS SETUP
# ============================================================
# Load reproducible inputs from .tsv snapshots
# ============================================================

SEQ_FILE = sorted(pathlib.Path(".").glob("uniprot_polII_seqs_*.tsv"))[-1]
PTM_FILE = sorted(pathlib.Path(".").glob("ptms_polII_*.tsv"))[-1]

assert pathlib.Path(SEQ_FILE).exists(), "Sequence file not found"
assert pathlib.Path(PTM_FILE).exists(), "PTM atlas file not found"

print()
print(f"Using sequence file: {SEQ_FILE}")
print()
print(f"Using PTM file: {PTM_FILE}")

sequences_df = pd.read_csv(SEQ_FILE, sep="\t")
sequences = dict(zip(sequences_df["Accession"], sequences_df["Sequence"]))
all_databases_merge = pd.read_csv(PTM_FILE, sep="\t")

print()
print(f"Loaded sequences: {len(sequences_df)} entries")

# ============================================================
# CTD structural definition
# ============================================================

CTD_START  = 1593
CTD_END    = 1970
TAIL_START = 1961

sequence = sequences["P24928"]

heptad_region = sequence[CTD_START - 1:TAIL_START - 1]
heptads = re.findall(r"Y[^Y]*", heptad_region)

heptad_position_map = {}
running_pos = CTD_START

for h_idx, heptad in enumerate(heptads, start=1):
    for p_in_heptad in range(1, len(heptad) + 1):
        heptad_position_map[running_pos] = (h_idx, p_in_heptad)
        running_pos += 1

print()
print(f"Reconstructed {len(heptads)} CTD heptads")

# ============================================================
# Derived CTD state-space object 
# ============================================================

# 1-letter to 3-letter amino acid map for human-readable labels
aa3 = {
    'A': 'Ala', 'R': 'Arg', 'N': 'Asn', 'D': 'Asp', 'C': 'Cys',
    'Q': 'Gln', 'E': 'Glu', 'G': 'Gly', 'H': 'His', 'I': 'Ile',
    'L': 'Leu', 'K': 'Lys', 'M': 'Met', 'F': 'Phe', 'P': 'Pro',
    'S': 'Ser', 'T': 'Thr', 'W': 'Trp', 'Y': 'Tyr', 'V': 'Val'
}

# Map database PTM names to past-tense adjective form (for the state model string)
ptm_adjectives = {
    'Phosphorylation':           'phosphorylated',
    'Acetylation':               'acetylated',
    'Methylation':               'monomethylated',
    'Citrullination':            'citrullinated',
    'Dimethylation':             'dimethylated',
    'Trimethylation':            'trimethylated',
    'Asymmetric dimethylation':  'asymmetrically dimethylated',
    'Symmetric dimethylation':   'symmetrically dimethylated',
    'SUMOylation':               'SUMOylated',
    'Ubiquitination':            'ubiquitinated',
    'O-GlcNAcylation':           'O-GlcNAcylated',
    'O-Glycosylation':           'O-glycosylated',
    'N-Glycosylation':           'N-glycosylated',
    'Glutathionylation':         'glutathionylated',
    'Sulfoxidation':             'sulfoxidated',
    'Caspase cleavage':          'cleaved',
}

def _build_ctd_subclass(_df):
    """Canonical CTD subclass tally for this notebook (single implementation).
    Operates on the final long-form (Accession, Residue, PTM) table."""
    _sub = defaultdict(lambda: {'count': 0, 'positions': []})
    for (acc, res), grp in _df.groupby(['Accession', 'Residue']):
        if acc != 'P24928':
            continue
        _pm = re.search(r'(\d+)', str(res))
        if not _pm:
            continue
        pos = int(_pm.group(1))
        if not (CTD_START <= pos <= CTD_END):
            continue
        site_ptms = set(grp['PTM'])
        aa = res[0]
        h_idx, heptad_pos = heptad_position_map.get(pos, (None, None))
        if heptad_pos is None:
            label_base = f"{aa3.get(aa, aa)}?"
        elif heptad_pos <= 7:
            label_base = f"{aa3.get(aa, aa)}{heptad_pos}"
        else:
            label_base = f"{aa3.get(aa, aa)}{heptad_pos} (heptad {h_idx})"
        if site_ptms == {'cis / trans isomerization'}:
            state_model = 'cis / trans isomerization'
        else:
            state_model = ' / '.join(
                ['unmodified'] +
                sorted(ptm_adjectives.get(p, p.lower()) for p in site_ptms)
            )
        key = (label_base, state_model)
        _sub[key]['count'] += 1
        _sub[key]['positions'].append(pos)
    return _sub

ctd_subclass = _build_ctd_subclass(all_databases_merge)

# ============================================================
# Region partitions
# ============================================================

_pos  = all_databases_merge['Residue'].str.extract(r'(\d+)', expand=False).astype(int)
_rpb1 = all_databases_merge['Accession'] == 'P24928'

ctd_df  = all_databases_merge[_rpb1 & _pos.between(CTD_START, CTD_END)]
core_df = all_databases_merge[_rpb1 & _pos.lt(CTD_START)]
r212_df = all_databases_merge[~_rpb1]

# ============================================================
# Information capacity function
# ============================================================

def _I(df):
    """I = Σ log2(n_ptm_types + 1) over unique (Accession, Residue) sites."""
    if df.empty:
        return 0.0

    return float(
        df.groupby(['Accession', 'Residue'])['PTM']
          .nunique()
          .add(1) # For the unmodified baseline state
          .apply(log2)
          .sum()
    )

# %% [code]
## Reproducibility Module 11 - Color-coded CTD PTM heptad map (Table 1).

# ============================================================
# Imports and sequence tail extraction
# ============================================================

from IPython.display import HTML
from math import log10, isclose, prod

tail = sequence[TAIL_START - 1 : CTD_END]

# ============================================================
# PTM mapping at residue level (CTD only)
# ============================================================

ptm_by_pos = (
    ctd_df.groupby(ctd_df['Residue'].str.extract(r'(\d+)', expand=False).astype(int))['PTM']
    .apply(frozenset).to_dict()
)

# ============================================================
# PTM combinatorial state space construction
# ============================================================

_combos  = sorted({v for v in ptm_by_pos.values()}, key=lambda s: (len(s), sorted(s)))
_palette = ["#0022ff","#ff0d00","#00C10A",'#9900ff',
            '#2fbeef','#FF10F0','#FFD700',"#AD7600"]

PAL = {
    combo: (_palette[i % len(_palette)], ' / '.join(sorted(combo)))
    for i, combo in enumerate(_combos)
}

# ============================================================
# Visualization helpers (color encoding of PTM states)
# ============================================================

UNMODIFIED_COLOR = '#000000'

def ptm_to_key(ptms):
    k = frozenset(ptms)
    return k if k in PAL else None

def colored(aa, key):
    if key:
        return f'<span style="color:{PAL[key][0]};font-weight:bold">{aa}</span>'
    return aa

def color_heptad(hep, start, flag=''):
    return ''.join(
        colored(a, ptm_to_key(ptm_by_pos.get(start + j, frozenset())) if j < 7 else None)
        for j, a in enumerate(hep)
    ) + flag

def color_tail(t):
    return ''.join(
        colored(a, ptm_to_key(ptm_by_pos.get(TAIL_START + j, frozenset())))
        for j, a in enumerate(t)
    )

rows, pos = [], CTD_START
for i, h in enumerate(heptads):
    flag = '*' if len(h) > 7 else ''
    rows.append((str(i+1), str(pos), color_heptad(h, pos, flag),
                 'Consensus' if h == 'YSPTSPS' else 'Nonconsensus'))
    pos += len(h)
rows.append(('Tail', str(TAIL_START), color_tail(tail), 'Terminal tail (10 amino acids)'))

mid = (len(rows) + 1) // 2
left, right = rows[:mid], rows[mid:]
header = ''.join(f'<th>{c}</th>' for c in ['n', 'Start', 'Heptad', 'Class'] * 2)
body   = '\n'.join(
    '<tr>' + ''.join(f'<td>{v}</td>' for v in left[i]) +
    (''.join(f'<td>{v}</td>' for v in right[i]) if i < len(right) else '<td></td>'*4) + '</tr>'
    for i in range(mid)
)

# ============================================================
# Legend construction
# ============================================================

unmod_legend = f'<b style="font-size:10pt;"><span style="color:{UNMODIFIED_COLOR};">■</b> unmodified</span>'
ptm_legend = ', '.join(f'<b style="color:{c};font-size:10pt;">■</b> <span style="font-size:10pt;">{l}</span>' for c, l in PAL.values())
legend = unmod_legend + ', ' + ptm_legend

# ============================================================
# HTML rendering
# ============================================================

html = f"""<div id="ctd-tbl"><style>
  #ctd-tbl {{font-family:Calibri,sans-serif;font-size:10pt}}
  #ctd-tbl table {{border-collapse:collapse;margin-top:7px}}
  #ctd-tbl th,#ctd-tbl td {{border:1px solid #bbb;padding:3px 5px;text-align:center;
          background:transparent;color:inherit}}
  #ctd-tbl tr:nth-child(even),#ctd-tbl tr:nth-child(odd) {{background:transparent!important}}
  #ctd-tbl th    {{font-weight:bold!important}}
  #ctd-tbl td:nth-child(1),#ctd-tbl td:nth-child(5) {{font-weight:bold}}
  #ctd-tbl td:nth-child(3),#ctd-tbl td:nth-child(7) {{font-size:13pt}}
</style>
  <table><tr>{header}</tr>{body}</table>
  <p style="font-size:10pt;"><b>n</b> represents the heptamer number; Start represents the position of the first amino acid in the heptamer.
  *Two of the heptamers include additional residues (heptamer n=2 features an extra Gly, and heptamer n=49 an extra Gly-Ser-Thr). 
  In both cases, none of these extra amino acid residues include PTMs³⁷⁻⁴¹. The color code corresponds to the following PTM classes: {legend}.</p>
</div>"""

print('Table 1. CTD structure comprising 52 heptapeptides and a 10-residue C-terminal tail (UniProt P24928 canonical sequence).')
display(HTML(html))


# %% [code]
## Reproducibility Module 12 - CTD combinatorial state space size, upper-bound theoretical information capacity,
# and residue-class decomposition of CTD PTM patterns (Table 2 of the manuscript main text).
# Construction of CTD PTM classes from heptad and C-terminal tail regions.
# Defines residue-type–ordered state classes derived from CTD subclasses.
# Computes per-class combinatorial state space size (sᵢ) and information contributions (log₂(sᵢ), total bits).
# Includes consistency audit ensuring that total information equals I_CTD.

# ============================================================
# Imports
# ============================================================

import math
import re
from collections import defaultdict
from math import isclose
import pandas as pd
from IPython.display import display

# ============================================================
# Information capacity computation and audit
# ============================================================

_n_states   = ctd_df.groupby(["Accession", "Residue"])["PTM"].nunique().add(1) # add 1 for the baseline unmodified state
N_CTD       = prod(int(x) for x in _n_states)
I_CTD  = _I(ctd_df)
log10_N_CTD = I_CTD * log10(2)
print(f"Combinatorial state space size of Rpb1 CTD ≈ {N_CTD:.2e} states (≈ 10^{log10_N_CTD:.2f})")

assert isclose(log10_N_CTD, 63.58, abs_tol=0.005), (
    f"AUDIT FAIL: log10(N_CTD) = {log10_N_CTD:.2f}  (expected 63.58)"
)

print(f"\n  ✓  AUDIT  log10(N_CTD) = {log10_N_CTD:.2f}")

# ============================================================
# Global CTD information capacity audit
# ============================================================

print(f"\nMaximum Information capacity of Rpb1 CTD = {I_CTD:.2f} bits")

assert isclose(I_CTD, 211.22, abs_tol=0.005), (
    f"AUDIT FAIL: I_CTD = {I_CTD:.2f} bits  (expected 211.22 bits)"
)
print(f"\n  ✓  AUDIT  I_CTD = {I_CTD:.2f} bits")
print()

# ============================================================
# CTD PTM inventory
# ============================================================

ctd_ptm_types = sorted(ctd_df["PTM"].unique())
print("Unique PTM types in CTD (residues 1593–1970):")
for _t in ctd_ptm_types:
    print(f"  - {_t}")

_n_sites_ctd = ctd_df[["Accession", "Residue"]].drop_duplicates().shape[0]
_n_pairs_ctd = ctd_df[["Accession", "Residue", "PTM"]].drop_duplicates().shape[0]
print()
print(f"  Modified residues (unique sites):     {_n_sites_ctd}")
print(f"  Unique (residue, PTM type) pairs:     {_n_pairs_ctd}")
print()

# ============================================================
# Residue ordering (heptad biological structure)
# ============================================================

# Residue-type appearance order along a canonical YSPTSPS heptad,
# followed by common non-canonical position-7 residues, then the rest alphabetically
RESIDUE_GROUP_ORDER = [
    'Tyr', 'Ser', 'Pro', 'Thr',  # canonical heptad order Y-S-P-T-S-P-S
    'Lys', 'Arg',                # common non-canonical position-7 residues
    'Ala', 'Asn', 'Asp', 'Cys', 'Gln', 'Glu',
    'Gly', 'His', 'Ile', 'Leu', 'Met', 'Phe', 'Trp', 'Val',
]

# ============================================================
# Partition CTD subclasses into heptad vs tail regions
# ============================================================

# Split each subclass entry into heptad-region positions vs C-terminal-tail positions.
heptad_subclasses = {}                                                  # {(label_base, combo): data}
tail_subclasses = defaultdict(lambda: {'count': 0, 'positions': []})    # {(residue_type, combo): data}

for (label_base, combo), data in ctd_subclass.items():
    non_tail_positions = [p for p in data['positions'] if p <  TAIL_START]
    tail_positions     = [p for p in data['positions'] if p >= TAIL_START]

    if non_tail_positions:
        heptad_subclasses[(label_base, combo)] = {
            'count':     len(non_tail_positions),
            'positions': non_tail_positions,
        }
    if tail_positions:
        residue_type = re.match(r'^([A-Za-z]+)', label_base).group(1)
        key = (residue_type, combo)
        tail_subclasses[key]['count']     += len(tail_positions)
        tail_subclasses[key]['positions'].extend(tail_positions)


# ============================================================
# Construct CTD PTM classes (ordered decomposition)
# ============================================================

# Build ptm_classes in the requested order
ptm_classes = []

LABEL_OVERRIDES = {
    'Pro3': 'Pro3 (YSPTSPS, YSPSSPS & YSPTSPN)',
    'Pro6': 'Pro6 (YSPTSPS & YSPTSPN)',
}

for residue_type in RESIDUE_GROUP_ORDER:
    # All heptad-region subclasses of this residue type, sorted by minimum position
    type_rows = sorted(
        [(label_base, combo, d) for (label_base, combo), d in heptad_subclasses.items()
         if label_base.startswith(residue_type)],
        key=lambda x: (int(re.search(r'\d+', x[0]).group()), min(x[2]['positions']))
    )
    for label_base, combo, d in type_rows:
        count = d['count']
        label = LABEL_OVERRIDES.get(label_base,
                f"{label_base} ({d['positions'][0]})" if count == 1 else label_base)
        s = combo.count(' / ') + 1
        ptm_classes.append((label, combo, count, s))

# ====== C-terminal tail rows go LAST ======
# One row per (residue_type, observed combo); the label lists every tail-relative
# position sharing that combo, e.g. "Ser2, Ser6 C-terminal tail".
for (residue_type, combo), d in sorted(
    tail_subclasses.items(), key=lambda kv: min(kv[1]['positions'])
):
    tail_positions_relative = sorted(p - TAIL_START + 1 for p in d['positions'])
    label = ", ".join(f"{residue_type}{tp}" for tp in tail_positions_relative) + " C-terminal tail"
    s = combo.count(' / ') + 1
    ptm_classes.append((label, combo, d['count'], s))

# ============================================================
# Table construction (information decomposition)
# ============================================================

headers = ['Residue/site class', 'PTM state model', 'Count',
           'States per site (sᵢ)', 'Bits/site log₂(sᵢ)', 'Total bits']

rows = []
for site, combination, count, s in ptm_classes:
    b = math.log2(s) if s > 1 else 0.0
    rows.append([site, combination, count, s, b, count * b])

df_table2 = pd.DataFrame(rows, columns=headers)

# ============================================================
# Audit: consistency with global CTD information
# ============================================================

# Cross-check: Table 2 "Total bits" column total must equal I_CTD from manuscript.
_nb_col = "Total bits"
assert isclose(df_table2[_nb_col].sum(), I_CTD, abs_tol=0.005), (
    f"AUDIT FAIL: Table 2 Total bits = {df_table2[_nb_col].sum():.2f} bits, "
    f"expected I_CTD = {I_CTD:.2f} bits"
)
print(f"  ✓  AUDIT  Table 2 Total bits = {df_table2[_nb_col].sum():.2f} bits  (= I_CTD)")

# ============================================================
# Render Table 2
# ============================================================

styled_table = (
    df_table2.style
    .hide(axis="index")
    .format({
        headers[4]: "{:.2f}",
        headers[5]: "{:.2f}"
    })
    .set_properties(**{
        'font-size': '11pt',
        'font-family': 'Calibri, sans-serif',
        'font-weight': 'normal',
        'text-align': 'center',
        'border': '1px solid #bbb'
    })
    .set_properties(subset=['Residue/site class'], **{'font-weight': 'bold'})
    .set_table_styles([
        {'selector': 'table', 'props': [('border-collapse', 'collapse')]},
        {'selector': 'th', 'props': [('font-size', '11pt'),
                                     ('font-family', 'Calibri, sans-serif'),
                                     ('text-align', 'center'),
                                     ('border', '1px solid #bbb')]},
        {'selector': 'tbody tr:nth-child(even)', 'props': [('background-color', 'transparent')]},
        {'selector': 'tbody tr:nth-child(odd)',  'props': [('background-color', 'transparent')]},
        {'selector': 'td', 'props': [('background-color', 'transparent')]},
        {'selector': 'th', 'props': [('background-color', 'transparent')]}
    ])
)

print("\nTable 2. Residue-level information and contribution per PTM of the CTD (Rpb1 subunit).")
display(styled_table)

# %% [code]
## Reproducibility Module 13 - Upper-bound theoretical information capacity of PTMs on the Rpb1 core (non-CTD).
# Construction of Supplementary Table S1 for the Rpb1 core region (residues 1–1592).
# Defines residue-level PTM state models for the Rpb1 non-CTD region and
# quantifies their information content via log₂(sᵢ).
# I_core is validated against the manuscript value.
# Per-site decomposition is validated against the global core capacity (I_core) via audit.

# ============================================================
# Imports
# ============================================================
from math import log2, isclose
import pandas as pd
from IPython.display import display

# ============================================================
# Global core information capacity
# ============================================================

I_core = _I(core_df)

# Unique PTM types in RPB1 core
core_ptm_types = sorted(core_df["PTM"].unique())
print("Unique PTM types in RPB1 core (residues 1-1592):")
for _t in core_ptm_types:
    print(f"  - {_t}")

_n_sites = core_df[["Accession", "Residue"]].drop_duplicates().shape[0]
_n_pairs = core_df[["Accession", "Residue", "PTM"]].drop_duplicates().shape[0]
print()
print(f"  Modified residues (unique sites):     {_n_sites}")
print(f"  Unique (residue, PTM type) pairs:     {_n_pairs}")

# ============================================================
# Core state-space construction (per-residue PTM models)
# ============================================================

# Build Supplementary Table S1 POLR2A (RPB1 core, non-CTD)
def _core_res_pos(r):
    try: return int(r[1:])
    except: return 0

_rows_s1c = []
for (_acc, _res), _grp in core_df.groupby(["Accession", "Residue"]):
    _ptms = sorted(_grp["PTM"].unique())
    _si   = len(_ptms) + 1
    _rows_s1c.append({
        "Residue":                 _res,
        "PTM Type(s)":             " / ".join(_ptms),
        "UniProt accession":       _acc,
        "States (sᵢ)":          _si,
        "Bits per site log₂(sᵢ)": log2(_si),
    })

_df_s1c = (
    pd.DataFrame(_rows_s1c)
    .assign(_pos=lambda _d: _d["Residue"].apply(_core_res_pos))
    .sort_values("_pos")
    .drop(columns="_pos")
    .reset_index(drop=True)
)[["Residue", "PTM Type(s)", "UniProt accession",
   "States (sᵢ)", "Bits per site log₂(sᵢ)"]]

# ============================================================
# Audit: consistency of table-derived data with manuscript value and with I_core
# ============================================================

print()
assert isclose(I_core, 185.87, abs_tol=0.005), (
    f"  AUDIT FAIL: I_core = {I_core:.2f} bits (expected 185.87)"
)
print(f"  ✓  AUDIT  I_core = {I_core:.2f} bits, matches manuscript value")

_bits_sum = _df_s1c["Bits per site log₂(sᵢ)"].sum()
assert isclose(_bits_sum, I_core, abs_tol=0.005), (
    f"AUDIT FAIL: Table S1 core bits = {_bits_sum:.4f} != I_core = {I_core:.4f}"
)

print(f"  ✓  AUDIT  Table S1 core bits = {_bits_sum:.2f} bits (= I_core = {I_core:.2f})")

# ============================================================
# Render Supplementary Table S1
# ============================================================

print("\n\nSupplementary Table S1 POLR2A (RPB1 core, non-CTD)")
_styled_s1c = (
    _df_s1c.style
    .hide(axis="index")
    .format({"Bits per site log₂(sᵢ)": "{:.2f}"})
    .set_properties(**{
        "font-size": "11pt",
        "font-family": "Calibri, sans-serif",
        "text-align": "center",
        "border": "1px solid #bbb",
    })
    .set_properties(subset=["Residue"], **{"font-weight": "bold"})
    .set_table_styles([
        {"selector": "table", "props": [("border-collapse", "collapse")]},
        {"selector": "th", "props": [("font-size", "11pt"),
                                      ("font-family", "Calibri, sans-serif"),
                                      ("text-align", "center"),
                                      ("border", "1px solid #bbb"),
                                      ("background-color", "transparent")]},
        {"selector": "tbody tr:nth-child(even)", "props": [("background-color", "transparent")]},
        {"selector": "tbody tr:nth-child(odd)",  "props": [("background-color", "transparent")]},
        {"selector": "td", "props": [("background-color", "transparent")]},
    ])
)
display(_styled_s1c)

# %% [code]
## Reproducibility Module 14 - Upper-bound theoretical information capacity encoded by PTMs in subunits Rpb2–Rpb12

# Construction of per-subunit Supplementary Tables S2–S12 for the Rpb2–Rpb12 subunits.
# Quantifies residue-level PTM state models across all non-Rpb1 Pol II subunits, including observed
# PTM combinations and their information contributions.
# I_r212 must match manuscript value, sum of per-site contributions must match I_r212 within numerical tolerance.

# ============================================================
# Imports
# ============================================================
from math import log2, isclose
import pandas as pd
from IPython.display import display

# ============================================================
# Subunit mapping (UniProt → canonical name)
# ============================================================

# UniProt accession -> canonical subunit name
_RPB_NAMES = {
    "P30876": "Rpb2",
    "P19387": "Rpb3",
    "O15514": "Rpb4",
    "P19388": "Rpb5",
    "P61218": "Rpb6",
    "P62487": "Rpb7",
    "P52434": "Rpb8",
    "P36954": "Rpb9",
    "P62875": "Rpb10",
    "P52435": "Rpb11",
    "P53803": "Rpb12",
}

_RPB_ORDER = ["Rpb2","Rpb3","Rpb4","Rpb5","Rpb6",
              "Rpb7","Rpb8","Rpb9","Rpb10","Rpb11","Rpb12"]

# ============================================================
# PTM summary (Rpb2–Rpb12)
# ============================================================

# Unique PTM types in Rpb2-Rpb12
r212_ptm_types = sorted(r212_df["PTM"].unique())
print("Unique PTM types in Rpb2-Rpb12:")
for _t in r212_ptm_types:
    print(f"  - {_t}")

_n_sites_r = r212_df[["Accession", "Residue"]].drop_duplicates().shape[0]
_n_pairs_r = r212_df[["Accession", "Residue", "PTM"]].drop_duplicates().shape[0]
print()
print(f"  Modified residues (unique sites):     {_n_sites_r}")
print(f"  Unique (residue, PTM type) pairs:     {_n_pairs_r}")

# ============================================================
# Per-site state model construction
# ============================================================

# Build per-site data for all subunits
def _r212_res_pos(r):
    try: return int(r[1:])
    except: return 0

_rows_s1r = []
for (_acc, _res), _grp in r212_df.groupby(["Accession", "Residue"]):
    _ptms = sorted(_grp["PTM"].unique())
    _si   = len(_ptms) + 1
    _rows_s1r.append({
        "Residue":                 _res,
        "PTM Type(s)":             " / ".join(_ptms),
        "UniProt accession":       _acc,
        "States (sᵢ)":          _si,
        "Bits per site log₂(sᵢ)": log2(_si),
    })

_df_s1r = (
    pd.DataFrame(_rows_s1r)
    .assign(_pos=lambda _d: _d["Residue"].apply(_r212_res_pos),
            _ord=lambda _d: _d["UniProt accession"].map(_RPB_NAMES)
                             .map({n: i for i, n in enumerate(_RPB_ORDER)}).fillna(99))
    .sort_values(["_ord", "_pos"])
    .drop(columns=["_pos", "_ord"])
    .reset_index(drop=True)
)[["Residue", "PTM Type(s)", "UniProt accession",
   "States (sᵢ)", "Bits per site log₂(sᵢ)"]]

# ============================================================
# Subunit-level aggregation
# ============================================================

_sub_bits = (
    _df_s1r.groupby("UniProt accession")["Bits per site log₂(sᵢ)"]
    .sum()
    .rename(index=_RPB_NAMES)
)
_sub_bits = _sub_bits.reindex([n for n in _RPB_ORDER if n in _sub_bits.index])

# ============================================================
# Output: per-subunit information capacity
# ============================================================

print(f"\nInformation capacity by subunit (Rpb2-Rpb12):\n")
for _name, _b in _sub_bits.items():
    _prefix = "~" if abs(_b - round(_b, 0)) > 0.005 else ""
    print(f"  {_name}: {_prefix}{_b:.2f} bits")

# ============================================================
# Audit: per-subunit information capacity matches expected values from manuscript
# ============================================================

_EXPECTED_PER_SUBUNIT = {
  "Rpb2": 140.36,
  "Rpb3": 33.51,
  "Rpb4": 10.00,
  "Rpb5": 25.34,
  "Rpb6": 5.58,
  "Rpb7": 18.17,
  "Rpb8": 26.58,
  "Rpb9": 15.00,
  "Rpb10": 8.00,
  "Rpb11": 25.34,
  "Rpb12": 3.00
}

print()
for _name, _exp in _EXPECTED_PER_SUBUNIT.items():
    if _exp is None: continue
    _got = float(_sub_bits[_name])
    assert isclose(_got, _exp, abs_tol=0.005), (
        f"AUDIT FAIL: {_name} = {_got:.2f} bits (expected {_exp:.2f})"
    )
    print(f"  ✓  AUDIT  {_name} = {_got:.2f} bits, matches expected value")

print()

# ============================================================
# Global information capacity (Rpb2–Rpb12)
# ============================================================

I_r212 = _I(r212_df)
print(f"\nTotal information capacity of Rpb2-Rpb12 = {I_r212:.2f} bits")

# ============================================================
# Audit:I_r212 must match manuscript value.
#       Sum of per-site information contributions must match I_r212.
# ============================================================

print()
assert isclose(I_r212, 310.89, abs_tol=0.005), (
    f"AUDIT FAIL: I_r212 = {I_r212:.2f} bits  (expected 310.89)"
)
print(f"  ✓  AUDIT  I_r212 = {I_r212:.2f} bits, matches manuscript value")

_bits_sum_r = _df_s1r["Bits per site log₂(sᵢ)"].sum()
assert isclose(_bits_sum_r, I_r212, abs_tol=0.005), (
    f"AUDIT FAIL: Table S1 Rpb2-Rpb12 bit sum = {_bits_sum_r:.4f} != I_r212 = {I_r212:.4f}"
)
print(f"  ✓  AUDIT  Table S1 Rpb2-Rpb12 bits = {_bits_sum_r:.2f} bits, {_bits_sum_r/8:.2f} bytes (= I_r212 = {I_r212:.2f} bits)")

# ============================================================
# Supplementary tables (S2–S12)
# ============================================================

# One styled table per subunit
def _make_styled_r(df_sub):
    return (
        df_sub.style
        .hide(axis="index")
        .format({"Bits per site log₂(sᵢ)": "{:.2f}"})
        .set_properties(**{
            "font-size": "11pt",
            "font-family": "Calibri, sans-serif",
            "text-align": "center",
            "border": "1px solid #bbb",
        })
        .set_properties(subset=["Residue"], **{"font-weight": "bold"})
        .set_table_styles([
            {"selector": "table", "props": [("border-collapse", "collapse")]},
            {"selector": "th", "props": [("font-size", "11pt"),
                                          ("font-family", "Calibri, sans-serif"),
                                          ("text-align", "center"),
                                          ("border", "1px solid #bbb"),
                                          ("background-color", "transparent")]},
            {"selector": "tbody tr:nth-child(even)", "props": [("background-color", "transparent")]},
            {"selector": "tbody tr:nth-child(odd)",  "props": [("background-color", "transparent")]},
            {"selector": "td", "props": [("background-color", "transparent")]},
        ])
    )

for i, _name in enumerate(_RPB_ORDER):
    _acc = {v: k for k, v in _RPB_NAMES.items()}.get(_name)
    if _acc is None:
        continue
    _sub = _df_s1r[_df_s1r["UniProt accession"] == _acc].reset_index(drop=True)
    if _sub.empty:
        continue
    print(f"\nSupplementary Table S{i+2} {_name} ({_acc})")
    display(_make_styled_r(_sub))


# %% [code]
## Reproducibility Module 15 - Upper-bound theoretical PTM information capacity of human RNA polymerase II at the
# molecular and nuclear scales.
# AUDIT: Computed values must equal manuscript values.

# ============================================================
# Imports
# ============================================================
from math import isclose

# ============================================================
# Global information capacity (per molecule)
# ============================================================

I_total = I_CTD + I_core + I_r212

# ============================================================
# Nuclear abundance assumption
# ============================================================

N_POLII = 80_200 # Estimate N_POLII = ~80,200 RNA polymerase II molecules per nucleus
# (BioNumbers ID 112321; Zhao et al., 2014)

# ============================================================
# Nuclear-scale information capacity
# ============================================================

I_nucleus    = I_total * N_POLII
I_nucleus_MB = I_nucleus / (8 * 10**6)

# ============================================================
# Output (per molecule and nucleus)
# ============================================================

print(f"I per molecule (bits):          {I_CTD:.2f} (CTD) + {I_core:.2f} (Rpb1-core) + {I_r212:.2f} (Rpb2-12) = {I_total:.2f} bits")
print(f"I per molecule (bytes):         {I_total / 8:.2f}")
print(f"Pol II molecules per nucleus:  ~{N_POLII:,} (estimate; Zhao et al. 2014)")
print(f"\nUpper-bound capacity per nucleus (bits):        {I_nucleus:.2e}")
print(f"Upper-bound capacity per nucleus (bytes):       {I_nucleus / 8:.2e}")
print(f"Upper-bound capacity per nucleus (Megabytes):  ~{I_nucleus_MB:.2f}")

# ============================================================
# Audits
# ============================================================

print()
assert isclose(I_total, 707.98, abs_tol=0.005), (
    f"AUDIT FAIL: I_total = {I_total:.2f} bits  (expected 707.98)"
)

assert isclose(I_nucleus / (8 * 10**6), 7.10, abs_tol=0.005), (
    f"AUDIT FAIL: I_nucleus = {I_nucleus / (8*10**6):.2f} MB  (expected ~7.10)"
)

print(f"  ✓  AUDIT  I_total = {I_total:.2f} bits\n  ✓  I_nucleus = ~{I_nucleus / (8*10**6):.2f} MB")


# %% [code]
## Reproducibility Module 16 - Pol II PTMs form a multi-timescale biochemical functional architecture
# Literature-derived PTM layers are used to estimate a physiologically
# constrained Pol II information capacity across distinct regulatory
# timescales.

# Layer definitions:
# Fast layer         (~5 s–15 min): phosphorylation
# Intermediate layer (~1–30 min): Pro cis/trans isomerization and O-GlcNAcylation
# Slow layer         (~15 min–24 h): ubiquitination

# I_physio = I_fast + I_intermediate + I_slow

# AUDIT: I_physio and I_phys_nucleus_MB must match manuscript values.

# ============================================================
# Imports
# ============================================================

from math import log2, isclose

# ============================================================
# Layer definitions (literature-derived constants)
# ============================================================

# Fast layer (~5 s to 15 min): phosphorylation - ~30 sites, 5 effective states/site
n_fast, s_fast                 = 30, 5
# Intermediate layer (~1–30 min): Pro isomerization / O-GlcNAcylation - ~32 sites, 2 states/site
n_intermediate, s_intermediate = 32, 2
# Slow layer (~15 min to 24 h): ubiquitination - ~10 sites, 2.5 effective states/site
n_slow, s_slow                 = 10, 2.5

# ============================================================
# Layer-wise information capacity
# ============================================================

I_fast         = n_fast         * log2(s_fast)
I_intermediate = n_intermediate * log2(s_intermediate)
I_slow         = n_slow         * log2(s_slow)
I_physio       = I_fast + I_intermediate + I_slow

S_physio            = 2**I_physio
delta_I             = I_total - I_physio
I_phys_nucleus      = I_physio * N_POLII
I_phys_nucleus_MB   = I_phys_nucleus / (8 * 10**6)

# ============================================================
# Results
# ============================================================

print(f"Fast layer   (~5 s – 15 min):      I = {I_fast:.2f} bits  ({2**I_fast:.2e} states)")
print(f"Intermediate (~1 – 30 min):        I = {I_intermediate:.2f} bits  ({2**I_intermediate:.2e} states)")
print(f"Slow layer   (~15 min – ~24 h):    I = {I_slow:.2f} bits  ({2**I_slow:.2e} states)")
print()
print(f"Physiological total per molecule:  I = {I_physio:.2f} bits  ({S_physio:.2e} states)")
print(f"Theoretical total per molecule:    I = {I_total:.2f} bits")
print(f"Theoretical minus physiological:  ΔI = {delta_I:.2f} bits")
print()
print(f"Physiological capacity per nucleus (bits): {I_phys_nucleus:.2e}")
print(f"Physiological capacity per nucleus (MB):   {I_phys_nucleus_MB:.2f}")

# ============================================================
# Audit
# ============================================================

assert isclose(I_physio, 114.88, abs_tol=0.005), (
    f"AUDIT FAIL: I_physio = {I_physio:.2f} bits  (expected 114.88)"
)
assert isclose(I_phys_nucleus_MB, 1.15, abs_tol=0.005), (
    f"AUDIT FAIL: I_phys_nucleus_MB = {I_phys_nucleus_MB:.2f} MB  (expected 1.15)"
)
print()
print(f"  ✓  AUDIT  I_physio = {I_physio:.2f} bits,  I_phys_nucleus = {I_phys_nucleus_MB:.2f} MB")


# %% [code]
## Reproducibility Module 17 - Total unique PTM sites and modified residues in the integrated Pol II PTM database.

_n_sites_total = all_databases_merge[["Accession", "Residue"]].drop_duplicates().shape[0]
_n_pairs_total = all_databases_merge[["Accession", "Residue", "PTM"]].drop_duplicates().shape[0]
print()
print(f"  Modified residues (unique sites):     {_n_sites_total}")
print(f"  Unique (residue, PTM type) pairs:     {_n_pairs_total}")

# ============================================================
# Audit
# ============================================================

print()
assert _n_sites_total == 631, (
    f"AUDIT FAIL: unique modified residues = {_n_sites_total} (expected 631)"
)
assert _n_pairs_total == 776, (
    f"AUDIT FAIL: unique (residue, PTM) pairs = {_n_pairs_total} (expected 776)"
)
print(f"  ✓  AUDIT  unique sites = {_n_sites_total}, unique (res,PTM) pairs = {_n_pairs_total}")


# %% [code]
## Reproducibility Module 18 - Cellular level PTM information capacity of Pol II (sensitivity analyses).

# Reproduces Supplementary Information S3 (Sections S3.1–S3.5) and
# Supplementary Information S3-Table S1.

# Alternative PTM-state models are evaluated to assess the robustness
# of information-capacity estimates under increasingly conservative
# biological assumptions affecting CTD and non-CTD PTM state spaces.
#
# Sensitivity scenarios:
# S0 = baseline model
# S1 = conservative non-CTD Thr phosphorylation model
# S2 = CTD Pro cis/trans isomerization excluded
# S3 = maximally conservative PTM-state model
#
# AUDIT: all S0–S3 scenario capacities must reproduce manuscript values.

# ============================================================
# Imports
# ============================================================

from math import log10, floor, isclose
import pandas as pd
from IPython.display import display

# ============================================================
# Alternative CTD state models
# ============================================================

# S2: remove CTD cis/trans isomerization; all 51 Pro sites are in the CTD, each binary (1 bit)
I_CTD_no_pro = _I(ctd_df[ctd_df["PTM"] != "cis / trans isomerization"])

# Max-conservative CTD (used in S3):
#   (i)  remove cis/trans isomerization (Pro: 2 -> 1 state)
#   (ii) remove Thr-P                   (Thr: 2 -> 1 state)
#   (iii) remove all Lys mods            (Lys: >=4 -> 1 state)
#   (iv) remove all Arg mods             (Arg: >=2 -> 1 state)

_lys7_labels = {f"K{pos}" for pos, (_, hp) in heptad_position_map.items() if hp == 7}
_ctd_max_df = ctd_df[
    (ctd_df["PTM"] != "cis / trans isomerization") &
    ~(ctd_df["Residue"].str[0].eq("T") & ctd_df["PTM"].str.contains("Phosphorylation", case=False, na=False)) &
    ~ctd_df["Residue"].isin(_lys7_labels) &
    ~ctd_df["Residue"].str[0].eq("R")
]
I_CTD_max = _I(_ctd_max_df)

# ============================================================
# Conservative non-CTD models
# ============================================================

_drop_thr_p = lambda df: df[
    ~(df["Residue"].str[0].eq("T") & df["PTM"].str.contains("Phosphorylation", case=False, na=False))
]
I_core_TC = _I(_drop_thr_p(core_df))
I_r212_TC = _I(_drop_thr_p(r212_df))

# ============================================================
# Scenario overview (Supplementary Information S3)
# ============================================================

print("=" * 110)
print("Supplementary Information S3 -- Sensitivity analyses of Pol II PTM")
print("                               information-capacity estimates")
print("=" * 110)
print(f"Baselines:  I_CTD = {I_CTD:.2f}   I_Rpb1-core = {I_core:.2f}   I_Rpb2-Rpb12 = {I_r212:.2f}   I_PolII = {I_total:.2f} bits")

print()
print("-" * 110)
print("S3.1  Alternative state models tested")
print("-" * 110)
print("(A) CTD Pro cis/trans isomerization:  baseline s=2 (cis/trans); conservative s=1 (excluded as conformational, not covalent)")
print("(B) CTD Arg granularity:    baseline s>=2 (unmod/me...); conservative s=1")
print("(C) CTD Lys granularity:    baseline s>=4 (unmod/Ac/me...); conservative s=1")
print("(D) Thr phosphorylation:    baseline s=2 (unmod/P); conservative s=1")

# ============================================================
# Sensitivity of subcomponents
# ============================================================

print()
print("-" * 110)
print("S3.2  Sensitivity of Rpb2-Rpb12")
print("-" * 110)
print(f"  Baseline                I_(Rpb2-Rpb12) = {I_r212:.2f} bits")
print(f"  Conservative (Thr s=1)  I_(Rpb2-Rpb12) = {I_r212_TC:.2f} bits")

print()
print("-" * 110)
print("S3.3  Sensitivity of Rpb1 core")
print("-" * 110)
print(f"  Baseline                I_(Rpb1-core)  = {I_core:.2f} bits")
print(f"  Conservative (Thr s=1)  I_(Rpb1-core)  = {I_core_TC:.2f} bits")

print()
print("-" * 110)
print("S3.4  Sensitivity - CTD cis/trans Pro isomerization exclusion")
print("-" * 110)
print(f"  Baseline I_(CTD)                          = {I_CTD:.2f} bits")
print(f"  Pro cis/trans excluded I_(CTD)            = {I_CTD_no_pro:.2f} bits")
print(f"  Delta                   ΔI_(CTD)          = {I_CTD - I_CTD_no_pro:.2f} bits  (51 binary Pro sites, all in CTD)")
print(f"  Maximally conservative  I_(CTD)           = {I_CTD_max:.2f} bits")
print("     (i)   Pro cis/trans -> s=1 (unmodified);")
print("     (ii)  Thr-P -> s=1 (unmodified);")
print("     (iii) Lys7 -> s=1 (unmodified);")
print("     (iv)  Arg -> s=1 (unmodified).")

# ============================================================
# Supplementary Information S3-Table S1
# ============================================================

def _sci(x):
    if x == 0: return "0"
    n   = floor(log10(abs(x)))
    sup = str(n).translate(str.maketrans("-0123456789", "⁻⁰¹²³⁴⁵⁶⁷⁸⁹"))
    return f"{x/10**n:.2f} x 10{sup}"

_scenario_I = {}
def _s1_row(lbl, ctd_m, nctd_m, I_ctd, I_c, I_r):
    Ip = I_ctd + I_c + I_r
    _scenario_I[lbl] = Ip
    return [lbl, ctd_m, nctd_m,
            f"{Ip:.2f}",      f"{Ip/8:.2f}",
            _sci(Ip*N_POLII), _sci(Ip*N_POLII/8),
            f"{Ip*N_POLII/8e6:.2f}"]

HDR = ["Scenario",
       "CTD state model",
       "non-CTD state model (Rpb1-core + Rpb2-Rpb12)",
       "I_Pol II (bits per molecule)",
       "Bytes per molecule (I÷8)",
       "I_Cell (bits per nucleus)",
       "Bytes per nucleus (I÷8)",
       "MB per nucleus (10⁶ bytes)"]

ROWS = [
    _s1_row("S0 (baseline)",             "Baseline",           "Baseline",  I_CTD,       I_core,    I_r212),
    _s1_row("S1 (non-CTD conservative)", "Baseline",           "Thr conservative",  I_CTD,       I_core_TC, I_r212_TC),
    _s1_row("S2 (no isomerization)", "Pro cis/trans excluded",    "Baseline",  I_CTD_no_pro, I_core, I_r212),
    _s1_row("S3 (all conservative)",      "Maximally conservative", "Thr conservative",  I_CTD_max,   I_core_TC, I_r212_TC),
]

print()
print("=" * 110)
print("Supplementary Information S3-Table S1. Representative sensitivity scenarios for total Pol II maximal information capacity")
print(f"   (N = {N_POLII:,} Pol II molecules per nucleus)")
print("=" * 110)

df_s1 = pd.DataFrame(ROWS, columns=HDR)
styled_s1 = (
    df_s1.style
    .hide(axis="index")
    .set_properties(**{"font-size": "11pt", "font-family": "Calibri, sans-serif",
                       "text-align": "center", "border": "1px solid #bbb"})
    .set_properties(subset=["Scenario"], **{"font-weight": "bold"})
    .set_table_styles([
        {"selector": "table", "props": [("border-collapse", "collapse")]},
        {"selector": "th",    "props": [("font-size", "11pt"), ("font-family", "Calibri, sans-serif"),
                                         ("text-align", "center"), ("border", "1px solid #bbb")]},
        {"selector": "tbody tr:nth-child(even)", "props": [("background-color", "transparent")]},
        {"selector": "tbody tr:nth-child(odd)",  "props": [("background-color", "transparent")]},
        {"selector": "td", "props": [("background-color", "transparent")]},
    ])
)
display(styled_s1)

# ============================================================
# Audit
# ============================================================

_EXPECTED = {
    "S0 (baseline)":             707.98,
    "S1 (non-CTD conservative)": 644.98,
    "S2 (no isomerization)":     656.98,
    "S3 (all conservative)":     544.44,
}

for _lbl, _exp in _EXPECTED.items():
    _got = _scenario_I[_lbl]
    assert isclose(_got, _exp, abs_tol=0.005), (
        f"AUDIT FAIL: {_lbl} = {_got:.2f} bits  (expected {_exp:.2f})"
    )

print()
for lbl, Ip in _scenario_I.items():
    print(f"  {lbl:<35}  {Ip:.2f} bits/molecule")

print()
print("  ✓  AUDIT  all four S0-S3 scenarios match expected values")
```
